# Bioinformatics Illustrations Decoded by ChatGPT: The Good, The Bad, and The Ugly

**DOI:** 10.1101/2023.10.15.562423

**Authors:** Jinge Wang, Qing Ye, Li Liu, Nancy Lan Guo, Gangqing Hu

## Abstract

Emerging studies underscore the promising capabilities of large language model-based chatbots in conducting fundamental bioinformatics data analyses. The recent feature of accepting image-inputs by ChatGPT motivated us to explore its efficacy in deciphering bioinformatics illustrations. Our evaluation with examples in cancer research, including sequencing data analysis, multimodal network-based drug repositioning, and tumor clonal evolution, revealed that ChatGPT can proficiently explain different plot types and apply biological knowledge to enrich interpretations. However, it struggled to provide accurate interpretations when quantitative analysis of visual elements was involved. Furthermore, while the chatbot can draft figure legends and summarize findings from the figures, stringent proofreading is imperative to ensure the accuracy and reliability of the content.

## Introduction

Known for its remarkable conversational capabilities and extensive knowledge spanning numerous disciplines, large language model chatbots like ChatGPT have gathered significant interests in education^1^, research^2^, and clinical practice^3^. In the field of bioinformatics, ChatGPT serves as an instrumental aid for learning basic bioinformatics^4,5^. Several literatures offer recommendations on harnessing chatbots for more efficient data analysis^4-10^. Further evaluations, centering on biomedical text mining^11,12^, code generation^13,14^, and expertise in genomics/genetics^15-17^, underscore ChatGPT’s potential to facilitate biomedical research. However, present evaluations predominantly focus on text-based inputs. The capability of ChatGPT to interpret alternative inputs, such as bioinformatics illustrations, a task demanding both skills in imaging pattern recognition and domain-specific knowledge, remains to be evaluated.

In this study, we assessed ChatGPT’s ability to interpret bioinformatics illustrations using case studies of data analysis frequently used in cancer research. The initial case addressed differential gene expression analysis, highlighting transcriptional alterations in multiple myeloma when exposed to bone marrow stromal cells^18^. The subsequent study centered on an integrative approach for drug repositioning in non-small cell lung cancer (NSCLC), leveraging network analysis of the transcription factor ZNF71^ref19-21^. The third case embarked on an exploration of the clonal evolution underlying the progression of multiple myeloma using MMRF-CoMMpass data from GDC portal^22^. The final investigation characterized the epigenetic regulatory landscapes surrounding the *YY1* locus in the human B-Lymphoid cancer cell line by harnessing public data from the WashU epigenome browser^23^. We found that the chatbot effectively identified plot types, provided reasonable interpretations, and summarized the findings albeit requiring a careful proofreading. Our in-depth assessment revealed that the chatbot is limited in tasks requiring quantitative analysis, such as estimating numerical values and integrating different visual elements from an illustration to draw conclusions.

## Materials and Methods

### Source of data and procedure of figure generation

Gene expression data used for the “DE gene” case study was sourced from our previous work^18^. This case specifically contrasted the RPMI8226 (a multiple myeloma cell line) in trans-well coculture (T) with bone marrow stromal cells against its monoculture (M). Gene expression values were used to generate a principal component analysis (PCA) plot. Fold change (FC) in expression (T/M; log_2_) and false discovery rate (FDR; -log_10_), which measures the significance of differential expression, were utilized to craft a volcano plot. Additionally, a dot plot illustrating hits in pathway enrichment analysis for up-regulated genes (log_2_FC > 1 and FDR < 0.05) was produced using ShinyGO^24^.

For the “ZNF71” case study, gene expression data with survival information from 194 patients of NSCLC^25^ were used to generate Kaplan-Meier (K-M) curves. We combined gene expression data and PRISM drug screening data of NSCLC cell lines from the DepMap data portal^26^ to relate docetaxel-sensitivity with ZNF71 KRAB expression and to generate Pearson’s correlation plot for examining drug response and CD27 expression. Gene expression of tumors and cell lines were expressed as transcripts per million (TPM). We applied the Boolean implication network algorithm^27^ to construct gene association networks within the context of tumors and NATs using Xu’s LUAD cohort^28^.

For the “Clonal evolution” case study, somatic mutations in a pair of primary and recurrent tumors from an MM patient (ID: 1201) in the MMRF-CoMMpass study were sourced from the GDC data portal^22^. Clonal and subclonal mutations were identified using the MAGOS method^29^. Relative prevalence of (sub)clones and their evolutionary relationships were inferred using the ClonEvol method^30^.

For the “YY1” case study, a screenshot for gene expression, genomic distributions of histone modifications, and chromatin-chromatin interactions in the genomic region encompassing *YY1* in GM12878 was sourced from the WashU Epigenome Browser^23^. Specifically, the ChIP-seq data for histone modifications H3K27me3, H3K4m3, and H3K27ac, along with strand-specific RNA-Seq for gene expression, were loaded from the Encyclopedia of DNA Elements (ENCODE) data hub. The PLAC-seq data for chromatin-chromatin interactions was loaded from the 4DN Nucleome Network. Regions denoting active and inactive chromatin domains, as well as key areas of chromatin-chromatin interactions, were annotated manually.

### Prompts for ChatGPT

We instructed ChatGPT to function as a bioinformatics expert for each case study, introducing the research question associated with each figure at a high level for context. ChatGPT’s interpretive capabilities may be assessed under two steps. In the initial assessments, ChatGPT operated with minimal guidance to provide an overview of an illustration, while in the in-depth assessments, it was guided through a sequence of questions. The questions were prompted to the chatbot one by one, which yielded more detailed responses compared to prompting all questions at once. At the conclusion of each evaluation, ChatGPT was optionally tasked to draft a figure legend and a summary paragraph for the illustration. Prompts used for each case were detailed in **Supplementary File 1**. We repeated each assessment three times using ChatGPT Plus (version dated Sep 25).

## Results

We designed four case studies in cancer research to assess ChatGPT’s proficiency in deciphering bioinformatics illustrations. For each case in the following, we began with a background introduction, presented our interpretations, and then evaluated ChatGPT’s interpretations. Our key findings for each case study were documented below, with additional comments added to Supplementary Files 2 to 19, which are screenshots of our chat history with ChatGPT.

### The “DE gene” case study: gene expression analysis for Multiple Myeloma induced by bone marrow stromal cells

Multiple myeloma (MM) is a hematologic malignancy predominantly proliferating within the bone marrow (BM). The BM stromal cells are known to support tumor growth and promote drug resistance^31^. RNA-Seq gene expression data from our early work^18^ compares MM cells in trans-well coculture with BM stromal cells to those in monoculture. Results from PCA analysis in **Fig. 1** echoed the findings from our earlier work^18^, showing substantial transcriptomic changes in MM cells induced by BM stromal cells (**Fig. 1A**). The upregulation in the expression of *SOCS3* and *JUNB* acts as established positive controls drawn from the literature (**Fig. 1B**). KEGG pathway enrichment analysis pinpointed the JAK-STAT signaling pathway among the top hits for upregulated genes (**Fig. 1C**). This pathway is typically activated by IL-6 secreted from stromal cells when cocultured with MM cells^32^. The observation supports that BM stromal cells induce substantial gene expression changes in MM cells, including the expression upregulation of key signaling pathways such as JAK-STAT relevant regulatory genes such as *SOCS3* and *JUNB*^18^.

**Fig. 1:**
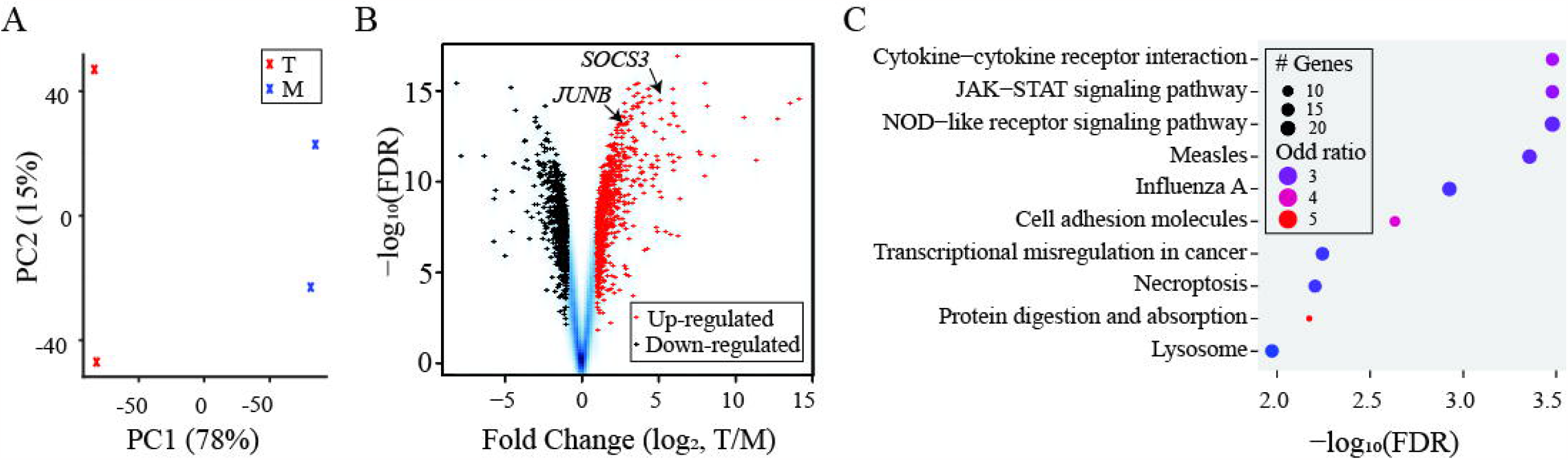
The “DE gene” case study. (**A**) PCA analysis showing two groups of samples: “T” in red and “M” in blue. (**B**) A volcano plot for FCs and FDRs from differentially expressed gene analysis with up-regulated genes in red and down-regulated genes in black. *SOCS3* and *JUNB* indicated by arrowheads. (**C**) Dot plot summarizing the top ten hits from KEGG pathway enrichment analysis applied to the up-regulated genes.

We initially prompted ChatGPT to offer an overview of each panel, suggest enhancements for data presentation, draft a figure legend, and summarize the findings (**Supplementary Files 2 to 4**). Across all three evaluations, ChatGPT accurately identified the plot types for **Fig. 1A** (PCA plot) and for **Fig. 1B** (volcano plot). For **Fig. 1C** although not explicitly mentioning the plot type, it correctly detailed each component of the plot. Interestingly, all tests failed to interpret color-coding between groups in **Fig. 1B**. It mentioned the two indicated genes in **Fig. 1B**, *SOCS3* and *JUNB*, but did not delve deeper into their significance. Drafts on summary paragraph were satisfactory, albeit they could be improved by adding details. Refinements suggested by ChatGPT for data presentation aligned with standard practices.

In the in-depth assessment, ChatGPT was challenged to estimate specific numbers for indicated genes from the figure and align findings with current knowledge to interpret the figure (**Supplementary Files 5 to 7**). Impressively, it satisfactorily estimated the log_2_FC for *SOCS3* from **Fig. 1B** but over-estimated the numbers for *JUNB*. It failed to decode colors for sample or gene groups in **Fig. 1A&1B**. Throughout the evaluations, ChatGPT correctly recalled SOCS3 as a suppressor of JAK-STAT signaling pathway and JUNB as a proto-oncogene in the context MM biology. It mentioned the activation of the JAK-STAT signaling pathway by BM stromal cell and utilized this knowledge to deduce that up-regulated genes were used for the KEGG enrichment analysis. The summary paragraphs uniformly added the details on expression up-regulation of *SOCS3* and *JUNB*. However, imperfections from response to previous inquiries such as misinterpretations of color-coding were repeated in the legends and/or summaries.

### The “ZNF71” case study: Drug-reposition inferred through ZNF71 in Non-Small Cell Lung Cancer

Zinc finger protein 71 (ZNF71) is a pivotal member of a seven-gene assay we previously developed to predict chemotherapy efficacy for NSCLC^19^. As part of the expansive KRAB zinc finger transcription factors family, termed KRAB-ZNFs, ZNF71 possesses the distinctive KRAB domain and functions as a transcriptional repressor. **Fig. 2** summarizes key findings from our previous for this TF in NSCLC: 1) ZNF71 KRAB’s expression correlates with a diminished overall survival rate in NSCLC patients^20^ (**Fig. 2A**); 2) Docetaxel-resistant NSCLC cell lines exhibit elevated ZNF71 KRAB expression levels in comparison to their chemotherapy-responsive counterparts^20^ (**Fig. 2B**); 3) ZNF71 is centered in a comprehensive association network with genes crucial for the intracellular innate immune response (IIIR) and suppresses their expression^21^ (**Fig. 2C**); and 4) Our gene association networks identified PQ-401 as a drugs repositioning candidate for NSCLC treatment: Its Ln(EC_50_) in NSCLC cell lines negatively correlates with CD27 mRNA expression (**Fig. 2D)**, suggesting potential immune-enhancing effects induced by the drug in NSCLC therapy^21^.

**Fig. 2:**
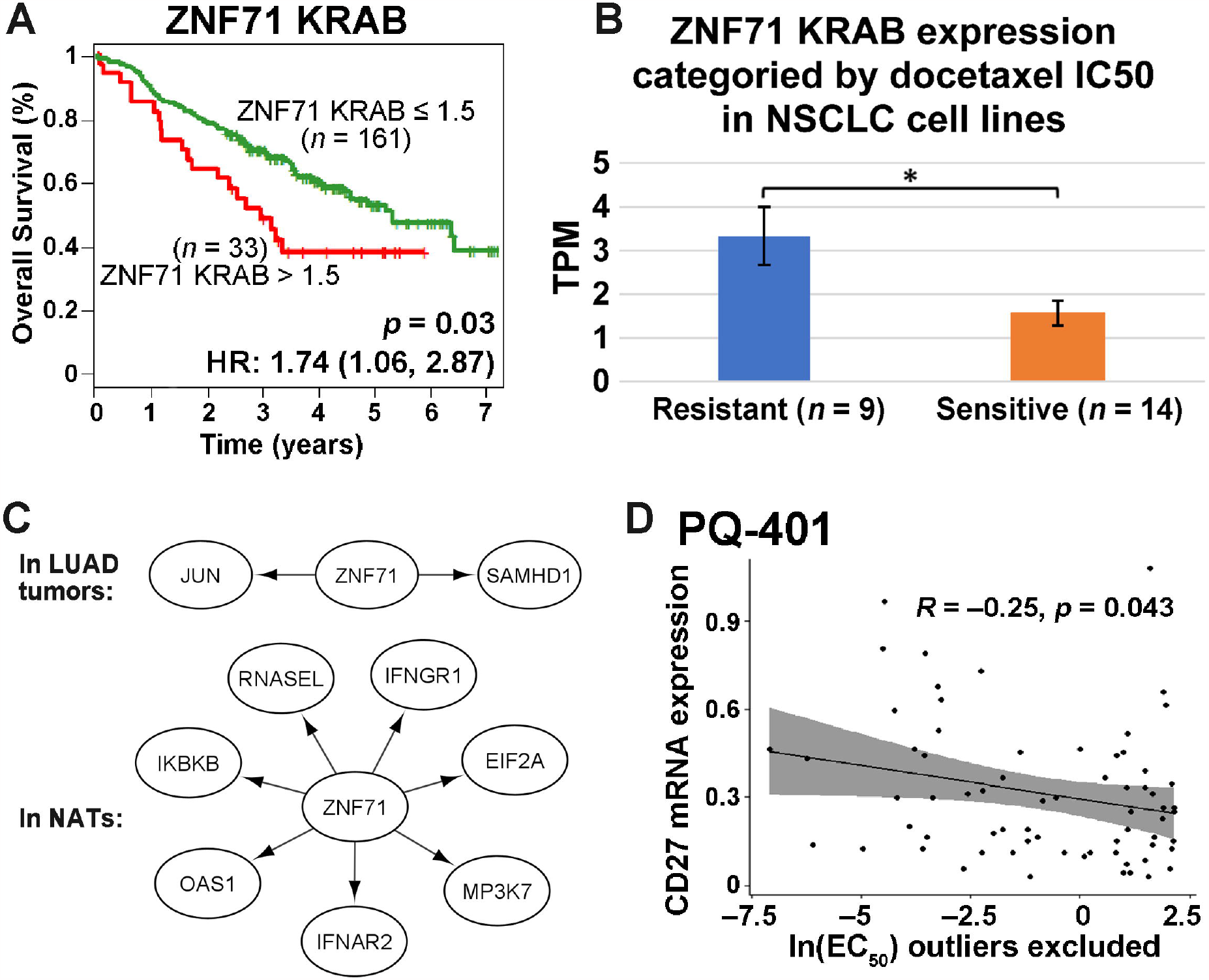
The “ZNF71” case study. (**A**) Kaplan–Meier survival plot for NSCLC patients (GSE81089) sorted by ZNF71 KRAB isoform expression. Red: patients with a higher ZNF71 KRAB expression (TPM > 1.5); Green: othes. (**B**) A bar chart for the expression of ZNF71 KRAB isoform among NSCLC cell lines sorted by docetaxel sensitivity. (**C**) Gene association network of ZNF71 and IIIR genes in LUAD tumors and NATs. (**D**) Scatter plot for CD27 mRNA expression level and drug response to PQ-401 (EC_50_) among NSCLS cell lines.

In our evaluation, we engaged ChatGPT to provide an overview of **Fig. 2** and then address specific queries related to each panel (**Supplementary Files 8 to 10**). The overview accurately captured the plot type from each panel. Upon further probing, ChatGPT reliably pinpointed ZNF71 KRAB as an unfavorable prognosis marker and substantialized its findings by hazard ratio extracted from the Kaplan-Meier plot (**Fig. 2A**). However, it failed to discern the color representing different patient groups on the plot. For **Fig. 2B**, we tasked ChatGPT to estimate the ZNF71 KRAB expression value from a bar plot, contrasting two cell line groups categorized by their sensitivity to docetaxel. The chatbot estimated the average expression as 4-4.5 for the resistance group, while the reference was 3.3. In **Fig. 2C**, ChatGPT was assessed based on its ability to discern connected genes from a gene association network. It successfully identified genes interacting with ZNF71 in all tests but incorrectly spelled the gene name *IKBKB*, with similar incidences observed for *CD27* in **Fig. 2D**. Lastly, we asked ChatGPT to extract the Pearson correlation coefficient (R) in **Fig. 2D**. The bot adeptly determined the R value (−0.25) and deduced its negative correlation between CD27 expression and EC_50_ of PQ-401 (**Supplementary Files 8 to 10**). However, in one instance, it incorrectly inferred that a higher CD27 expression implied more resistance to PQ-401 (**Supplementary File 10**).

### The “Clonal Evolution” case study: Inference of clonal evolution inference in Multiple Myeloma

A tumor is composed of clonal and subclonal cell populations. Understanding the evolutionary relationships among these cell populations is crucial for deciphering tumor development and progression. Different (sub)clones acquire unique somatic mutations that often exhibit varied variant allele frequencies (VAFs), which can be used to infer their cellular prevalence within a tumor. The “clonal evolution” case study illustrated somatic mutations in a pair of primary and recurrent MM tumor samples. Our observations from **Fig. 3** included: 1) The scatterplot contrasting the VAFs of somatic mutations between the primary and recurrent MM tumors shows four distinct clusters, whereas high-impact mutations predominantly residing in the founding clone, denoted as cluster 1 (**Fig. 3A**); 2) The connected line plot illustrates the mean VAF of clusters 3 and 4 increases in the recurrent tumors as compared to the primary tumor, while the mean VAF of cluster 2 decreases and cluster 1 remains largely unchanged (**Fig. 3B**); 3) The bell plots, also known as simplified fish plots, provide a schematic representation of the tumor’s evolutionary dynamics over time – new subclones emerge and substitute the founding clone and old subclones (**Fig. 3C&3D**); 4) Spheres of cells illustrate cellular composition of the tumors at the time of sample taken – the founding clone disappears from both the primary and recurrent tumors and the dominant subclone in the primary tumor represented by cluster 2 disappears from the recurrent tumor (**Fig. 3E&3F**); and 5) A branch-based tree illustrating the relational and seeding patterns shows the founding clone evolves into two separate subclones (cluster 2 and 4) and cluster 2 further gives rise to cluster 4 (**Fig. 3G**). These observations suggest that high-impact functional mutations facilitated the growth of the founding clone. During the development of the primary tumor, emergent mutations gave rise to a subclone (cluster 2), which eventually surpassed the founding clone in cellular dominance. Subsequent mutations led to the evolution of two additional (sub)clones (clusters 3 and 4 albeit with low abundance), originating from the founding and dominant (sub)clones, respectively. Upon the tumor’s recurrence, subclone 4 underwent expansion, becoming the predominant population, while subclone 2 experienced a reduction in prevalence.

**Fig. 3:**
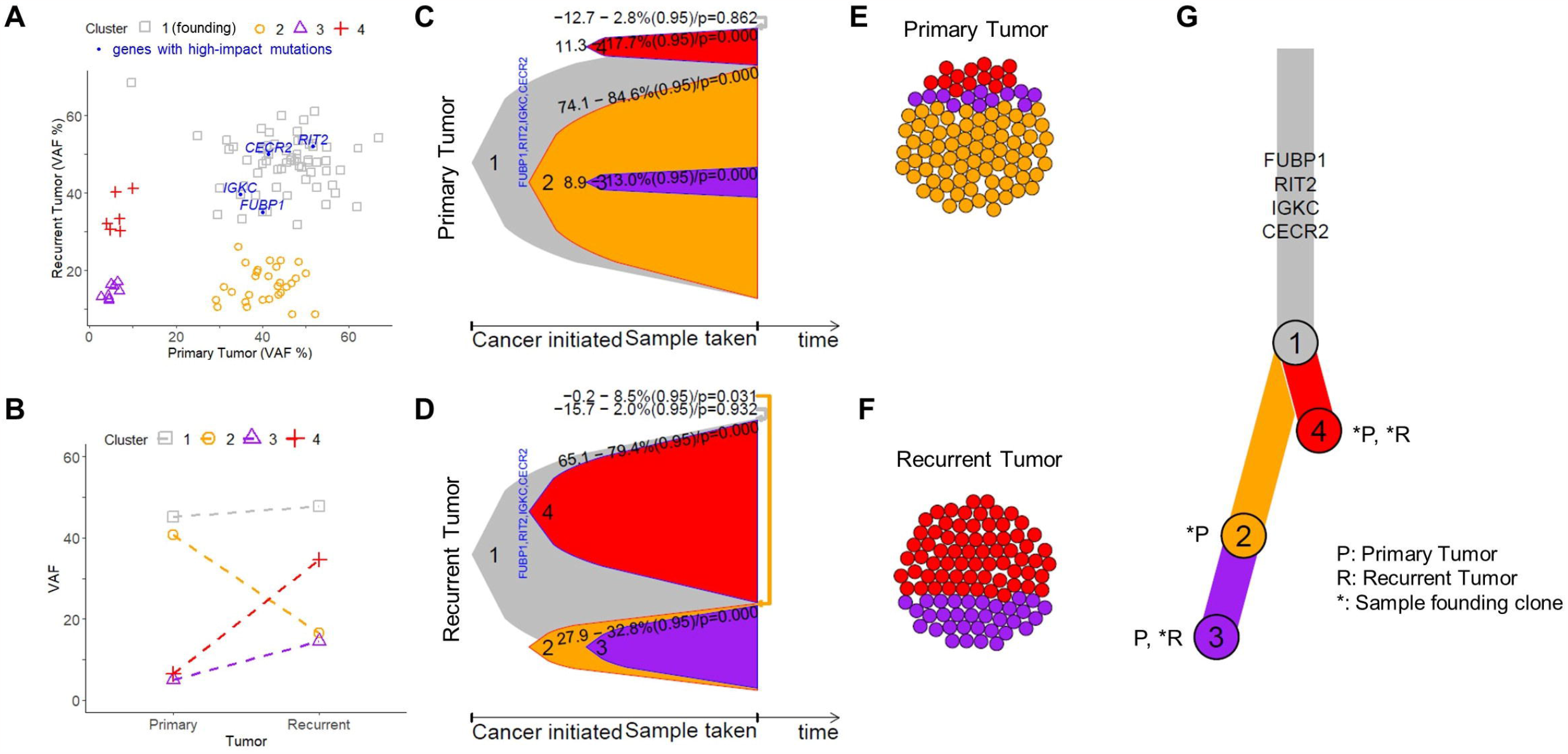
The “Clonal evolution” case study. (**A**) Scatter plot for VAF% contrasting recurrent tumor (y-axis) to primary tumor (x-axis). Each dot is a gene and genes with significant impact are indicated. Primary and recurrent tumors sourced from the same patient. (**B**) Connected dot plot illustrating the changes of mean VAF of each cluster from the primary tumor to the recurrent. (**C**) Bell plot illustrating the relative prevalence of each (sub)clone as a function of cancer progression for primary tumor. Each “bell” symbolizes a (sub)clone, with the area of the bell being proportional to cellular prevalence within the tumor. A 95% confidence interval, along with a *p*-value testing for non-zero prevalence, is displayed for each (sub)clone. (**D**) Similar to panel **C** but for the recurrent tumor. (**E**) Circle packing plots illustrating cellular composition of the primary tumor, exampled with a total of 100 cells. (**F**) Similar to panel **E** but for the recurrent tumor. (**G**) A branch-based tree illustrating the relational and seeding patterns of the (sub)clones between the primary and recurrent samples. Indicated are genes facilitating the growth of the founding clone.

When inspecting its overview on figure panels and the summary paragraph, we found that ChatGPT demonstrated a commendable ability to interpret the figure (**Supplementary Files 11 to 13**). It accurately identified the correct types of plots, except for **Fig. 3C&3D**, where bell plots were misidentified as area plots. We further made in-depth inquiries aimed at assessing ChatGPT’s quantitative analysis capability, involving tasks like color decoding, sorting clusters by sizes, identifying VAF-changes in clusters, and recognizing leaf nodes from a clonal evolutionary tree. For these tasks, the chatbot encountered difficulties in connecting colors to designated clusters and at times referred to non-existing colors. Similarly, it failed to accurately sort clusters by size for **Fig. 3A** and often referenced non-existent clusters for **Fig. 3E&3F** (**Supplementary Files 11 to 13**). Intriguingly, ChatGPT sometimes over-interpreted figures. For instance, it erroneously suggested that node sizes in **Fig. 3G** were proportional to cellular prevalence, despite all nodes being of equal size (**Supplementary Files 11**). The chatbot only sporadically spelled cluster names in the summary paragraphs and could benefit from adding details to improve the generally sound content. In conclusion, although ChatGPT showcased a decent interpretation of cancer clonal evolution from the figure, a careful review of the legend and summary is advised, especially when interpreting color-coded elements and when quantitative analysis is required, even at a basic level.

### The “YY1” case study: Epigenetic regulation of YY1 in a human B-Lymphoid cancer line

Epigenetic regulation of gene expression operates through mechanisms such as post-translational modifications of histone proteins, DNA methylation, and chromatin-chromatin interactions. Active transcriptional promoters are typically marked by H3K4me3, while transcriptionally silent promoters are characterized by H3K27me3^ref33^. Active enhancer regions, on the other hand, are often marked by H3K27ac^34^. The 3D structure of chromatin also plays a crucial role in transcriptional regulation through promoter-promoter and promoter-enhancer interactions^35^. In this case study, we explored the epigenetic landscapes surrounding the *YY1* locus in GM12878, a human B-Lymphoid cancer cell line prioritized by ENCODE for comprehensive data generation. Our observations of **Fig. 4** are: 1) The *YY1* locus is transcriptionally active, evident from its promoter regions enriched with H3K4me3 and H3K27ac but devoid of H3K27me3 (as indicated in the “A2” region of **Fig. 4**); 2) Adjacent genes, namely *DEGS2, SLC25A29*, and *SLC25A47*, appear transcriptionally inactive, marked by promoters enriched with H3K27me3 (I1 and I2 regions). Conversely, *EVL* and *WARS1* genes display transcriptional activity, resembling the histone modification pattern of *YY1* (A1 and A3 regions); 3) Four presumptive enhancer regions characterized by H3K27ac are situated: three between *DEGS2* and *YY1* (E1, E2, and E3 regions) and one immediately downstream of *YY1* (E4 region); 4) PLAC-seq 3D chromatin interaction data signifies *YY1* promoter interactions with the three upstream enhancers (indicated as green circles on the heatmap in **Fig. 4**). Additionally, the *YY1* promoter appears to interact with the promoters of *EVL* (left circle), *DEGS2* (middle circle), and *WARS1* (right circle). The observations identified *YY1* as a center element of a local chromatin-chromatin interaction hub, which included three upstream enhancers, three nearby genes with different transcriptional status, and the promoter of *YY1* itself.

**Fig. 4:**
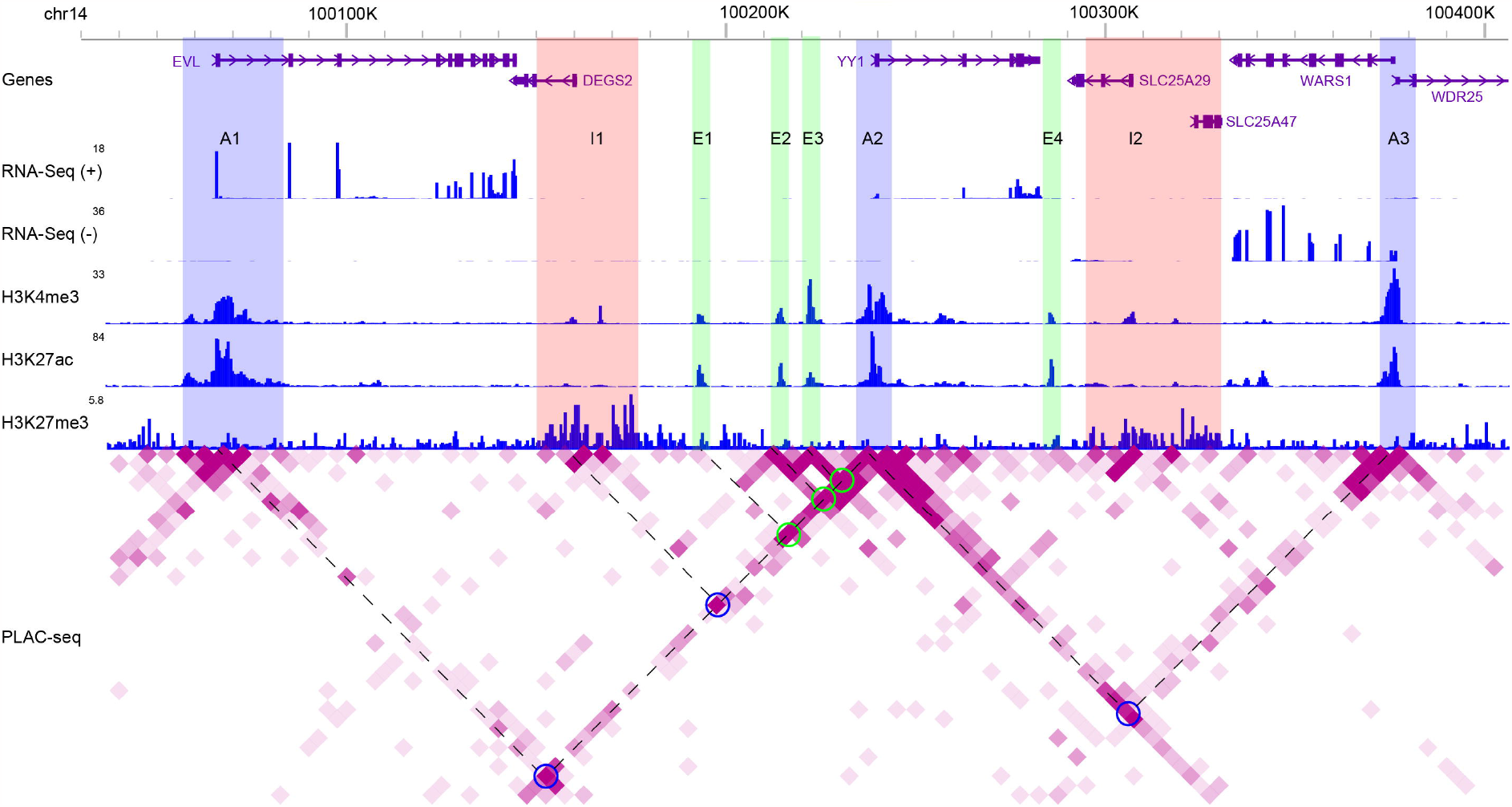
The “YY1” case study. WashU epigenome browser tracks illustrating distributions of strand-specific RNA-Seq reads, ChIP-seq reads for H3K4me3, H3K27ac, and H3K27me3, as well as PLAC-seq reads for chromatin-chromatin interactions. Blue rectangles (A1 to A3): active promoter regions; Pink rectangles (I1 and I2): inactive promoter regions; Green rectangles (E1 to E4): active enhancer regions; Blue circles: promoter regions (A1, I1, and A3) interacting with YY1; Green circles: enhancer regions (E1 to E3) interacting with YY1. Dashed lines shared with one circle on the heatmap: two interacting genome regions. Degree of purpleness on the heatmap indicting interaction strength.

In our preliminary assessment, we directed ChatGPT to decipher the transcriptional regulation of the *YY1* gene. Without specific guidance, ChatGPT mainly focused on the *YY1* locus (**Supplementary Files 14 to 16**). It consistently identified *YY1* as transcriptionally active, basing this on the RNA-Seq signals and associated histone modification patterns. While the chatbot’s interpretation of the promoter-enhancer interactions for *YY1* was generally correct, it was generic and lacked detail. Notably, ChatGPT offered valuable suggestions to enhance data presentation, such as adopting a colorblind-friendly palette, emphasizing the gene of focus (*YY1* in this case), and introducing legends for distinct symbols. The figure legends crafted by the chatbot were concise and captured key elements but ignored the highlighted regulatory domains. While two tests explained the circles on the heatmap (**Fig. 4**), one mistakenly interpreted them as indicating interaction strength rather than hotspots (**Supplementary File 16**). The summary paragraphs of the findings from the illustration, although scientifically accurate, remained at surface-level.

We continued an in-depth evaluation of ChatGPT’s ability to analyze intricate details (**Supplementary Files 17 to 19**), beginning with a task to identify expressed genes via RNA-Seq signals and associated histone modifications. While the chatbot accurately identified *YY1, EVL*, and *WARS* as expressed, it frequently misinterpreted the expression and histone modification patterns of other genes (**Table 1**). Subsequent tasks involved the interpretation of chromatin domains A1-A3 (active promoters), I1 and I2 (inactive promoters), and E1-E4 (active enhancers) (**Fig. 4**). ChatGPT correctly counted the domains in each category and, utilizing histone modification patterns effectively recognized ‘A’ regions as participating in active transcriptional regulation. However, it stumbled in identifying their positional relationships to genes, occasionally misstating them in intragenic regions. While ‘I’ regions were generally classified correctly as transcriptionally repressive due to the presence of H3K27me3 (**Supplementary Files 17 & 19**), one instance failed to identify this marker, leading to a misleading interpretation (**Supplementary File 18**). Lastly, for the ‘E’ regions, ChatGPT rightly identified them as enhancers, noting their robust H3K27ac and relatively weak H3K4me3, and deduced their role in transcriptional regulation of target genes via chromatin looping. These outcomes suggest that while ChatGPT can leverage existing molecular biology knowledge to interpret results, it struggled to infer positional relationships among visual elements.

**Table 1:** Summary of ChatGPT’s identification of expressed genes and the associated histone modifications.

|  |  | *EVL* | *DEGS2* | *YY1* | *SLC25A29* | *SLC25A47* | *WARS1* | *WDR25* |
| --- | --- | --- | --- | --- | --- | --- | --- | --- |
| RNA-Seq signal | Manual | + | - | + | - | - | + | - |
|  | Eval 1 | + | + | + | + | + | + | - |
|  | Eval 2 | + | + | + | + | + | + | + |
|  | Eval 3 | + | - | + | - | - | + | - |
| H3K4me3 | Manual | + | + | + | + | - | + | + |
|  | Eval 1 | + | + | + | + | + | + | - |
|  | Eval 2 | + | + | + | na | na | + | na |
|  | Eval 3 | + | na | + | na | na | + | na |
| H3K27ac | Manual | + | - | + | - | - | + | + |
|  | Eval 1 | + | + | + | + | + | + | - |
|  | Eval 2 | + | + | + | + | + | + | + |
|  | Eval 3 | + | na | + | na | na | + | + |
| H3K27me3 | Manual | - | + | - | + | + | - | - |
|  | Eval 1 | - | - | - | - | - | na | + |
|  | Eval 2 | na | na | na | na | na | na | na |
|  | Eval 3 | - | + | na | + | + | na | + |

“+”: expressed or presence of the indicated histone marker; “-”: not expressed or absence of the indicated histone marker; “na”: not explicitly identified by ChatGPT; “Manual”: manual annotation; “Eval 1/2/3”: the three times of assessments; Red rectangle: ChatGPT's identification conflicting with manual annotation.

In our next assessment, we aimed to interpret the chromatin interactions between the *YY1* promoter and three enhancers (denoted by green circles) as well as with three promoter regions (denoted by blue circles; **Fig. 4**). Each circle signifies the interaction between two genomic regions, linked by dashed lines to the circle. For instance, chromatin domains A1 and A2 connected to the leftmost blue circle, denoting their physical proximity. ChatGPT could not accurately discern the colors of the two types of circles and reported non-existent purple circles. While manually counting the circles is a straightforward task for human analysts, ChatGPT consistently failed by inaccurately counting the circles in all three evaluations (**Supplementary Files 17 to 19**). When challenged to identify interacted chromatin regions indicated by circles, none of the reports from ChatGPT was correct. This underscored ChatGPT’s limitations in not only counting simple visual elements like circles but also deducing relationships between connected visual elements.

Figure legends and summaries from the initial evaluation were largely good, though they occasionally overlooked explanations for custom markings on plots and lacked sufficient details for concrete conclusions (**Supplementary Files 14 to 16**). During the in-depth assessment with specific questions, ChatGPT incorporated its responses into the figure legends and result summaries to enrich depth. This approach, however, sometimes led to the inclusion of errors or misleading information from the previous responses (**Supplementary Files 14 to 16**), emphasizing the need for rigorous human review to guarantee accuracy.

## Discussion

Data visualization is crucial in conveying results from bioinformatics analyses. Large language model chatbots such as ChatGPT have demonstrated an ability to transform natural language prompts into relevant visual representations through coding^36,37^. The newly introduced feature of ChatGPT to take image inputs offers a promising avenue for identifying patterns within figures, offering interpretations, summarizing findings, and beyond^38^. However, interpreting bioinformatics illustrations demands specific domain knowledge, an area where the chatbot might not be thoroughly trained. Additionally, chatbots exhibit tendencies toward ‘hallucinations’ when navigating tasks outside their training scope. Considering these factors, a systematic assessment becomes urgent to discern the strengths, weaknesses, and potential pitfalls of utilizing chatbots for interpreting scientific illustrations.

We carefully designed four use cases to assess various aspects of ChatGPT’s capability in interpreting bioinformatics illustrations. These case studies encompassed a diverse type of plots, including scatter plot, bar plot, box plot, dot plot, PCA plot, volcano plot, KM survival plot, interaction network, bell plot, circle-packing plot, and a multi-track genome browser image. Notably, ChatGPT adeptly recognized these different plot types (except for bell plot) and elucidated key elements within the plot effectively. On the other hand, another chatbot, Bard, often struggled to discern the plot type (data not shown). Hence, we focused our assessments on ChatGPT.

In bioinformatics illustrations, manual adjustments are frequently made to enhance content presentation. This is the case in several plots of our design: the highlighted genes in the volcano plot, the patient group annotations in the Kaplan-Meier plot, and the emphasized genomic regions and chromatin interaction hotspots in the WashU genome browser image. Interpreting these manually edited elements demands quantitative analysis, particularly by assessing their positional relationships to other elements. ChatGPT struggled in this challenge: It could not accurately determine the coordinates for the indicated genes (*JUNB* and *SOCS3*) in the volcano plot nor could it correctly associate patient group annotations with their respective color-coded survival curves in the KM plot. Its attempt to identify genomic regions linked to interaction hotspots was largely unsuccessful. These findings underscored the need for further refinement of ChatGPT in interpreting nuanced, manually edited elements in bioinformatics illustrations.

Correctly explaining bioinformatics illustrations requires domain-specific knowledge. Our testing indicated that ChatGPT taps into existing biological knowledge to interpret results. Even in initial evaluations without detailed instructions in the “YY1” case study, the chatbot referenced the active H3K4me3 and repressive H3K27me3 histone modifications to elucidate the transcriptional status of *YY1*; it further referred H3K27ac-decorated regions as enhancers and combined with chromatin interaction data to support their transcriptionally regulatory role of *YY1*, though without specifying the details. In the “DE gene case”, when prompted to combine literature for interpretation, the chatbot cited the canonical negative feedback loop between SOCS3 and cytokine signaling to explain *SOCS3*’s transcriptional activation. It then deduced that up-regulated genes were used for pathway enrichment analysis and were linked to the JAK-STAT signaling pathway, citing it typically triggered by cytokines. However, ChatGPT often will need to be explicitly prompted to cross-refer with existing knowledge; otherwise, its feedback remains predominantly centered on the direct content of the illustrations.

We further evaluated ChatGPT’s proficiency at summarizing illustrations by prompting it to craft figure legends and summative paragraphs. While its figure legends were largely satisfactory, minor issues were presented, such as inaccuracies in listing replicate numbers, misidentification of colors in the legend, and omissions of plot markers. ChatGPT’s summary paragraphs failed in short of details. During the in-depth assessments, while the chatbot did include details, it occasionally reflected errors made in early responses. This was particularly evident in the “YY1” case study: the summary cited marked regulatory regions from the plot but mischaracterized their interaction relationship with the *YY1* promoter or provided misleading interpretations for some of those regions. Thus, while ChatGPT is equipped to draft figure legends and summaries, rigorous manual proofreading and detailed revision are indispensable to ensure accuracy and prevent dissemination of misleading information.

The present study is not without limitations. Our evaluations are based on four use cases, with further extrapolation to other topics necessitating collaboration with experts in relevant domains. The initial evaluations, however, do illuminate consistent themes regarding ChatGPT’s performance in interpreting bioinformatics illustrations: While the chatbot demonstrates proficiency with standardized plots, it struggles with customized elements requiring visual quantification. Additionally, while it can draft figure legends and offer summaries, human proofreading is paramount to ensure accuracy and depth in interpretations. Nevertheless, we were impressed by the chatbot’s ability to leverage existing knowledge in making interpretations. Future endeavors should investigate the potential benefits of using prompt engineering techniques^38^, like one/few-shot learning, visual chain-of-thought^39^ and PanelGPT^40^, to enhance the chatbot’s interpretative capabilities. Lastly, we haven’t assessed ChatGPT’s ability to draft discussion in conjunction with references, due to the inability to simultaneously invoke image reading and BING access in the current version.

## Conclusions

In conclusion, our assessments revealed that ChatGPT exhibited significant promise in deciphering bioinformatics illustrations. Nevertheless, it faces challenges, especially in interpreting customized visual elements and conducting quantitative analyses. To harness ChatGPT’s full potential in this direction, human oversight is indispensable for validation and refinement of its outputs. Our analysis also underscored the needs of comparable evaluations when extending the image-reading capability of the chatbot to other critical domains such as medical diagnosis. As we progress, infusing chatbots with domain-specific expertise and image-input-specific prompts will be pivotal in enhancing the quality of their responses.

## Data availability

Prompts and scripts to support the conclusions are in the manuscript.

## Author Contributions

Conceptualization: G.H.; Formal analysis: J.W., Q.Y., and L.L.; Writing - Original Draft: Q.Y., L.L., and G.H. Writing - Review & Editing: all authors. Supervision: N.L.G. and G.H.

## Acknowledgements

NIH-NIGMS grants P20 GM103434, U54 GM-104942, and 1P20 GM121322, as well as NSF 2125872 (GH). NIH-NLM grant No. R01LM013438 to LL. NSF 2234456 to NG. The content is solely the responsibility of the authors and does not necessarily represent the official views of the NIH and NSF. The writing was polished by ChatGPT.

## Supplementary Files

**Supplementary File 1**: Prompts used to guide ChatGPT to interpret bioinformatics illustrations.

**Supplementary File 2**: Screen shot of chat session for the “DE gene” case study (initial assessment 1).

**Supplementary File 3**: Screen shot of chat session for the “DE gene” case study (initial assessment 2).

**Supplementary File 4**: Screen shot of chat session for the “DE gene” case study (initial assessment 3).

**Supplementary File 5**: Screen shot of chat session for the “DE gene” case study (in-depth assessment 1).

**Supplementary File 6**: Screen shot of chat session for the “DE gene” case study (in-depth assessment 2).

**Supplementary File 7**: Screen shot of chat session for the “DE gene” case study (in-depth assessment 3).

**Supplementary File 8**: Screen shot of chat session for the “ZNF71” case study (assessment 1).

**Supplementary File 9**: Screen shot of chat session for the “ZNF71” case study (assessment 2).

**Supplementary File 10**: Screen shot of chat session for the “ZNF71” case study (assessment 3).

**Supplementary File 11**: Screen shot of chat session for the “Clonal evolution” case study (assessment 1).

**Supplementary File 12**: Screen shot of chat session for the “Clonal evolution” case study (assessment 2).

**Supplementary File 13**: Screen shot of chat session for the “Clonal evolution” case study (assessment 3).

**Supplementary File 14**: Screen shot of chat session for the “YY1” case study (initial assessment 1).

**Supplementary File 15**: Screen shot of chat session for the “YY1” case study (initial assessment 2).

**Supplementary File 16**: Screen shot of chat session for the “YY1” case study (initial assessment 3).

**Supplementary File 17**: Screen shot of chat session for the “YY1” case study (in-depth assessment 1).

**Supplementary File 18**: Screen shot of chat session for the “YY1” case study (in-depth assessment 2).

**Supplementary File 19**: Screen shot of chat session for the “YY1” case study (in-depth assessment 3).

## References

1 Milano, S., McGrane, J. A. & Leonelli, S. Large language models challenge the future of higher education. Nat Mach Intell 5, 333–334 (2023). 10.1038/s42256-023-00644-2

2 van Dis, E. A. M., Bollen, J., Zuidema, W., van Rooij, R. & Bockting, C. L. ChatGPT: five priorities for research. Nature 614, 224–226 (2023). 10.1038/d41586-023-00288-7

3 Lee, P., Bubeck, S. & Petro, J. Benefits, Limits, and Risks of GPT-4 as an AI Chatbot for Medicine. New Engl J Med 388, 1233–1239 (2023). 10.1056/NEJMsr2214184

4 Shue, E., Liu, L., Li, B., Feng, Z., Li, X. & Hu, G. Empowering Beginners in Bioinformatics with ChatGPT. Quantitative Biology 11, 105–108 (2023). 10.15302/J-QB-023-0327

5 Piccolo, S. R., Denny, P., Luxton-Reilly, A., Payne, S. H. & Ridge, P. G. Evaluating a large language model’s ability to solve programming exercises from an introductory bioinformatics course. PLoS Comput Biol 19, e1011511 (2023). 10.1371/journal.pcbi.1011511

6 Merow, C., Serra-Diaz, J. M., Enquist, B. J. & Wilson, A. M. AI chatbots can boost scientific coding. Nat Ecol Evol (2023). 10.1038/s41559-023-02063-3

7 Perkel, J. M. Six tips for better coding with ChatGPT. Nature 618, 422–423 (2023). 10.1038/d41586-023-01833-0

8 Lubiana, T. et al. Ten quick tips for harnessing the power of ChatGPT in computational biology. PLoS Comput Biol 19, e1011319 (2023). 10.1371/journal.pcbi.1011319

9 Rahman, C. R. & Wong, L. How much can ChatGPT really help Computational Biologists in Programming? arXiv (2023). 10.48550/arXiv.2309.09126

10 Pells, R. Spice up your bioinformatics skill set with AI. Nature 622, S1–S3 (2023).

11 Chen, Q. et al. An Extensive Benchmark Study on Biomedical Text Generation and Mining with ChatGPT. Bioinformatics (2023). 10.1093/bioinformatics/btad557

12 Jin, Q., Yang, Y., Chen, Q. & Lu, Z. GeneGPT: Augmenting Large Language Models with Domain Tools for Improved Access to Biomedical Information. ArXiv (2023).

13 Tang, X., Qian, B., Gao, R., Chen, J., Chen, X. & Gerstein, M. BioCoder: A Benchmark for Bioinformatics Code Generation with Contextual Pragmatic Knowledge. arXiv (2023). 10.48550/arXiv.2308.16458

14 Sobania, D., Briesch, M., Hanna, C. & Petke, J. An Analysis of the Automatic Bug Fixing Performance of ChatGPT. 2023 Ieee/Acm International Workshop on Automated Program Repair, Apr, 23–30 (2023). 10.1109/Apr59189.2023.00012

15 Hou, W. & Ji, Z. GeneTuring tests GPT models in genomics. BioRxiv (2023). 10.1101/2023.03.11.532238

16 Duong, D. & Solomon, B. D. Analysis of large-language model versus human performance for genetics questions. Eur J Hum Genet (2023). 10.1038/s41431-023-01396-8

17 Hou, W. & Ji, Z. Reference-free and cost-effective automated cell type annotation with GPT-4 in single-cell RNA-seq analysis. BioRxiv (2023). 10.1101/2023.04.16.537094

18 Dziadowicz, S. et al. Bone Marrow Stroma-induced Transcriptome and Regulome Signatures of Multiple Myeloma. Cancers 14, 927 (2022).

19 Guo, N. L. et al. A Predictive 7-Gene Assay and Prognostic Protein Biomarkers for Non-small Cell Lung Cancer. EBioMedicine 32, 102–110 (2018). 10.1016/j.ebiom.2018.05.025

20 Ye, Q. et al. Molecular Analysis of ZNF71 KRAB in Non-Small-Cell Lung Cancer. International journal of molecular sciences 22 (2021). 10.3390/ijms22073752

21 Qing Ye, J. H., Kathleen Summers, Brianne Falatovich, Marieta Gencheva, Timothy D. Eubank,Alexey V. Ivanov*, Nancy Lan Guo*. Multi-omics immune interaction networks in lung cancer tumorigenesis, proliferation, and survival. International journal of molecular sciences Molecular Oncology (2022).

22 Grossman, R. L. et al. Toward a Shared Vision for Cancer Genomic Data. N Engl J Med 375, 1109–1112 (2016). 10.1056/NEJMp1607591

23 Li, D., Harrison, J. K., Purushotham, D. & Wang, T. Exploring genomic data coupled with 3D chromatin structures using the WashU Epigenome Browser. Nat Methods 19, 909–910 (2022). 10.1038/s41592-022-01550-y

24 Ge, S. X., Jung, D. & Yao, R. ShinyGO: a graphical gene-set enrichment tool for animals and plants. Bioinformatics 36, 2628–2629 (2020). 10.1093/bioinformatics/btz931

25 Mezheyeuski, A. et al. Multispectral imaging for quantitative and compartment-specific immune infiltrates reveals distinct immune profiles that classify lung cancer patients. J Pathol 244, 421–431 (2018). 10.1002/path.5026

26 DepMap. (In Broad: figshare, 2020).

27 Guo, L., Cukic, B. & Singh, H. Predicting Fault Prone Modules by the Dempster-Shafer Belief Networks. Proceedings of 18th IEEE International Conference on Automated Software Engineering (ASE’03), 249–252 (2003).

28 Xu, J. Y. et al. Integrative Proteomic Characterization of Human Lung Adenocarcinoma. Cell 182, 245–261.e217 (2020). 10.1016/j.cell.2020.05.043

29 Ahmadinejad, N. et al. Accurate Identification of Subclones in Tumor Genomes. Mol Biol Evol 39 (2022). 10.1093/molbev/msac136

30 Dang, H. X. et al. ClonEvol: clonal ordering and visualization in cancer sequencing. Ann Oncol 28, 3076–3082 (2017). 10.1093/annonc/mdx517

31 Chen, W. C., Hu, G. & Hazlehurst, L. A. Contribution of the bone marrow stromal cells in mediating drug resistance in hematopoietic tumors. Curr Opin Pharmacol 54, 36–43 (2020). 10.1016/j.coph.2020.08.006

32 Hideshima, T. & Anderson, K. C. Signaling Pathway Mediating Myeloma Cell Growth and Survival. Cancers 13, 216 (2021). 10.3390/cancers13020216

33 Barski, A. et al. High-resolution profiling of histone methylations in the human genome. Cell 129, 823–837 (2007). 10.1016/j.cell.2007.05.009

34 Creyghton, M. P. et al. Histone H3K27ac separates active from poised enhancers and predicts developmental state. Proc Natl Acad Sci U S A 107, 21931–21936 (2010). 10.1073/pnas.1016071107

35 Hu, G. et al. Transformation of Accessible Chromatin and 3D Nucleome Underlies Lineage Commitment of Early T Cells. Immunity 48, 227–242 e228 (2018). 10.1016/j.immuni.2018.01.013

36 Maddigan, P. & Susnjak, T. Chat2VIS: Generating Data Visualizations via Natural Language Using ChatGPT, Codex and GPT-3 Large Language Models. Ieee Access 11, 45181–45193 (2023). 10.1109/Access.2023.3274199

37 Wang, L., Ge, X., Liu, L. & Hu, G. Code Interpreter for Bioinformatics: Are We There Yet? Ann Biomed Eng (2023). 10.1007/s10439-023-03324-9

38 Yang, Z. et al. The Dawn of LMMs: Preliminary Explorations with GPT-4V(ision). arXiv (2023). 10.48550/arXiv.2309.17421

39 Rose, D. et al. Visual Chain of Thought: Bridging Logical Gaps with Multimodal Infillings. (2023). 10.48550/arXiv.2305.02317

40 McBee, J. C. et al. Interdisciplinary Inquiry via PanelGPT: Application to Explore Chatbot Application in Sports Rehabilitation. medRxiv (2023). 10.1101/2023.07.23.23292452

