## Supplementary File 1 for "Bioinformatics Illustrations Decoded by ChatGPT: The Good, The Bad, and The Ugly"

Prompts used to guide ChatGPT to interpret bioinformatics illustrations

### The “DE gene” case study

#### Prompts for the initial assessment

Acting as a bioinformatics professional proficient in RNA-Seq data analysis, you are interpreting a figure to better appreciate transcription response of multiple myeloma cells (MMs) to bone marrow stromal cells (BMSCs). You also have extensive laboratory research experience in bone marrow tumor microenvironment of MM. In this task, expression profile of MM cell line RMPI8226 in trans-well coculture (T) with a BMSC cell line was compared to in monoculture (M). The attached figure illustrates the differential expression between the two conditions. Please do the following:

1. Provide an overview on each panel.
2. Are there any specific suggestions to improve the data presentation of the figure?
3. Draft a title and figure legend. Make sure to include details.
4. Draft a paragraph to describe the findings from the figure. This is one paragraph for result part of a manuscript. 150 words or less.

#### Prompts for the in-depth assessment

Acting as a bioinformatics professional proficient in RNA-Seq data analysis, you are interpreting a figure to better appreciate transcription response of multiple myeloma cells (MMs) to bone marrow stromal cells (BMSCs). You also have extensive laboratory research experience in bone marrow tumor microenvironment of MM. In this task, expression profile of MM cell line RMPI8226 in trans-well coculture (T) with a BMSC cell line was compared to in monoculture (M). The attached figure illustrates the differential expression between the two conditions. Please do the following:

1. Provide an overview on each panel.
2. For panel A, what do the numbers in parentheses mean?
3. For panel B, the positions of SOCS3 and JUNB are indicated by arrow heads. Estimate the values from the x-axis for the two genes. Are the two genes up-regulated or down-regulated? Based on your findings in the literature, why are the two genes chosen and are these results expected?
4. For panel C, pick the top two pathways and explain the criteria. It is not indicated on the plot whether upregulated or downregulated genes were used for the pathway enrichment analysis. However, by correlating the findings to existing knowledge in the literature, which group of genes were likely used?
5. Are there any specific suggestions to improve the data presentation of the figure?
6. Draft a title and figure legend. Make sure to include details.
7. Draft a paragraph to describe the findings from the figure. This is one paragraph for result part of a manuscript. 150 words or less.

### Prompts for the “ZNF71” case study

Acting as a bioinformatics professional. Please do the following:
1. Provide an overview on each panel.
2. Panel A: Is ZNF71 KRAB favorable or unfavorable? Why?
3. Panel B: Is ZNF71 KRAB resistant or sensitive to docetaxel? Why and be quantitative?
4. Panel C: Which genes are associated with ZNF71 in LUAD tumors? Which genes are associated with ZNF71 in non-cancerous adjacent tissues?
5. Panel D: What is the R value in Panel C? What is the indication on the expression of CD27 and PQ-401 resistance?

### Prompts for the “Clonal Evolution” case study

Acting as a bioinformatics professional proficient in cancer genomics data analysis, you will help interpret a figure to better appreciate clonal evolution in multiple myeloma (MM). The attached figure illustrates the results of analyzing somatic mutations detected in a pair of primary and recurrent tumor samples from a patient diagnosed with MM. Please do the following:

1. Provide an overview on each panel. Include the type of plot in the description, e.g., scatter plot, bar plot, etc.
2. Panel A. Identify the number of clusters and their colors. Sort the clusters by size.
3. Panel B. Identify clusters showing an increase in VAF from primary to recurrent.
4. Panels C and D. Identify the major cluster at the time of sample taken for each panel.
5. Panels E and F. Sort the clusters by size for each panel.
6. Panel G: Identify cluster at the root and clusters are the leaves.
7. Are there any specific suggestions to improve the data presentation of the figure?
8. Draft a title and figure legend. Make sure to include details.
9. Draft a paragraph to describe the findings from the figure. This is one paragraph for result part of a manuscript. 150 words or less.

### The “YY1” case study

#### Prompts for the initial assessment

Acting as a bioinformatics professional proficient in RNA-Seq, ChIP-seq, and 3D chromatin analysis, you will help interpret a figure to appreciate transcription regulation at chromatin level for YY1 in GM12878. The attached figure is a screen shot from WashU genome browser, encompassing gene annotation, expression data, ChIP-Seq data and chromatin-chromatin interaction data, for a genomic region enclosing YY1. The aim is to infer transcriptional regulatory mechanisms for YY1 from the epigenomic data. Please do the following.

1. Go over the tracks and explain specific findings addressing the aim.
2. Are there any specific suggestions to improve the data presentation of the figure?
3. Draft a title and figure legend. Make sure to include details.
4. Draft a paragraph to describe the findings from the figure. This is one paragraph for result part of a manuscript. 150 words or less.

#### Prompts for the in-depth assessment

Acting as a bioinformatics professional proficient in RNA-Seq, ChIP-seq, and 3D chromatin analysis, you will help interpret a figure to appreciate transcription regulation at chromatin level for YY1 in GM12878. The attached figure is a screen shot from WashU genome browser, encompassing gene annotation, expression data, ChIP-Seq data and chromatin-chromatin interaction data, for a genomic region enclosing YY1. The aim is to infer transcriptional regulatory mechanisms for YY1 from the epigenomic data. You will be guided with questions to identify findings addressing the aim. Please do the following.

1. Go over the RefSeq gene track and spell out the genes shown on the plot. Explain the presentation of gene structure. For examples what does the arrowhead mean.
2. Check the expression data. What do “(+)” and “(-)” mean? Which genes are expressed and why?
3. Check the ChIP-seq data. Explain the association of H3K4me3, H3K27ac, and H3K27me3 with gene expression. Based on the three histone modifications, which genes are likely expressed, and which are not likely expressed?
4. Three groups of genomic regions are highlighted: “A”s in light blue, “I”s in pink, and “E”s in green. Identify the number of regions for each group.
5. For genomic regions in “A”s, what are their shared features in terms of positional relationship with genes and histone modification patterns? What is their potential function in transcription regulation?
6. How about genomic regions in “I”s? What is their potential function in transcription regulation?
7. How about genomic regions in “E”s? What is their potential function in transcription regulation?
8. On the heatmap for chromatin-chromatin interactions, there are several circles each connected with dashed lines. Explain what the circle means and how about the dashed lines connected to the circle. What are the common features of the circles in terms of indicated genomic region(s)?
9. There are two group of circles by colors. Identify the number of circles for each group.
10. Focus the circles in blue, identify their indicated interacting genomic regions.
11. Focus the circles in green, identify their indicated interacting genomic regions.
12. Are there any specific suggestions to improve the data presentation of the figure?
13. Draft a title and a legend for this figure based on our previous discussion. Make sure to include details.
14. Draft a paragraph to describe the findings from our previous discussion on transcription regulation of YY1. This is one paragraph for result part of a manuscript. 150 words or less.
