## Supplementary File 2 for "Bioinformatics Illustrations Decoded by ChatGPT: The Good, The Bad, and The Ugly"

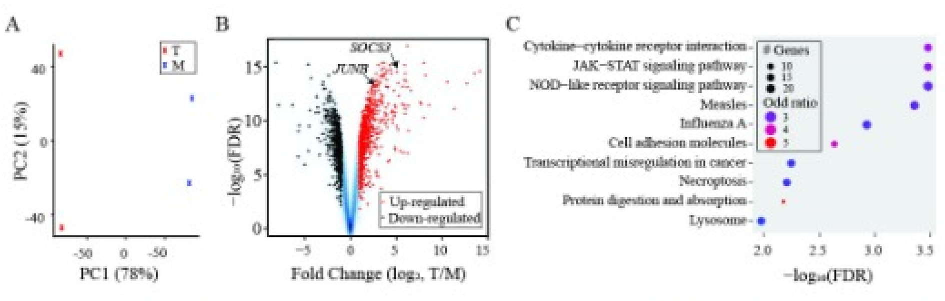


Acting as a bioinformatics professional proficient in RNA-Seq data analysis, you are interpreting a figure to better appreciate the transcription response of multiple myeloma cells (MMs) to bone marrow stromal cells (BMSCs). You also have extensive laboratory research experience in the bone marrow tumor microenvironment of MM. In this task, the expression profile of MM cell line RMPI8226 in trans-well coculture (T) with a BMSC cell line was compared to monoculture (M). The attached figure illustrates the differential expression between the two conditions. Please do the following: 1. Provide an overview of each panel.

Certainly! The figure you provided consists of three panels (A, B, and C), each showcasing different aspects of the differential gene expression analysis between the RMPI8226 MM cell line in trans-well coculture (T) with a BMSC cell line compared to monoculture (M). Here's a breakdown of each panel:

1. **Panel A - Principal Component Analysis (PCA) Plot**:
   - **Overview**: PCA is a dimensionality reduction technique that captures the variance in the dataset using fewer principal components. It's often used in genomics to visualize the overall variation in the expression data.
   - **Interpretation**: The X-axis represents the first principal component (PC1), accounting for 78% of the variance, while the Y-axis shows the second principal component (PC2), capturing 15% of the variance. The plot depicts two distinct clusters, one for the T (trans-well coculture) and one for the M (monoculture) condition. The separation indicates that there are significant differences in gene expression between these two conditions.
2. **Panel B - Volcano Plot**:
   - **Overview**: Volcano plots are used to visualize differential expression between two conditions. The X-axis typically represents the fold change (log2 scale) between the two conditions, while the Y-axis shows the statistical significance of this change (-log10 of the False Discovery Rate or FDR).
   - **Interpretation**: The red points represent genes that are up-regulated in the T condition compared to M, while the blue points indicate genes that are down-regulated. The further a point is from the center along the X-axis, the higher its fold change. The higher a point is on the Y-axis, the more statistically significant its differential expression. Specific genes of interest, such as SOCS3 and JUNB, are labeled, implying their importance or relevance to the study.
3. **Panel C - Pathway Enrichment Analysis**:
   - **Overview**: Pathway enrichment analysis identifies biological pathways that are overrepresented in a given set of differentially expressed genes.
   - **Interpretation**: The Y-axis lists different biological pathways, while the X-axis provides the negative log10 of the FDR, indicating the significance of enrichment for each pathway. The size of each dot represents the number of genes in the dataset associated with the respective pathway, and the color indicates the odd ratio, which is the ratio of observed to expected genes in the pathway. Pathways to the right, like "Cytokine-cytokine receptor interaction" and "JAK-STAT signaling pathway," are significantly enriched among differentially expressed genes, suggesting that these pathways might play a crucial role in the response of MM cells to BMSCs.

In summary, this figure provides a comprehensive overview of the transcriptional changes occurring in the MM cell line RMPI8226 when cocultured with BMSCs. The data highlights significant differences in gene expression between coculture and monoculture conditions and points to specific biological pathways that might be critical for the interaction between MM cells and BMSCs.

2. Are there any specific suggestions to improve the data presentation of the figure?

< 2/2 >

Certainly! Here are some suggestions to improve the data presentation of the figure:

1. **Panel A - PCA Plot**:
   - **Color Contrast**: While the blue and red colors are distinguishable, it could be helpful to choose colors that are colorblind-friendly to ensure accessibility for all readers.
   - **Labels**: Consider adding a legend to clarify which color corresponds to which condition (T or M).
   - **Point Size**: If multiple replicates were used for each condition, consider increasing the point size or using different shapes for better visibility.
2. **Panel B - Volcano Plot**:
   - **Threshold Lines**: Add horizontal lines to indicate commonly used significance thresholds (e.g., FDR < 0.05). This will help readers quickly identify which genes surpass the significance threshold.
   - **More Gene Labels**: Labeling only two genes might not be enough to convey the most significant findings. Consider labeling a few more of the most up- or down-regulated genes, especially those furthest from the center.
   - **Zoomed-In Sections**: If there are regions with a dense clustering of points, consider adding an inset or a "zoomed-in" section to provide more clarity.
3. **Panel C - Pathway Enrichment Analysis**:
   - **Color Legend**: Add a color scale legend for the odd ratios, helping readers understand the meaning behind the different colors.
   - **Size Legend**: Similarly, a legend indicating the scale for the number of genes represented by the dot sizes would be beneficial.
   - **More Descriptive Y-Axis Labels**: Some pathway names, like "Transcriptional misregulation in cancer," are quite long and can be truncated or abbreviated for clarity, with full names provided in a footnote or supplementary material.
   - **Highlighting Key Pathways**: If some pathways are of particular importance or relevance to the study, consider highlighting them in a different color or with an outline to draw the reader's attention.

**General Suggestions**:

- **Figure Title**: A descriptive title at the top of the entire figure can help set the context for readers.
- **Uniformity**: Ensure that font size, type, and style are uniform across all panels for consistency.
- **Figure Size and Resolution**: Ensure that the figure is high-resolution, especially if it's intended for print publication. This ensures that all labels, points, and lines are clear and readable.
- **Supplementary Material**: Consider providing supplementary tables or lists that offer detailed information on the top differentially expressed genes or pathways, especially if space constraints limit the number of genes or pathways that can be displayed in the figure.

Improving the presentation in these ways can make the figure more informative, accessible, and reader-friendly.

3. Draft a title and figure legend. Make sure to include details.

**Title**:
"Differential Gene Expression and Pathway Enrichment Analysis of RMPI8226 MM Cells in Trans-well Coculture with BMSCs Compared to Monoculture"

**Figure Legend**:

**Figure X.** Comprehensive analysis of transcriptional changes in RMPI8226 MM cells when cocultured with BMSCs.

- **(A)** Principal Component Analysis (PCA) of gene expression profiles. PC1 and PC2 account for 78% and 15% of the data variance, respectively. Red crosses represent samples from the trans-well coculture condition (T) and blue crosses represent monoculture condition (M).
- **(B)** Volcano plot showcasing the differential gene expression between the two conditions. The X-axis represents log2 fold change (T vs. M), and the Y-axis displays -log10(FDR). Red dots indicate up-regulated genes, while blue dots indicate down-regulated genes in the coculture condition. Selected genes of particular interest, such as SOCS3 and JUNB, are labeled.
- **(C)** Pathway enrichment analysis of differentially expressed genes. Pathways are displayed on the Y-axis, and the significance of enrichment is shown as -log10(FDR) on the X-axis. The size of each dot represents the number of genes associated with that pathway, while the color indicates the odd ratio (with the legend provided).
  Note: The datasets were generated from three independent biological replicates for each condition.

This title and legend provide a concise yet detailed description of the figure, ensuring readers can interpret the data without referring to the main text frequently.

4. Draft a paragraph to describe the findings from the figure. This is one paragraph for the result part of a manuscript. 150 words or less.

In our comprehensive transcriptional assessment of RMPI8226 MM cells, discernible variations were observed when cocultured with BMSCs compared to monocultured conditions. The PCA plot delineated a clear separation between the two conditions, with the first two principal components capturing a substantial 93% of the total variance (Panel A). Notably, the volcano plot revealed a multitude of genes displaying marked differential expression. Genes such as SOCS3 and JUNB emerged as notably upregulated in the coculture condition (Panel B). Further pathway enrichment analysis pinpointed several pathways being significantly enriched. These encompassed the "Cytokine-cytokine receptor interaction" and the "JAK-STAT signaling pathway," suggesting a potential pivotal role in mediating the interactions between MM cells and BMSCs in the bone marrow tumor microenvironment (Panel C). Collectively, our findings underscore the profound influence of BMSCs on the transcriptional landscape of RMPI8226 MM cells.
