## Supplementary File 3 for "Bioinformatics Illustrations Decoded by ChatGPT: The Good, The Bad, and The Ugly"

GPT-4

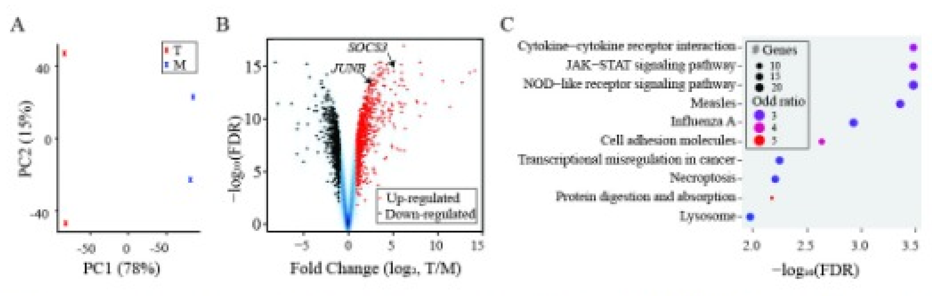


Acting as a bioinformatics professional proficient in RNA-Seq data analysis, you are interpreting a figure to better appreciate the transcription response of multiple myeloma cells (MMs) to bone marrow stromal cells (BMSCs). You also have extensive laboratory research experience in the bone marrow tumor microenvironment of MM. In this task, the expression profile of MM cell line RMPI8226 in trans-well coculture (T) with a BMSC cell line was compared to monoculture (M). The attached figure illustrates the differential expression between the two conditions. Please do the following: 1. Provide an overview of each panel.

Of course, based on the provided figure and your description, here's an interpretation of each panel:

**Panel A: Principal Component Analysis (PCA) Plot**

- This plot provides a visualization of the overall variance in the gene expression data between the two conditions: monoculture (M) and trans-well coculture (T) of MM cells with BMSCs.
- The x-axis (PC1) explains 78% of the variance, while the y-axis (PC2) explains 15%.
- Each point on the plot represents a sample, and the spatial distance between points indicates the degree of similarity in gene expression patterns.
- There's a clear separation between the two conditions, with the T (in red) and M (in blue) clusters being distinct. This indicates significant differences in gene expression between the two conditions.

**Panel B: Volcano Plot**

- This plot visualizes the differential expression of genes between the T and M conditions.
- The x-axis shows the fold change in gene expression (log2 scale), with genes to the right being up-regulated in the T condition and those to the left being down-regulated.
- The y-axis displays the statistical significance of these changes in terms of -log10(FDR), with higher values indicating higher significance.
- Genes that are highly up-regulated and statistically significant are in the upper right corner (in red), and those that are highly down-regulated and statistically significant are in the upper left corner (in blue).
- Two genes, "SOCS3" and "JUNB", are specifically labeled, suggesting they might be of particular interest or have shown significant differential expression between the two conditions.

**Panel C: Pathway Enrichment Analysis**

- This panel provides insights into the biological pathways that are enriched among the differentially expressed genes.
- The x-axis indicates the significance of the enrichment in terms of -log10(FDR).
- Each dot represents a specific biological pathway, and the size of the dot represents the number of genes associated with that pathway.
- The color of each dot represents the odd ratio, indicating the strength of the association between the differentially expressed genes and the given pathway.
- Pathways such as "Cytokine-cytokine receptor interaction," "JAK-STAT signaling pathway," and "NOD-like receptor signaling pathway" appear to be highly significant and may play critical roles in the response of MM cells to BMSCs.

**Overall Interpretation:**
The provided figure paints a comprehensive picture of the transcriptional changes that occur in MM cells when cocultured with BMSCs. The clear separation in the PCA plot emphasizes the substantial difference in gene expression profiles between the two conditions. The volcano plot pinpoints specific genes that are highly up-regulated or down-regulated. Lastly, the pathway enrichment analysis provides a biological context, suggesting that certain signaling pathways and processes might be pivotal in mediating the interactions between MM cells and BMSCs in the bone marrow tumor microenvironment.

2. Are there any specific suggestions to improve the data presentation of the figure?

Certainly! While the figure is informative, there are always potential improvements that can enhance clarity and ease of interpretation. Here are some suggestions to improve the data presentation:

1. **Panel A (PCA Plot)**:
   - **Legend and Labels**: Ensure the legend is clear and easy to distinguish between the T and M conditions. If not already done, consider using distinct colors or shapes for each condition.
   - **Percent Variance**: For clarity, explicitly label the percent variance explained by each principal component directly on the axes (e.g., "PC1 (78% variance)").
   - **Zooming**: If there are more samples, consider zooming into regions where data points overlap to provide better clarity.
2. **Panel B (Volcano Plot)**:
   - **Threshold Lines**: Add horizontal lines to indicate commonly used FDR thresholds (e.g., FDR = 0.05) and vertical lines for fold change thresholds (e.g., 2-fold). This will help readers quickly identify genes of interest.
   - **Highlight Key Genes**: While "SOCS3" and "JUNB" are labeled, consider highlighting other genes that might be of biological significance or interest.
   - **Color Gradient**: Instead of just red and blue, use a gradient to represent the degree of up-regulation or down-regulation, which can provide more nuance to the visualization.
3. **Panel C (Pathway Enrichment Analysis)**:
   - **Y-axis Labeling**: Explicitly label the y-axis with the pathway names for better readability. If space is limited, rotating the text or using abbreviations can help.
   - **Interactive Features**: If presenting digitally, consider using an interactive plot where hovering over a dot provides more details about the pathway (e.g., gene names, specific p-values).
   - **Order of Pathways**: Rearrange the pathways in decreasing order of significance, with the most significant pathways at the top.
4. **General**:
   - **Consistent Color Scheme**: Ensure that the color schemes used in different panels are consistent. For instance, if red represents the T condition in Panel A, it should represent the same in Panels B and C.
   - **Title and Descriptive Captions**: Provide a clear and concise title for the figure, and consider adding descriptive captions below each panel to guide the reader.
   - **Font Size and Style**: Ensure that all text is legible, especially when the figure is resized. Use consistent font styles and sizes across panels.
   - **High-Resolution Export**: If the figure is intended for publication, ensure it is exported in high resolution with lossless compression.
5. **Additional Data**: If there's supplementary data or detailed results for some genes/pathways, consider providing an appendix or a link to a supplementary file. This ensures that readers interested in delving deeper have access to detailed information.

Improving the presentation not only makes the data more aesthetically pleasing but also aids in conveying the findings more effectively to the audience.

3. Draft a title and figure legend. Make sure to include details.

**Title**:
"Differential Gene Expression and Pathway Enrichment Analysis in MM Cells Co-cultured with BMSCs"

**Figure Legend**:

**Figure 1**: Transcriptional changes observed in MM cell line RMPI8226 when co-cultured with a BMSC cell line compared to monoculture conditions.

- **(A)** Principal Component Analysis (PCA) plot showcasing variance in gene expression between MM cells in monoculture (M, shown in blue) and trans-well co-culture with BMSCs (T, shown in red). PC1 accounts for 78% of the variance, while PC2 accounts for 15%.
- **(B)** Volcano plot illustrating the differential expression of genes between the two conditions. The x-axis depicts fold change in gene expression (log2 scale), with genes on the right being up-regulated and those on the left being down-regulated in the T condition relative to M. The y-axis presents the statistical significance of these changes as -log10(FDR). Genes "SOCS3" and "JUNB" are labeled, indicating their noteworthy differential expression.
- **(C)** Pathway Enrichment Analysis indicating significant biological pathways associated with the differentially expressed genes. Each dot represents a pathway, with the x-axis denoting the significance of enrichment as -log10(FDR). Dot size corresponds to the number of genes in each pathway, while the color gradient signifies the odd ratio, reflecting the strength of the association.

Note: The data provides insights into the molecular mechanisms underlying MM cell interactions with the bone marrow stromal environment, highlighting the importance of specific genes and signaling pathways in this context.

4. Draft a paragraph to describe the findings from the figure. This is one paragraph for the result part of a manuscript. 150 words or less.

In our examination of the transcriptional response of the MM cell line RMPI8226 to BMSC coculture, distinct transcriptional shifts were evident. The Principal Component Analysis (PCA) in Figure 1A revealed a pronounced separation between monocultured MM cells and those co-cultured with BMSCs, with PC1 accounting for a striking 78% of the variance. Delving into individual gene expression differences, the volcano plot (Figure 1B) spotlighted several genes with significant differential expression, notably the up-regulation of "SOCS3" and "JUNB". Furthermore, pathway enrichment analysis shed light on potential biological implications of these transcriptional changes. As shown in Figure 1C, pathways related to "Cytokine-cytokine receptor interaction", "JAK-STAT signaling", and "NOD-like receptor signaling" were among the most significantly enriched, underscoring their potential roles in mediating MM cell interactions within the bone marrow stromal environment. These findings provide a foundational understanding of the molecular dialogue between MM cells and their surrounding niche.
