## Supplementary File 4 for "Bioinformatics Illustrations Decoded by ChatGPT: The Good, The Bad, and The Ugly"

GPT-4

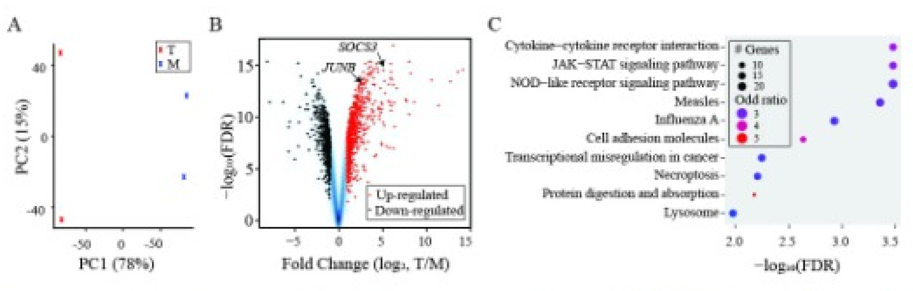


Acting as a bioinformatics professional proficient in RNA-Seq data analysis, you are interpreting a figure to better appreciate the transcription response of multiple myeloma cells (MMs) to bone marrow stromal cells (BMSCs). You also have extensive laboratory research experience in the bone marrow tumor microenvironment of MM. In this task, the expression profile of MM cell line RMPI8226 in trans-well coculture (T) with a BMSC cell line was compared to monoculture (M). The attached figure illustrates the differential expression between the two conditions. Please do the following: 1. Provide an overview of each panel.

Sure, based on the provided figure, here's an overview of each panel:

**Panel A: Principal Component Analysis (PCA) Plot**

- This plot represents a PCA analysis of the expression profiles from the MM cell line RPMI8226 when co-cultured with a BMSC cell line (T) versus when it's monocultured (M).
- PCA is a dimensionality reduction method used to visualize the variance in large datasets. Here, it helps us observe the overall transcriptional differences between the two conditions.
- The X-axis represents the first principal component (PC1) which captures 78% of the total variance, while the Y-axis represents the second principal component (PC2) capturing 15% of the variance.
- The blue and red crosses represent the transcriptional profiles of the monoculture (M) and trans-well co-culture (T) respectively. The clear separation between them suggests significant transcriptional differences between the two conditions.

**Panel B: Volcano Plot**

- This plot showcases the differential expression of genes between the MM cell line RPMI8226 in trans-well coculture (T) with a BMSC cell line compared to its monoculture (M).
- The X-axis represents the fold change (in log2 scale) of gene expression between the two conditions. Positive values indicate up-regulation in the co-culture, while negative values indicate down-regulation.
- The Y-axis represents the statistical significance of this change in a -log10 scale. Higher values mean more significant differences.
- Genes with higher statistical significance and higher fold change are located in the top right (up-regulated) and top left (down-regulated) corners. Two genes, SOCS3 and JUNB, are explicitly labeled, suggesting they might be of particular interest.
- The red dots represent up-regulated genes, and the blue dots represent down-regulated genes.

**Panel C: Pathway Enrichment Analysis**

- This plot displays the results of a pathway enrichment analysis, which determines which biological pathways are significantly represented among the differentially expressed genes.
- The Y-axis lists different pathways, while the X-axis shows the significance (-log10 of FDR, or False Discovery Rate) of each pathway's enrichment.
- The size of each dot represents the number of genes in the pathway, and the color represents the odd ratio, which indicates the likelihood that the genes from the dataset are part of a given pathway compared to random chance. A higher odd ratio suggests a stronger association.
- Several pathways are highlighted, including cytokine-cytokine receptor interaction, JAK-STAT signaling, and others. These pathways might play a crucial role in the interaction between MM cells and BMSCs in the bone marrow tumor microenvironment.

In summary, this figure provides a comprehensive overview of the transcriptional changes in MM cells when co-cultured with BMSCs, highlighting specific genes of interest and potential biological pathways influenced by this interaction.

2. Are there any specific suggestions to improve the data presentation of the figure?

Certainly! Here are some suggestions to improve the data presentation of the figure:

**Panel A: Principal Component Analysis (PCA) Plot**

1. **Label clarity**: Add clear labels or legends for the data points representing monoculture (M) and trans-well coculture (T) to help viewers quickly understand the plot.
2. **Proportion of variance**: Consider displaying the percentage of variance explained by other principal components (e.g., PC3, PC4) in a small inset or table, especially if they contribute significantly to the dataset's variance.
3. **Gridlines**: Adding subtle gridlines can help in estimating the coordinates of the points more precisely.

**Panel B: Volcano Plot**

1. **Significance threshold**: Add horizontal lines indicating common significance thresholds (e.g., p-value of 0.05) to quickly identify genes that surpass this threshold.
2. **Fold-change threshold**: Add vertical lines to mark common fold change thresholds (e.g., 2-fold or log2(2)) to easily identify highly upregulated or downregulated genes.
3. **Highlight key genes**: Besides SOCS3 and JUNB, consider highlighting or labeling other genes that might be of biological significance or those with extreme values of fold change and significance.
4. **Color choice**: Use a gradient color scale or another scheme to indicate the magnitude of fold change and significance, which can provide additional visual cues.

**Panel C: Pathway Enrichment Analysis**

1. **Interactive capabilities**: If presented online, consider adding interactive capabilities to the plot. Hovering over a point could display the specific genes involved in that pathway or show additional information about the pathway.
2. **Scale clarity**: For the color scale representing the odd ratio, consider adding a legend or scale bar for clarity.
3. **Sort pathways**: Sort pathways based on their significance or another meaningful metric to help readers quickly identify the most relevant pathways.
4. **Pathway details**: For pathways of particular interest, consider providing a small inset or additional figure detailing the specific genes from the dataset involved in that pathway.

**General suggestions**:

1. **Consistent color schemes**: Ensure that colors used are consistent across all panels if they represent the same data points or conditions.
2. **Resolution and clarity**: Ensure that the figure is of high resolution, especially if it's for print. All text, labels, and data points should be clear and legible.
3. **Figure caption**: Provide a comprehensive figure caption that briefly describes each panel and any experimental or analytical details that might be essential for interpretation.
4. **Size and scale**: Depending on where this figure will be presented (e.g., journal article, presentation, poster), adjust the size and scale to ensure it's easily interpretable without overwhelming the viewer.

These enhancements can make the figure more informative and user-friendly, aiding in clearer data interpretation.

3. Draft a title and figure legend. Make sure to include details.

**Title**:
"Differential Gene Expression and Pathway Enrichment Analysis of RPMI8226 Multiple Myeloma Cells Co-cultured with BMSCs"

**Figure Legend**:

**Figure X**: Transcriptional profiling and pathway analysis of RPMI8226 multiple myeloma cells in response to co-culture with bone marrow stromal cells.

**(A) Principal Component Analysis (PCA) Plot**: Transcriptional variance is depicted between the RPMI8226 MM cells in monoculture (M, blue crosses) and in trans-well coculture with BMSCs (T, red crosses). The X-axis displays the first principal component (PC1), accounting for 78% of the total variance, while the Y-axis displays the second principal component (PC2), capturing 15% of the variance.

**(B) Volcano Plot**: Differential gene expression between the two conditions is illustrated. The X-axis indicates the fold change in gene expression (log2 scale), with positive values showing up-regulation in the co-culture and negative values indicating down-regulation. The Y-axis represents the statistical significance of these changes in a -log10 scale. Genes with higher statistical significance and fold change are seen in the top corners. Specific genes of potential significance, such as SOCS3 and JUNB, are labeled.

**(C) Pathway Enrichment Analysis**: Significant biological pathways influenced by the differential gene expression are shown. Pathways are listed on the Y-axis, with their enrichment significance represented on the X-axis (-log10 of FDR). The size of each dot corresponds to the number of genes involved in that pathway, and the color indicates the odds ratio, with a darker hue suggesting a stronger association between the dataset genes and the pathway.

Note: FDR stands for False Discovery Rate, a method used to correct for multiple testing. The odds ratio quantifies the likelihood that the observed association between the dataset genes and a pathway is not due to random chance.

4. Draft a paragraph to describe the findings from the figure. This is one paragraph for the result part of a manuscript. 150 words or less.

In our exploration of the transcriptional response of RPMI8226 multiple myeloma cells (MMs) to bone marrow stromal cells (BMSCs), notable alterations in gene expression were observed (Figure X). A distinct separation in the Principal Component Analysis (PCA) plot highlighted the profound transcriptional differences between MM cells co-cultured with BMSCs and those in monoculture, with the first two principal components accounting for 93% of the total variance. Our volcano plot pinpointed genes like SOCS3 and JUNB as notably upregulated. Pathway enrichment analysis further elucidated the biological implications of these transcriptional changes. Key pathways, such as cytokine-cytokine receptor interaction, JAK-STAT signaling, and NOD-like receptor signaling, were found to be significantly enriched. These findings underscore the pivotal role of the bone marrow microenvironment in influencing the gene expression landscape of MM cells, providing insights into potential therapeutic targets and the intricate interplay between tumor cells and their surroundings.
