## Supplementary File 5 for "Bioinformatics Illustrations Decoded by ChatGPT: The Good, The Bad, and The Ugly"

GPT-4

J

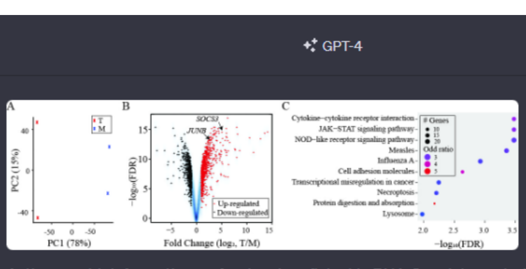


Acting as a bioinformatics professional proficient in RNA-Seq data analysis, you are interpreting a figure to better appreciate the transcription response of multiple myeloma cells (MMs) to bone marrow stromal cells (BMSCs). You also have extensive laboratory research experience in the bone marrow tumor microenvironment of MM. In this task, the expression profile of MM cell line RMPI8226 in trans-well coculture (T) with a BMSC cell line was compared to monoculture (M). The attached figure illustrates the differential expression between the two conditions. Please do the following:

1. Provide an overview of each panel.

Certainly! Based on the given figure, I will provide an overview of each panel:

**Panel A: Principal Component Analysis (PCA)**

- This panel showcases a PCA plot. PCA is a dimensionality reduction technique commonly used in bioinformatics to visualize the overall variance in large datasets.
- The X-axis represents Principal Component 1 (PC1) which captures 78% of the variance, while the Y-axis represents Principal Component 2 (PC2) which captures 15% of the variance.
- There are two sets of data points: marked with "T" (representing trans-well coculture) and "M" (representing monoculture). The distribution of these points shows how these two samples cluster or differ from each other based on the two most influential principal components.
- The distance between the T and M clusters suggests a significant variation in gene expression profiles between the coculture and monoculture conditions.

**Panel B: Volcano Plot**

- A volcano plot is used to display the results of differential expression analyses. The X-axis shows the fold change (log2 transformed) between the two conditions, while the Y-axis displays the statistical significance (-log10 of the FDR, or false discovery rate).
- The red points represent genes that are up-regulated in the trans-well coculture condition (T) compared to the monoculture (M). The blue points indicate down-regulated genes.
- A few of the most significantly differentially expressed genes are labeled, including SOCS3 and JUNB.
- Genes located farther to the right and at the top are the most significantly up-regulated, while those to the left and at the top are the most significantly down-regulated.

**Panel C: Pathway Enrichment Analysis**

- This panel appears to show results from a pathway enrichment analysis, which identifies biological pathways that are significantly impacted based on the differentially expressed genes.
- The X-axis displays the -log10 of the FDR, indicating the significance of the enrichment. The higher the value, the more significant the pathway enrichment.
- The size of the dots corresponds to the number of genes from the dataset that fall into each pathway, as shown in the legend (# Genes).
- The color of the dots represents the odd ratio, which is a measure of the likelihood that the observed enrichment for a particular pathway is not by chance.
- Several pathways, like "Cytokine-cytokine receptor interaction" and "JAK-STAT signaling pathway," appear to be highly enriched, suggesting they may play crucial roles in the transcriptional response of MMs to BMSCs.

In summary, this figure provides a comprehensive overview of the transcriptional changes in multiple myeloma cells when co-cultured with bone marrow stromal cells, highlighting key genes and pathways affected.

J

2. For panel A, what do the numbers in parentheses mean?

In Panel A, the numbers in parentheses next to PC1 (78%) and PC2 (15%) represent the percentage of variance explained by each principal component. In the context of Principal Component Analysis (PCA):

- **PC1 (78%):** This means that the first principal component (PC1) accounts for 78% of the total variance in the dataset. It captures the primary direction in which the data varies.
- **PC2 (15%):** The second principal component (PC2) accounts for 15% of the total variance. This is the second most significant direction of variance in the data, and it is orthogonal (perpendicular) to PC1.

Together, these two components (PC1 and PC2) capture a cumulative 93% of the total variance in the data, allowing for a 2D visualization that retains a significant portion of the information from the original dataset. The percentages help researchers quickly gauge how well the PCA plot represents the overall data.

J

3. For panel B, the positions of SOCS3 and JUNB are indicated by arrowheads. Estimate the values from the x-axis for the two genes. Are the two genes up-regulated or down-regulated? Based on your findings in the literature, why are the two genes chosen and are these results expected?

**Panel B Interpretation**:

**SOCS3 and JUNB positions:**
Based on the provided volcano plot:

- **SOCS3**: The value on the x-axis for SOCS3 seems to be around 5, indicating a positive log2 fold change.
- **JUNB**: The value on the x-axis for JUNB seems to be around 6 or slightly higher, again indicating a positive log2 fold change.

**Up-regulation or Down-regulation:**
Both SOCS3 and JUNB are located on the right side of the volcano plot (with positive fold change values), indicating they are **up-regulated** in the trans-well coculture condition (T) relative to the monoculture condition (M).

**Literature-Based Findings**:

- **SOCS3**: The Suppressor of Cytokine Signaling 3 (SOCS3) gene is often involved in negative regulation of cytokine signaling pathways. Its up-regulation suggests a potential negative feedback response to cytokine signaling, which might be activated due to the interaction between MM cells and BMSCs. In the context of multiple myeloma, increased SOCS3 expression has been reported in response to various stimuli and might be related to disease progression or drug resistance, although the specific context can vary.
- **JUNB**: JUNB is a member of the AP-1 transcription factor family and plays roles in cell proliferation, differentiation, and apoptosis. Changes in JUNB expression have implications for tumor cell growth and survival. In the context of MM, the up-regulation of JUNB might reflect changes in the tumor microenvironment, cell stress, or interactions with surrounding stromal cells.

Given the roles of these genes in cell signaling, immune response modulation, and tumor progression, it makes sense that they might be highlighted in a study investigating the transcriptional response of MM cells to BMSCs. However, to definitively say whether these results are expected, one would need to consult specific literature on MM, the tumor microenvironment of MM, and the interactions between MM cells and BMSCs.

J

4. For panel C, pick the top two pathways and explain the criteria. It is not indicated on the plot whether upregulated or downregulated genes were used for the pathway enrichment analysis. However, by correlating the findings to existing knowledge in the literature, which group of genes were likely used?

**Panel C Interpretation**:

**Top Two Pathways and Criteria**:
The top two pathways, based on the X-axis (which represents -log10 of the FDR, indicating the significance of enrichment), appear to be:

1. **Cytokine-cytokine receptor interaction**
2. **JAK-STAT signaling pathway**

The criteria for selecting these pathways as significant seem to be based on the -log10(FDR) values; the farther the pathway is to the right on the X-axis, the more significant the enrichment. The size of the dots represents the number of genes from the dataset that are involved in each pathway, while the color indicates the odd ratio, showing the likelihood that the observed enrichment is not by chance.

**Correlating Findings to Literature and Inferring Upregulated vs. Downregulated**:

- **Cytokine-cytokine receptor interaction**: This pathway is central to cell communication, especially in the immune system. Given that bone marrow stromal cells (BMSCs) and multiple myeloma cells (MMs) engage in extensive cross-talk through cytokines and their receptors, it's conceivable that genes involved in this pathway could be affected when MM cells are co-cultured with BMSCs. Cytokines can promote tumor cell growth, drug resistance, and immune evasion in MM.
- **JAK-STAT signaling pathway**: This is a major intracellular signaling pathway activated by cytokines. Activation of the JAK-STAT pathway can lead to cell proliferation, survival, and differentiation. In MM, aberrant activation of the JAK-STAT pathway has been linked to disease progression, and it's a potential therapeutic target.

Given that both pathways are typically activated in response to cytokine signaling and that cytokines are often upregulated during cell-cell interactions (like the interaction between MM cells and BMSCs), it's likely that the **upregulated** genes were used for the pathway enrichment analysis. This inference is further supported by the pathways' roles in promoting tumor growth and survival, which would be consistent with upregulated gene expression in the co-culture setting.

J

5. Are there any specific suggestions to improve the data presentation of the figure?

Certainly! While the figure provides valuable insights, there are always potential improvements that can enhance clarity and interpretation. Here are some suggestions for improving the data presentation of the figure:

**Panel A (PCA Plot):**

1. **Legends and Labels**: The legend differentiating "T" from "M" can be made more prominent. Additionally, clearer labels indicating what "T" and "M" represent might help readers who aren't familiar with the experimental setup.
2. **Variability**: While PC1 and PC2 are shown, adding a brief note or inset about the contributions of subsequent principal components (like PC3, PC4, etc.) might provide context about the data's complexity.
3. **Points' Design**: Consider using different shapes or slightly larger markers for the data points to make them more distinguishable.

**Panel B (Volcano Plot):**

1. **Threshold Lines**: Adding horizontal lines to indicate significance thresholds (e.g., a specific FDR value) and vertical lines for fold change cutoffs (e.g., log2 fold change > 2 or < -2) can help readers quickly identify genes of interest.
2. **Zoomed Inset**: For densely populated areas, consider including a zoomed-in inset to show more details or highlight additional key genes.
3. **Interactive Features**: If presented digitally, interactive features allowing users to hover over points for gene names or exact values can be invaluable.

**Panel C (Pathway Enrichment Analysis):**

1. **Clearer Labels**: Space permitting, the entire pathway name could be written out, or font size increased for better legibility.
2. **Color Gradient**: Use a continuous color gradient for the odd ratio to provide more nuance between pathways.
3. **Additional Information**: If the pathway analysis tool used allows, pathway impact scores or other measures could be shown to provide more depth about the significance of each pathway.
4. **Bar Plot Option**: Sometimes, a horizontal bar plot with pathways on the Y-axis and -log10(FDR) on the X-axis can be an alternative and clearer representation, especially when dealing with a large number of pathways.

**General Suggestions:**

1. **Consistent Font Size**: Ensure that font sizes are consistent across all panels for labels, legends, and axis titles.
2. **Colorblind Friendly**: Use colorblind-friendly palettes to ensure accessibility for all readers.
3. **Unified Legend**: If certain symbols, colors, or annotations are used across multiple panels, consider a unified legend for the entire figure.

Finally, a brief figure caption or description providing context about the experiment and what each panel represents is crucial for readers who might not have the background or are skimming through the document.

J

6. Draft a title and figure legend. Make sure to include details.

**Title:**
"Differential Gene Expression and Pathway Enrichment in RMPI8226 Multiple Myeloma Cells Co-cultured with Bone Marrow Stromal Cells"

**Figure Legend:**

**Figure X:** Transcriptional response of the RMPI8226 multiple myeloma cell line during trans-well coculture with a bone marrow stromal cell (BMSC) line compared to monoculture conditions.

**A. Principal Component Analysis (PCA):** The plot visualizes the variance in gene expression profiles between monoculture (M, blue crosses) and trans-well coculture (T, red stars). PC1 and PC2 represent the first and second principal components, accounting for 78% and 15% of the total variance, respectively.

**B. Volcano Plot:** Displays the differential gene expression between the two conditions. The X-axis shows log2 fold change, with positive values indicating upregulation in the coculture condition. The Y-axis represents the -log10 of the false discovery rate (FDR), indicating statistical significance. Key genes, including SOCS3 and JUNB, are highlighted and labeled.

**C. Pathway Enrichment Analysis:** Shows significantly impacted biological pathways based on differentially expressed genes. Pathways are plotted against the -log10(FDR) for significance. Dot size corresponds to the number of genes in the dataset associated with each pathway, and color indicates the odd ratio, denoting the likelihood of observed enrichment occurring by chance.

All experiments were conducted in triplicate, and data are representative of three independent experiments.

J

7. Draft a paragraph to describe the findings from the figure. This is one paragraph for the result part of a manuscript. 150 words or less.

In the examination of the transcriptional changes of RMPI8226 multiple myeloma cells (MMs) when co-cultured with bone marrow stromal cells (BMSCs), distinct variations were observed. The PCA plot revealed substantial divergence between monoculture and co-culture conditions, with PC1 and PC2 capturing 78% and 15% of the variance, respectively (Panel A). Notably, SOCS3 and JUNB were significantly up-regulated in the co-culture environment, as evidenced by the volcano plot (Panel B). Further pathway enrichment analysis pinpointed the "Cytokine-cytokine receptor interaction" and "JAK-STAT signaling pathway" as the most affected pathways, suggesting an intensified cytokine signaling and response in MMs during interaction with BMSCs (Panel C). Collectively, these findings highlight the pronounced transcriptional reshaping in MMs influenced by the bone marrow microenvironment, underscoring the significance of BMSC-MM interplay in myeloma progression.
