## Supplementary File 6 for "Bioinformatics Illustrations Decoded by ChatGPT: The Good, The Bad, and The Ugly"

GPT-4

J

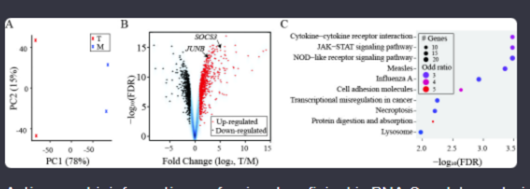


Acting as a bioinformatics professional proficient in RNA-Seq data analysis, you are interpreting a figure to better appreciate the transcription response of multiple myeloma cells (MMs) to bone marrow stromal cells (BMSCs). You also have extensive laboratory research experience in the bone marrow tumor microenvironment of MM. In this task, the expression profile of MM cell line RMPI8226 in trans-well coculture (T) with a BMSC cell line was compared to monoculture (M). The attached figure illustrates the differential expression between the two conditions. Please do the following:

1. Provide an overview of each panel.

Certainly! Let's go through each panel:

### Panel A:

- **Title**: Principal Component Analysis (PCA) of Transcriptional Profiles.
- **Description**: This panel showcases the PCA plot, which is a common method used to reduce dimensionality and visualize the variation in large datasets. The x-axis represents the first principal component (PC1) accounting for 78% of the variance, and the y-axis represents the second principal component (PC2) accounting for 15% of the variance.
- **Observation**: Two distinct clusters are evident. The blue cluster represents MM cells in trans-well coculture (T) with BMSCs, while the red cluster represents MM cells in monoculture (M). The clear separation indicates significant transcriptional differences between the two conditions.

### Panel B:

- **Title**: Volcano Plot of Differentially Expressed Genes.
- **Description**: The volcano plot visually displays differential gene expression between the two conditions. The x-axis represents the fold change in expression (log2 scale of T/M), where positive values indicate up-regulation in coculture and negative values indicate down-regulation. The y-axis is the negative logarithm (base 10) of the false discovery rate (FDR), which corrects for multiple testing; higher values mean more statistical significance.
- **Observation**: The red points represent genes that are up-regulated in coculture, while the blue points represent down-regulated genes. Two genes, SOCS3 and JUNB, are labeled, suggesting they are of particular interest, possibly due to their high fold-change and/or statistical significance.

### Panel C:

- **Title**: Pathway Enrichment Analysis.
- **Description**: This panel shows the results of a pathway enrichment analysis. The x-axis represents the negative logarithm (base 10) of the FDR, indicating the significance of enrichment. The y-axis lists various biological pathways. The size of the dot corresponds to the number of genes in the dataset associated with that pathway, and the color indicates the odd ratio, which signifies the likelihood of observing the given number of genes in that pathway by chance.
- **Observation**: Several pathways are enriched in the dataset, suggesting their potential relevance to the response of MM cells to BMSCs. For instance, "Cytokine-cytokine receptor interaction" and "JAK-STAT signaling pathway" appear significant, indicating a possible involvement of these pathways in the interaction between MM cells and BMSCs.

In summary, the figure provides comprehensive insights into the transcriptional response of MM cells when cocultured with BMSCs. The distinct clustering in PCA, differentially expressed genes in the volcano plot, and the enriched pathways collectively highlight the profound influence of BMSCs on the MM cell line's gene expression.

J

2. For panel A, what do the numbers in parentheses mean?

In Panel A, the numbers in parentheses next to PC1 and PC2 denote the proportion of the total variance in the dataset that is captured by each respective principal component. Specifically:

- **PC1 (78%)**: This means that the first principal component (PC1) captures 78% of the total variance in the dataset.
- **PC2 (15%)**: Similarly, the second principal component (PC2) captures 15% of the total variance.

In other words, the two dimensions shown in the PCA plot (PC1 and PC2) together account for 93% (78% + 15%) of the total variability in the gene expression data. The higher the percentage, the more variation that particular principal component captures, and thus it's more informative for differentiating between different conditions or groups in the dataset.

J

3. For panel B, the positions of SOCS3 and JUNB are indicated by arrowheads. Estimate the values from the x-axis for the two genes. Are the two genes up-regulated or down-regulated? Based on your findings in the literature, why are the two genes chosen and are these results expected?

### Panel B Analysis:

Estimating Values:

- **SOCS3**: This gene appears to have a fold change value (from the x-axis) of slightly above 5.
- **JUNB**: This gene's fold change value seems to be around 6 or slightly above.

Up-regulation or Down-regulation:

Both SOCS3 and JUNB are on the right side of the 0 value on the x-axis, which means both genes have a positive fold change. Hence, both SOCS3 and JUNB are **up-regulated** in the trans-well coculture (T) compared to the monoculture (M).

### Literature-based Insights:

- **SOCS3 (Suppressor of Cytokine Signaling 3)**:
  - **Role**: SOCS3 is known to play a crucial role in the negative regulation of cytokine signaling. It's part of a classical feedback mechanism to prevent excessive activation of JAK-STAT signaling, which can be initiated by various cytokines.
  - **Expected Result**: Given that the interaction between MM cells and BMSCs may stimulate cytokine production or response, an up-regulation of SOCS3 could be anticipated as a regulatory response to keep the cytokine signaling in check.
- **JUNB**:
  - **Role**: JUNB is a member of the AP-1 transcription factor complex and is involved in various cellular processes, including cell growth, differentiation, and apoptosis. Its role in the bone marrow environment has been associated with cellular responses to external stimuli, including cytokines.
  - **Expected Result**: The up-regulation of JUNB in MM cells when in coculture with BMSCs might be reflective of altered cellular states or responses to the signaling molecules secreted by BMSCs.

### Conclusion:

It's likely that both SOCS3 and JUNB were highlighted due to their known roles in cytokine signaling and cellular responses in the bone marrow environment, respectively. The up-regulation of these genes suggests a pronounced influence of BMSCs on the MM cell line's transcriptional landscape, which could be in line with existing literature on the role of the bone marrow microenvironment in MM progression. However, to get a full picture, one would need to delve deeper into specific research articles and studies focused on these genes in the context of MM and BMSC interactions.

J

4. For panel C, pick the top two pathways and explain the criteria. It is not indicated on the plot whether upregulated or downregulated genes were used for the pathway enrichment analysis. However, by correlating the findings to existing knowledge in the literature, which group of genes were likely used?

2/2

### Panel C Analysis:

Top Two Pathways:

1. **Cytokine-cytokine receptor interaction**: This pathway has the most significant enrichment, as indicated by the farthest value on the x-axis.
2. **JAK-STAT signaling pathway**: This is the second most enriched pathway, given its position on the x-axis.

Criteria for Selection:

The significance of enrichment for each pathway is represented by the negative logarithm (base 10) of the FDR on the x-axis. A pathway with a higher value on this axis indicates stronger evidence against the null hypothesis (i.e., that the observed gene set overlap occurred by chance), meaning the pathway is more significantly enriched in the dataset. The size of the dot corresponds to the number of genes in the dataset associated with that pathway, while the color indicates the odds ratio.

Inference from Existing Knowledge:

- **Cytokine-cytokine receptor interaction**: This pathway is central to cell signaling, especially in the context of immune responses and inflammation. Given the bone marrow microenvironment's rich cytokine milieu and the interaction between MM cells and BMSCs, one would expect significant changes in genes associated with cytokine signaling.
- **JAK-STAT signaling pathway**: This pathway is directly linked with cytokine signaling. When cytokines bind to their receptors, they often activate the JAK-STAT signaling cascade, leading to the transcriptional activation of target genes. Dysregulation of JAK-STAT signaling is implicated in various diseases, including cancers.

Given that both of these pathways play crucial roles in cell signaling, and considering the transcriptional differences seen in Panel B, it's likely that the up-regulated genes were used for the pathway enrichment analysis. This assumption is further supported by the fact that interactions with BMSCs would typically stimulate MM cells, potentially leading to the up-regulation of genes associated with signaling pathways like those mentioned.

### Conclusion:

The top two pathways from Panel C are central to cytokine signaling and cellular responses. Correlating with existing knowledge, the significant enrichment of these pathways suggests that up-regulated genes in MM cells (when cocultured with BMSCs) were likely used for the pathway enrichment analysis, emphasizing the activated signaling and interactive responses of MM cells in the presence of BMSCs.

J

5. Are there any specific suggestions to improve the data presentation of the figure?

Certainly, while the figure provides a wealth of information, there are some improvements that could be made to enhance clarity and comprehensiveness:

1. **Panel A (PCA Plot)**:
   - **Legends**: Introducing a legend box might make it clearer. While 'T' and 'M' are annotated on the plot, a legend box would give a clear demarcation between coculture and monoculture samples.
   - **Percent Variance**: It could be helpful to add a third principal component (PC3) if there's any additional meaningful variance. This might provide further insights into the dataset's structure.
2. **Panel B (Volcano Plot)**:
   - **Threshold Lines**: Adding horizontal lines to indicate the significance threshold (e.g., FDR < 0.05) and vertical lines for fold change cutoffs (e.g., log2 fold change > 2 or < -2) would provide a clearer visual cue for determining significant up- and down-regulated genes.
   - **More Labels**: While SOCS3 and JUNB are highlighted, it might be informative to label other extreme outliers, which could represent genes of interest.
3. **Panel C (Pathway Enrichment Analysis)**:
   - **Color Scale**: The color scale indicating odd ratios could benefit from a legend or color bar to allow for quantitative interpretation.
   - **Directionality**: To provide clarity on whether the enriched pathways are a result of up-regulated or down-regulated genes, a separate color scheme or symbol (e.g., upward and downward triangles) could be utilized.
   - **Full Pathway Names**: Ensure that pathway names aren't cut off or overlapped. This might involve adjusting text size, plot dimensions, or text positioning.
4. **General Suggestions**:
   - **Font Consistency**: Ensure consistent font sizes and types across all panels.
   - **Uniform Color Palette**: Adopt a consistent color scheme across panels for 'T' and 'M' conditions to avoid confusion.
   - **Figure Title**: A title for the entire figure would offer immediate context to readers.
   - **Methodological Details**: Beneath the figure, a brief note on the methods (e.g., statistical tests used, software/tools, sample size) could offer readers a quick understanding of the analysis performed.
5. **Additional Plots**:
   - **Heatmap**: If space allows, a heatmap of the top differentially expressed genes could provide a deeper view into the expression patterns between the two conditions.
   - **Histogram or Box Plot**: A representation of the overall distribution of fold changes or p-values might help readers gauge the overall trends in the data.

These suggestions aim to offer a more comprehensive, clearer, and user-friendly presentation of the data while maintaining the scientific integrity of the findings.

J

6. Draft a title and figure legend. Make sure to include details.

### Title:

**Transcriptional Response of Multiple Myeloma Cells to Bone Marrow Stromal Cell Coculture: Differential Gene Expression and Pathway Enrichment Analysis**

### Figure Legend:

**Figure X. Differential gene expression and associated biological pathways in RMPI8226 MM cells under trans-well coculture with BMSCs.**

**A. Principal Component Analysis (PCA):** Plot represents the transcriptional variability between MM cells in monoculture (M, in red) and trans-well coculture with BMSCs (T, in blue). PC1 accounts for 78% of the total variance, while PC2 accounts for 15%.

**B. Volcano Plot of Gene Expression:** Differential expression of genes is plotted with fold change (log2, T/M) on the x-axis and statistical significance (-log10 FDR) on the y-axis. Genes up-regulated in coculture are shown in red, while down-regulated genes are in blue. Notable genes, SOCS3 and JUNB, are highlighted and labeled.

**C. Pathway Enrichment Analysis:** Biological pathways enriched in the differentially expressed gene set. The x-axis indicates the statistical significance of enrichment (-log10 FDR), while the y-axis lists various enriched biological pathways. Dot size corresponds to the number of associated genes in the dataset, and color indicates the odds ratio, with warmer colors suggesting higher likelihoods of the observed gene set overlap.

**Methods:** Gene expression profiles from RMPI8226 MM cells, either in monoculture or coculture with BMSCs, were obtained using RNA-Seq. Differential expression was determined using [specific software/tool], with an FDR-adjusted p-value <0.05 considered statistically significant. Pathway enrichment was conducted using [specific tool/software], focusing on the [up-regulated/down-regulated] gene set.

**Note:** Exact methods and software/tools should be filled in based on the actual methodologies used in the study.

J

7. Draft a paragraph to describe the findings from the figure. This is one paragraph for the result part of a manuscript. 150 words or less.

In our examination of the transcriptional interplay between RMPI8226 MM cells and BMSCs, clear distinctions in gene expression profiles were observed (Figure X). Principal Component Analysis revealed two discrete clusters, highlighting pronounced transcriptional variations between monoculture and trans-well coculture conditions, with the latter capturing a combined variance of 93% (PC1: 78%, PC2: 15%). A subsequent volcano plot elucidated differential gene expression patterns, notably showcasing the up-regulation of genes SOCS3 and JUNB in coculture. Delving deeper, pathway enrichment analysis underscored the pertinence of "Cytokine-cytokine receptor interaction" and "JAK-STAT signaling pathway", both of which exhibited significant enrichment. These pathways, pivotal in cytokine signaling and cellular interactions, suggest a heightened responsiveness of MM cells to the stimulatory environment provided by BMSCs. Collectively, our findings illuminate the intricate transcriptional alterations in MM cells modulated by the presence of BMSCs, spotlighting potential molecular players and pathways in this dynamic interaction.
