## Supplementary File 7 for "Bioinformatics Illustrations Decoded by ChatGPT: The Good, The Bad, and The Ugly"

GPT-4

J

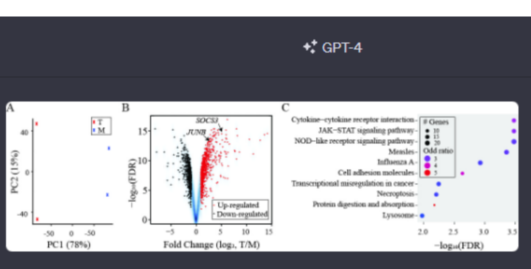


Acting as a bioinformatics professional proficient in RNA-Seq data analysis, you are interpreting a figure to better appreciate the transcription response of multiple myeloma cells (MMs) to bone marrow stromal cells (BMSCs). You also have extensive laboratory research experience in the bone marrow tumor microenvironment of MM. In this task, the expression profile of MM cell line RMPI8226 in trans-well coculture (T) with a BMSC cell line was compared to monoculture (M). The attached figure illustrates the differential expression between the two conditions. Please do the following:

1. Provide an overview of each panel.

Of course, let's break down the attached figure panel by panel.

### Panel A:

This panel depicts a principal component analysis (PCA) plot. PCA is a dimension-reduction tool that is commonly used in RNA-Seq analysis to visualize the overall variance in gene expression data.

- **X-axis (PC1, 78%):** Represents the first principal component which captures the highest variance (78%) in the dataset.
- **Y-axis (PC2, 15%):** Represents the second principal component which captures the second highest variance (15%) in the dataset.
- **Blue Crosses (M):** Represent the monoculture condition (M) of the RMPI8226 MM cell line.
- **Red Stars (T):** Represent the trans-well coculture condition (T) with the BMSC cell line.

The distance between the points provides an indication of the difference in gene expression profiles. Here, the two conditions (M and T) form distinct clusters, suggesting a significant difference in their gene expression profiles.

### Panel B:

This is a volcano plot, a commonly used visualization in differential gene expression analysis.

- **X-axis (Fold Change (log2, T/M)):** Represents the log2 transformed fold change in gene expression between the trans-well coculture (T) and monoculture (M) conditions. A value >0 indicates up-regulation in the T condition, while a value <0 indicates down-regulation.
- **Y-axis (-log10(FDR)):** Represents the -log10 transformed False Discovery Rate (FDR). Higher values indicate greater significance in differential expression.
- **Blue Dots:** Represent down-regulated genes in the T condition as compared to M.
- **Red Dots:** Represent up-regulated genes in the T condition as compared to M.
- **Black Dots:** Represent genes that are not significantly differentially expressed.
- **Highlighted Genes (e.g., SOCS3, JUNB):** These are specific genes that have been labeled, probably because they have a known biological significance or are of particular interest in the study.

### Panel C:

This panel represents a pathway enrichment analysis, which identifies biological pathways that are over-represented in the set of differentially expressed genes.

- **X-axis (-log10(FDR)):** Represents the -log10 transformed False Discovery Rate (FDR) for the enrichment of each pathway. More significant pathways (lower FDR values) are further to the right.
- **Y-axis (Pathways):** Lists various biological pathways.
- **Dot Size (# Genes):** The size of the dots indicates the number of genes from the differential expression analysis that fall into each pathway. Bigger dots mean more genes.
- **Dot Color (Odd ratio):** The color of the dots represents the odds ratio, a measure of the strength of association between the presence of differential expression and each pathway. Higher odds ratios are represented with deeper colors, indicating that the pathway is more likely to be affected by the changes in gene expression between T and M conditions.

From the overview, it's evident that interaction with BMSCs in the bone marrow microenvironment leads to notable changes in gene expression in MM cells, affecting various pathways that might be critical in MM pathology or progression.

J

2. For panel A, what do the numbers in parentheses mean?

In Panel A, the numbers in parentheses next to PC1 and PC2 represent the percentage of variance explained by the respective principal components.

- **PC1 (78%):** The first principal component (PC1) captures 78% of the total variance in the dataset. This means that PC1 alone accounts for 78% of the observed differences in gene expression between the samples.
- **PC2 (15%):** The second principal component (PC2) captures 15% of the total variance in the dataset.

Together, PC1 and PC2 account for 93% (78% + 15%) of the total variability in the data. These percentages help researchers understand how well the reduced dimensionality (in this case, two dimensions) represents the original high-dimensional data (i.e., expression levels of thousands of genes). In this case, a large portion of the variability in the data is captured by just these two components, making the PCA plot a meaningful visualization of the differences between the samples.

J

3. For panel B, the positions of SOCS3 and JUNB are indicated by arrowheads. Estimate the values from the x-axis for the two genes. Are the two genes up-regulated or down-regulated? Based on your findings in the literature, why are the two genes chosen and are these results expected?

### Panel B Analysis:

1. **Estimation of Values for SOCS3 and JUNB**:
   - **SOCS3:** Appears to be located at approximately a value of 5 or slightly more on the x-axis.
   - **JUNB:** Seems to be situated around the value of 7 or slightly above on the x-axis.
2. **Up-regulated or Down-regulated**:
   - Since the x-axis represents the log2-transformed fold change in gene expression between the trans-well coculture (T) and monoculture (M) conditions, values greater than 0 indicate up-regulation in the T condition. Given that both SOCS3 and JUNB have positive values on the x-axis, we can conclude that both genes are **up-regulated** in the coculture condition (T) compared to the monoculture (M).

### Literature Insights on SOCS3 and JUNB:

- **SOCS3 (Suppressor of Cytokine Signaling 3):**
  - Function: It is known to be a negative regulator of cytokine signaling and can inhibit the JAK-STAT signaling pathway, which is critical in cytokine-mediated cellular responses.
  - Relevance to MM: The bone marrow microenvironment in MM is rich in cytokines, which play a role in promoting MM cell growth, survival, and drug resistance. An up-regulation of SOCS3 might be an attempt of MM cells to regulate or counteract excessive cytokine signaling.
  - Expected Results: The up-regulation of SOCS3 in the presence of BMSCs might be anticipated given the cytokine-rich environment and the role of SOCS3 in negative feedback regulation of cytokine responses.
- **JUNB (Jun B proto-oncogene):**
  - Function: JUNB is a component of the AP-1 transcription factor complex and is involved in a variety of cellular processes, including proliferation, differentiation, and apoptosis.
  - Relevance to MM: Changes in JUNB expression can influence the behavior of cancer cells, including MM. In some contexts, JUNB acts as a tumor suppressor, while in others, it may have oncogenic roles.
  - Expected Results: Given the multifaceted role of JUNB in cancer biology, its up-regulation in the presence of BMSCs suggests that BMSCs might be influencing MM cell behavior, possibly driving proliferation or altering differentiation status. The exact role would need to be validated with further experiments, but its selection indicates its potential importance in the MM and BMSC interaction.

In conclusion, both genes, SOCS3 and JUNB, have been chosen for their known roles in cellular signaling and cancer biology. Their up-regulation in the coculture condition suggests they might be key players in the interaction between MM cells and BMSCs. The observed results are consistent with what might be expected given the roles of these genes in similar cellular contexts.

J

4. For panel C, pick the top two pathways and explain the criteria. It is not indicated on the plot whether upregulated or downregulated genes were used for the pathway enrichment analysis. However, by correlating the findings to existing knowledge in the literature, which group of genes were likely used?

### Panel C Analysis:

**Top Two Pathways**:

1. **Cytokine-Cytokine Receptor Interaction:** This pathway is the top-most one on the y-axis. Its significance is given by its position on the x-axis, which has a very low FDR (as indicated by the -log10(FDR) value being farthest to the right).
2. **JAK-STAT Signaling Pathway:** This is the second pathway listed. Like the first pathway, it also shows a significant enrichment, as indicated by its position on the x-axis, though slightly less significant than the cytokine-cytokine receptor interaction.

### Criteria for Selecting the Top Two Pathways:

1. **Significance:** Based on the -log10(FDR) value on the x-axis, which measures the statistical significance of the enrichment of differentially expressed genes in a particular pathway. The farther to the right a dot is, the more significant the pathway enrichment.
2. **Number of Genes:** The size of the dot represents the number of differentially expressed genes that are associated with that pathway. Larger dots indicate more genes, meaning that pathway is highly represented in the list of differentially expressed genes.

### Correlation with Existing Knowledge:

**Cytokine-Cytokine Receptor Interaction and JAK-STAT Signaling Pathway** are closely linked. The JAK-STAT pathway is activated by cytokines and is essential for transmitting information from chemical signals outside the cell to the cell nucleus. Given that both pathways are significant, this suggests a high level of activity related to cytokine signaling in the coculture of MM cells with BMSCs.

Given the nature of these pathways:

1. **Cytokine-Cytokine Receptor Interaction:** It deals with interactions between different cytokines and their receptors. Cytokines are signaling molecules that play a significant role in the bone marrow microenvironment, influencing cell growth, differentiation, and immune responses.
2. **JAK-STAT Signaling Pathway:** A central pathway mediating the effects of cytokines. In the context of MM, the JAK-STAT pathway has been implicated in MM cell proliferation, survival, and resistance to apoptosis.

Considering the emphasis of these pathways and their role in cell signaling, growth, and survival, it is likely that the **up-regulated genes** were used for the pathway enrichment analysis. This is because the interaction of MM cells with BMSCs would be expected to stimulate or activate signaling pathways that promote MM cell survival and growth in the bone marrow microenvironment. Thus, up-regulated genes in the coculture condition (as opposed to down-regulated ones) would be more indicative of the pathways that are being actively engaged in response to BMSC interaction.

J

5. Are there any specific suggestions to improve the data presentation of the figure?

Certainly, the presentation of scientific data is crucial for clear interpretation and communication. Here are some suggestions to improve the data presentation of the figure:

1. **Panel A (PCA Plot):**
   - **Legend Expansion:** The symbols for T and M might be obvious to those familiar with the experiment, but it would be beneficial to have a more detailed legend indicating what "T" and "M" stand for.
   - **Axes Labels:** It might be helpful to provide more descriptive labels for the axes, such as "Principal Component 1 (78% Variance Explained)" instead of just "PC1 (78%)".
2. **Panel B (Volcano Plot):**
   - **Threshold Lines:** Add horizontal dashed lines to indicate the significance threshold (commonly an FDR or p-value threshold). Similarly, vertical dashed lines can be added to indicate fold-change thresholds that were used to determine significance.
   - **Color Gradient or Size Gradient for Significance:** Instead of just three colors, a gradient could represent the range of significance, giving a better idea of the genes that are just on the cusp of significance versus those that are highly significant.
   - **More Highlighted Genes:** If there are other genes of importance or relevance to the study, they could also be labeled, not just SOCS3 and JUNB.
3. **Panel C (Pathway Enrichment Analysis):**
   - **Bar Plot Option:** Instead of or in addition to the dot plot, a horizontal bar plot could be used to represent the -log10(FDR) values, making it easier to visually compare the significance of different pathways.
   - **Direction of Regulation:** It might be helpful to use different symbols or colors for pathways enriched in up-regulated versus down-regulated genes. This would give the reader a quick understanding of whether the pathway is activated or inhibited in the coculture condition.
   - **Full Descriptions:** Ensure that all pathway names are fully visible and not cut off or overlapped.
4. **General Suggestions:**
   - **Consistent Color Palette:** Ensure that colors representing the same conditions (T and M or up-regulated and down-regulated) are consistent across all panels.
   - **Figure Title:** Adding a concise title to the entire figure can provide immediate context to the viewer.
   - **Font Size & Clarity:** Ensure that all labels, legends, and annotations are of a readable size and clear font.
   - **Methods Reference:** If space allows, a brief note or reference pointing to the methods section for more details on the data processing and analysis can be helpful.

Remember, the goal is to make the figure as informative and clear as possible, minimizing any potential confusion for readers who may not be intimately familiar with the experimental details or data analysis methods.

J

6. Draft a title and figure legend. Make sure to include details.

Certainly! Here's a draft title and figure legend for the provided figure:

---

**Title:**
**Differential Gene Expression and Pathway Enrichment Analysis of Multiple Myeloma Cells Co-cultured with Bone Marrow Stromal Cells**

---

**Figure Legend:**

**Panel A:** Principal Component Analysis (PCA) of gene expression profiles. The plot depicts the first two principal components, PC1 and PC2, capturing 78% and 15% of the variance, respectively. Red stars represent MM cell line RMPI8226 in trans-well coculture with BMSCs (T), while blue crosses depict the monoculture condition (M).

**Panel B:** Volcano plot illustrating the differential gene expression between the coculture (T) and monoculture (M) conditions. The x-axis shows the log2-transformed fold change (T/M), while the y-axis presents the -log10 transformed False Discovery Rate (FDR). Red dots denote up-regulated genes, blue dots signify down-regulated genes, and black dots represent genes with no significant differential expression. Arrowheads highlight two genes of interest, SOCS3 and JUNB.

**Panel C:** Pathway enrichment analysis of differentially expressed genes. Pathways are listed on the y-axis, and their enrichment significance is represented on the x-axis as -log10(FDR). Dot size corresponds to the number of genes from the differential expression analysis associated with each pathway, while dot color indicates the odds ratio of the association.

---

This title and legend aim to provide a clear and concise summary of the figure while detailing the methods and findings. It can be adjusted based on any additional specifics or nuances of the study.

J

7. Draft a paragraph to describe the findings from the figure. This is one paragraph for the result part of a manuscript. 150 words or less.

In our investigation of the transcriptional response of multiple myeloma (MM) cells to bone marrow stromal cells (BMSCs), distinct gene expression profiles were observed between MM cells cocultured with BMSCs and those in monoculture, as evidenced by the pronounced separation in the PCA plot (Panel A). Notably, genes such as SOCS3 and JUNB were significantly up-regulated in the coculture condition (Panel B). Pathway enrichment analysis further illuminated the biological implications of these transcriptional changes (Panel C). The most pronounced pathways were the "Cytokine-Cytokine Receptor Interaction" and the "JAK-STAT Signaling Pathway," both pivotal in mediating cellular responses in the tumor microenvironment. The prominence of these pathways, especially in the context of cytokine signaling, underlines the influential role of the BMSCs in shaping the MM cellular response and possibly driving disease progression or therapeutic resistance. This interplay between MM cells and their bone marrow milieu is crucial for understanding disease dynamics and potential therapeutic targets.
