## Supplementary File 8 for "Bioinformatics Illustrations Decoded by ChatGPT: The Good, The Bad, and The Ugly"

GPT-4

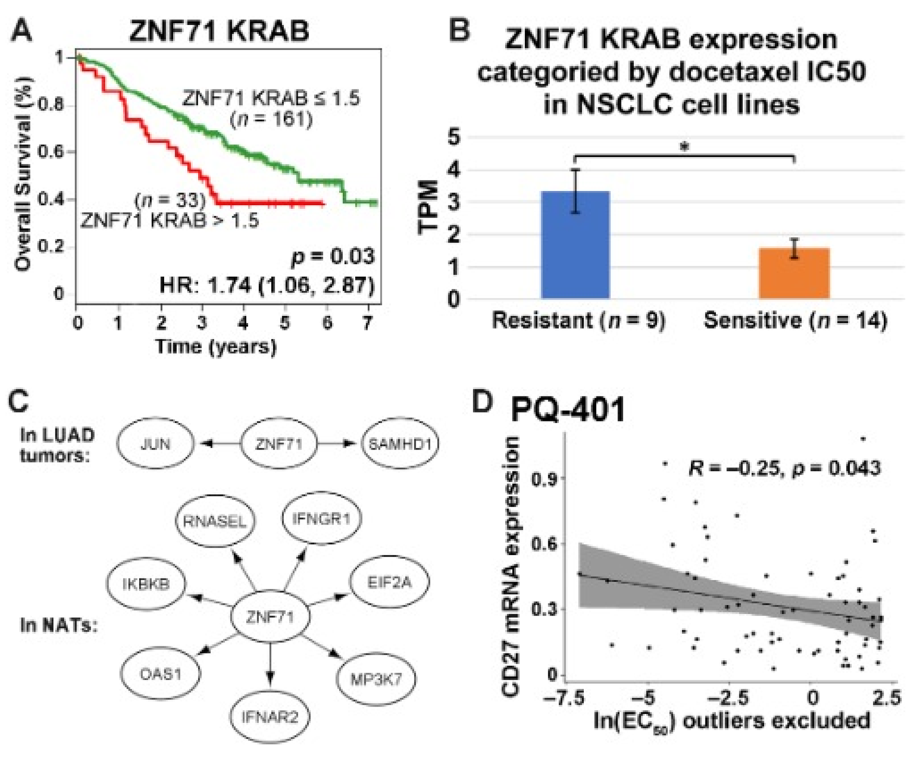


Acting as a bioinformatics professional. Please provide an overview on each panel.

Certainly! Let's break down the information presented in each panel:

**Panel A: Kaplan-Meier Survival Curve**

- This graph represents the overall survival rates of patients over a period of time based on the expression of the ZNF71 KRAB gene.
- Two groups are presented:
  1. Patients with ZNF71 KRAB expression ≤ 1.5 (depicted in red) with a total of 161 patients.
  2. Patients with ZNF71 KRAB expression > 1.5 (depicted in green) with a total of 33 patients.
- The Hazard Ratio (HR) is 1.74 with a 95% confidence interval of 1.06 to 2.87, which suggests that there might be a statistically significant difference in survival between the two groups.
- The p-value is 0.03, which indicates that the difference in survival between the two groups is statistically significant at a 0.05 significance level.

**Panel B: Bar Chart of ZNF71 KRAB Expression by Docetaxel Sensitivity**

- This graph shows the ZNF71 KRAB gene expression (in TPM - Transcripts Per Million) in NSCLC (Non-Small Cell Lung Cancer) cell lines categorized by their sensitivity to the drug docetaxel.
- Two groups are presented:
  1. Resistant cell lines (blue bar) with a total of 9 samples.
  2. Sensitive cell lines (orange bar) with a total of 14 samples.
- There's a statistically significant difference (indicated by the asterisk) between the two groups, suggesting that ZNF71 KRAB expression might be related to docetaxel sensitivity.

**Panel C: Gene Interaction Diagram**

- This depicts the interactions of the ZNF71 gene with other genes in LUAD (Lung Adenocarcinoma) tumors and in NATs (presumably Normal Adjacent Tissues).
- In LUAD tumors, ZNF71 interacts with genes like JUN, SAMHD1, etc.
- In NATs, ZNF71 interacts with different genes such as RNASEL, IFNGR1, and others.

**Panel D: Scatter Plot of PQ-401 vs. CD271 mRNA Expression**

- The graph represents the relationship between the natural logarithm (ln) of the effective concentration at 50% (EC50) for PQ-401 (excluding outliers) and the mRNA expression levels of CD271.
- The correlation coefficient, R, is -0.25, which suggests a weak negative correlation between the two variables.
- The shaded region indicates the confidence interval for the regression line.
- The p-value of 0.043 indicates that the correlation is statistically significant at the 0.05 level.

In summary, these panels seem to provide evidence on the significance of ZNF71 KRAB in relation to patient survival, drug sensitivity in NSCLC cell lines, gene interactions in different tissue types, and its potential relation with CD271 expression in the context of PQ-401.

In Panel A: Is ZNF71 KRAB favorable or unfavorable? Why?

In Panel A, the Kaplan-Meier Survival Curve compares the overall survival rates of patients based on the expression of the ZNF71 KRAB gene.

From the graph:

- The red curve represents patients with ZNF71 KRAB expression ≤ 1.5.
- The green curve represents patients with ZNF71 KRAB expression > 1.5.

By examining the survival curves:

- Patients with a ZNF71 KRAB expression ≤ 1.5 (red curve) have a better survival rate over time than those with ZNF71 KRAB expression > 1.5 (green curve).

Given this information, **ZNF71 KRAB expression at levels greater than 1.5 appears to be unfavorable for patient survival**. This is because patients with higher expression of this gene have a lower survival rate over time compared to those with lower expression.

In Panel B: Is ZNF71 KRAB resistant or sensitive to docetaxel? Why and be quantitative?

In Panel B, the bar chart displays the expression levels of ZNF71 KRAB in NSCLC (Non-Small Cell Lung Cancer) cell lines based on their sensitivity to docetaxel.

From the graph:

- The blue bar represents cell lines that are resistant to docetaxel.
- The orange bar represents cell lines that are sensitive to docetaxel.

Quantitatively:

- Resistant cell lines (blue bar) show a higher ZNF71 KRAB expression, roughly around 4 TPM (Transcripts Per Million).
- Sensitive cell lines (orange bar) have a lower ZNF71 KRAB expression, approximately near 2 TPM.

Given this quantitative data, **cell lines with higher expression levels of ZNF71 KRAB (around 4 TPM) are resistant to docetaxel**, while those with lower expression levels (around 2 TPM) are more sensitive to the drug. Thus, ZNF71 KRAB can be considered as associated with resistance to docetaxel in these NSCLC cell lines.

In Panel C: Which genes are associated with ZNF71 in LUAD tumors? Which genes are associated with ZNF71 in non-cancerous adjacent tissues?

In Panel C, the gene interaction diagram illustrates the associations of the ZNF71 gene with other genes in different tissue types.

**For LUAD (Lung Adenocarcinoma) tumors:**
ZNF71 is associated with the following genes:

- JUN
- SAMHD1

**For NATs (presumably Normal Adjacent Tissues):**
ZNF71 is associated with the following genes:

- RNASEL
- IFNGR1
- IKKBK
- EIF2A
- OAS1
- MP3K7
- IFNAR2

These associations indicate that the ZNF71 gene might interact differently in tumorous vs. non-tumorous tissues, suggesting potential roles in tumor biology and possibly in normal cellular functions.

In Panel D: What is the R value in Panel C? What is the indication on the expression of CD27 and PQ-401 resistance?

In Panel D, the scatter plot represents the relationship between the natural logarithm (ln) of the effective concentration at 50% (EC50) for PQ-401 and the mRNA expression levels of CD271.

The given R value in Panel D is:
**R = -0.25**

This R value indicates a weak negative correlation between the two variables. In the context of this scatter plot:

- A negative R value means that as the expression of CD271 increases, the ln(EC50) for PQ-401 decreases (or vice versa).
- Since EC50 represents the concentration of the drug required to achieve 50% of its maximum effect, a lower ln(EC50) suggests increased sensitivity to the drug (i.e., less drug is needed to achieve the desired effect), while a higher ln(EC50) suggests resistance.

Given the negative correlation:

- **Higher expression levels of CD271 are associated with increased sensitivity to PQ-401**, whereas lower expression levels of CD271 might be linked to resistance to PQ-401. However, it's essential to note that the correlation is weak, so while there is a trend, it might not be a very strong predictor on its own. Further studies would be beneficial to confirm this association.
