## Supplementary File 9 for "Bioinformatics Illustrations Decoded by ChatGPT: The Good, The Bad, and The Ugly"

GPT-4

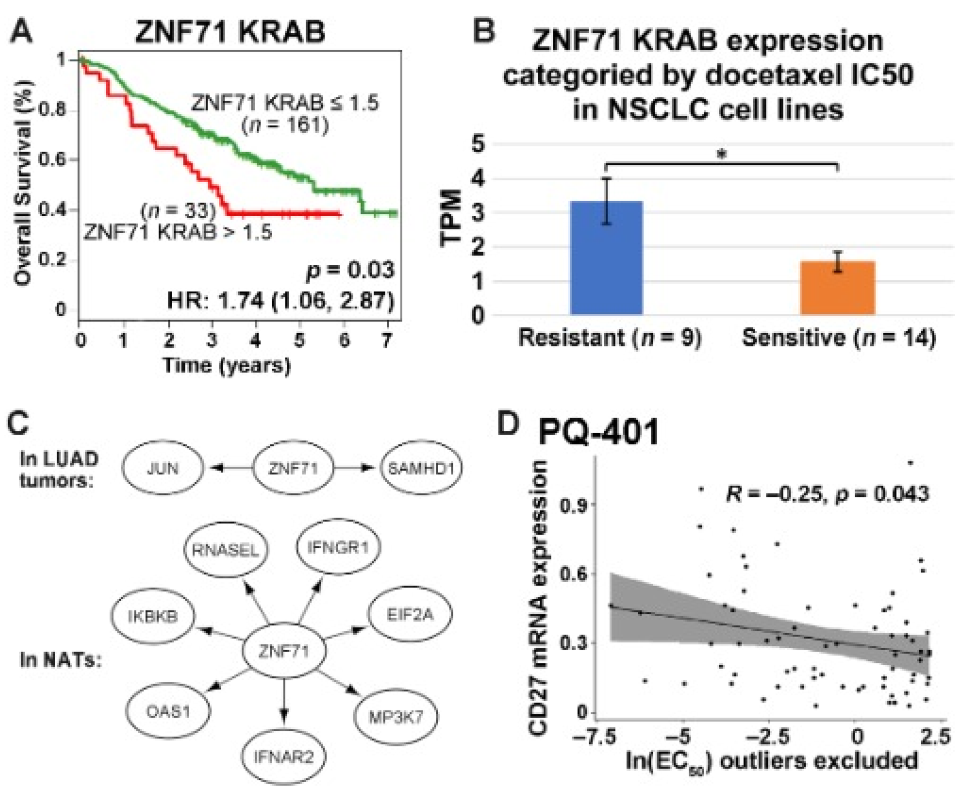


Acting as a bioinformatics professional. Please provide an overview on each panel.

Certainly! Here's an overview of each panel in the image:

**Panel A: Kaplan-Meier Survival Curve**

- The Y-axis represents the overall survival rate of patients, and the X-axis represents the time in years.
- Two survival curves are shown:
  1. Red curve: Represents patients with ZNF71 KRAB expression ≤1.5. The number of patients in this group is 161.
  2. Green curve: Represents patients with ZNF71 KRAB expression >1.5. The number of patients in this group is 33.
- The survival curve shows that patients with ZNF71 KRAB expression >1.5 have a lower survival rate over time.
- The hazard ratio (HR) for this comparison is 1.74 with a 95% confidence interval of 1.06 to 2.87. The p-value for this difference is 0.03, suggesting that the difference in survival between the two groups is statistically significant.

**Panel B: Bar Graph of ZNF71 KRAB Expression and Docetaxel Resistance**

- The Y-axis represents ZNF71 KRAB expression in Transcripts Per Million (TPM).
- Two bars are shown:
  1. Blue bar: Represents NSCLC (Non-Small Cell Lung Cancer) cell lines that are resistant to docetaxel. There are 9 cell lines in this group.
  2. Orange bar: Represents NSCLC cell lines that are sensitive to docetaxel. There are 14 cell lines in this group.
- Cell lines that are resistant to docetaxel have higher ZNF71 KRAB expression compared to sensitive ones. The difference is marked by an asterisk, indicating statistical significance.

**Panel C: Pathway Diagram**

- Displays the interactions and pathways involving the ZNF71 gene in Lung Adenocarcinoma (LUAD) tumors (top part) and in Normal Adjacent Tissues (NATs) (bottom part).
- In LUAD tumors, ZNF71 interacts with JUN and SAMHD1.
- In NATs, ZNF71 has several interactions, including with RNASEL, IFNGR1, IKKB, EIF2A, OAS1, IFNAR2, and MP3K7. The arrows indicate the directionality or influence of these interactions.

**Panel D: Scatter Plot of PQ-401 and CD271 mRNA Expression**

- The Y-axis represents CD271 mRNA expression, and the X-axis represents the natural logarithm of EC50 for PQ-401 (with outliers excluded).
- The scatter plot shows a correlation between the two variables, represented by the trendline.
- The correlation coefficient (R) is -0.25, indicating a negative correlation between CD271 mRNA expression and the EC50 of PQ-401. The p-value for this correlation is 0.043, suggesting that the correlation is statistically significant.

In summary, these panels provide insights into the role and significance of ZNF71 KRAB expression in patient survival, its association with docetaxel resistance in NSCLC cell lines, its interaction pathways in both tumors and adjacent tissues, and its correlation with CD271 mRNA expression in the context of PQ-401 response.

In Panel A: Is ZNF71 KRAB favorable or unfavorable? Why?

In Panel A, ZNF71 KRAB expression is depicted in relation to patient survival over time.

When we compare the two survival curves:

- Patients with ZNF71 KRAB expression ≤1.5 (Red curve) have a higher survival rate over the years than those with ZNF71 KRAB expression >1.5 (Green curve).

Given this observation, ZNF71 KRAB expression levels >1.5 are associated with a lower survival rate, making it **unfavorable** for patients. The reduced survival rate for this group suggests that higher expression of ZNF71 KRAB might be linked to a more aggressive disease or reduced response to treatment. Additionally, the hazard ratio (HR) of 1.74 for this group indicates an increased risk of death compared to the group with lower expression, reinforcing the unfavorable nature of higher ZNF71 KRAB expression.

In Panel B: Is ZNF71 KRAB resistant or sensitive to docetaxel? Why and be quantitative?

In Panel B, the bar graph depicts ZNF71 KRAB expression in NSCLC cell lines categorized by their sensitivity to docetaxel.

From the graph:

- The blue bar represents cell lines that are resistant to docetaxel, and their average ZNF71 KRAB expression is higher, approximately around 4.5 TPM.
- The orange bar represents cell lines that are sensitive to docetaxel, and their average ZNF71 KRAB expression is lower, approximately around 1.5 TPM.

Given this quantitative data, cell lines with higher ZNF71 KRAB expression (around 4.5 TPM) are **resistant** to docetaxel, whereas cell lines with lower ZNF71 KRAB expression (around 1.5 TPM) are **sensitive** to docetaxel. This indicates that higher levels of ZNF71 KRAB expression may confer resistance to the drug docetaxel.

In Panel C: Which genes are associated with ZNF71 in LUAD tumors? Which genes are associated with ZNF71 in non-cancerous adjacent tissues?

In Panel C, the interactions and pathways involving the ZNF71 gene in different tissue types are displayed.

For LUAD tumors:

- The genes associated with ZNF71 are **JUN** and **SAMHD1**.

For Normal Adjacent Tissues (NATs):

- The genes associated with ZNF71 are **RNASEL**, **IFNGR1**, **IKKB**, **EIF2A**, **OAS1**, **IFNAR2**, and **MP3K7**.

In Panel D: What is the R value in Panel C? What is the indication on the expression of CD27 and PQ-401 resistance?

In Panel D:

- The correlation coefficient, $R$, is **-0.25**.

Given this value, there's a negative correlation between CD271 mRNA expression and the natural logarithm of EC50 for PQ-401. This means that as CD271 mRNA expression increases, the EC50 for PQ-401 decreases (and vice versa).

In terms of drug resistance, a lower EC50 value generally indicates that a lower concentration of the drug is required to achieve half-maximal inhibition, suggesting higher sensitivity to the drug. Conversely, a higher EC50 value suggests reduced drug sensitivity or increased resistance.

Given the negative correlation:

- Higher CD271 mRNA expression is associated with greater sensitivity to PQ-401 (or decreased resistance).
- Lower CD271 mRNA expression is associated with reduced sensitivity or increased resistance to PQ-401.
