## Supplementary File 10 for "Bioinformatics Illustrations Decoded by ChatGPT: The Good, The Bad, and The Ugly"

GPT-4

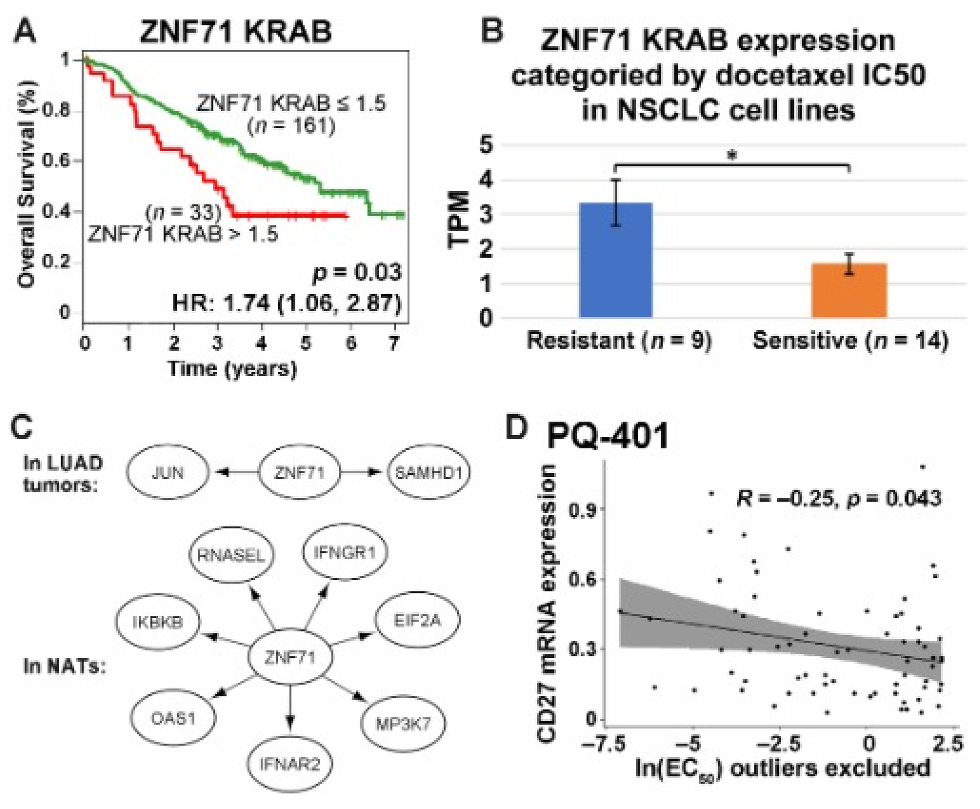


Acting as a bioinformatics professional. Please provide an overview on each panel.

Certainly! Here's a breakdown of each panel:

**Panel A: Survival Analysis**

- This panel depicts the Kaplan-Meier survival curve for patients based on the expression levels of ZNF71 KRAB.
- Patients are divided into two groups: one with ZNF71 KRAB expression levels ≤1.5 (red line) and one with levels >1.5 (green line).
- The x-axis represents time in years, and the y-axis represents the percentage of patients surviving.
- The red group seems to have a better overall survival than the green group.
- The hazard ratio (HR) is 1.74 with a 95% confidence interval of 1.06 to 2.87, suggesting a statistically significant difference in survival between the two groups.
- A p-value of 0.03 indicates this difference is statistically significant.

**Panel B: Expression Analysis in Relation to Drug Sensitivity**

- This bar chart shows the expression levels of ZNF71 KRAB categorized by docetaxel IC50 (a measure of drug sensitivity) in NSCLC (Non-Small Cell Lung Cancer) cell lines.
- The y-axis represents expression levels in TPM (Transcripts Per Million).
- NSCLC cell lines are divided into two groups based on their sensitivity to docetaxel: Resistant (blue bar) and Sensitive (orange bar).
- Resistant cell lines show higher ZNF71 KRAB expression levels compared to sensitive ones.
- The asterisk denotes a significant difference between the two groups.

**Panel C: Gene Interaction Diagram**

- This panel shows a gene interaction network centered around ZNF71.
- For LUAD tumors (Lung Adenocarcinoma): ZNF71 appears to influence JUN and SAMHD1.
- For NATs (potentially normal adjacent tissues): ZNF71 seems to interact or influence multiple genes, including RNASEL, IFNGR1, IKKBK, EIF2A, OAS1, IFNAR2, and MP3K7. The interactions are denoted by arrows.

**Panel D: Correlation Plot**

- This scatter plot represents the relationship between the expression of CD271 mRNA and a drug named PQ-401 (probably an inhibitor) with its ln(EC50) values. Outliers have been excluded for clarity.
- The x-axis represents ln(EC50) values, a logarithmic measure of drug efficacy, and the y-axis represents CD271 mRNA expression levels.
- The trend line indicates a negative correlation (R = -0.25) between CD271 mRNA expression and ln(EC50), with a p-value of 0.043 suggesting this correlation is statistically significant.

In summary, this figure provides information on the role of ZNF71 KRAB in patient survival, its correlation with drug sensitivity in NSCLC cell lines, its interaction with other genes, and the correlation between CD271 mRNA expression and drug efficacy.

In Panel A: Is ZNF71 KRAB favorable or unfavorable? Why?

In Panel A, ZNF71 KRAB appears to be unfavorable for patient survival. The reason for this interpretation is:

- The group of patients with higher expression levels of ZNF71 KRAB (>1.5) has a lower overall survival rate compared to those with lower expression levels (≤1.5). The green survival curve for the higher expression group declines more rapidly than the red curve of the lower expression group.
- The hazard ratio (HR) provided is 1.74 (with a 95% confidence interval of 1.06 to 2.87). An HR greater than 1 suggests that there's a higher risk of the event occurring (in this case, death) in the group with higher expression levels compared to the group with lower expression levels.

The combined observation of the survival curve and the hazard ratio suggests that higher levels of ZNF71 KRAB are associated with decreased overall survival, making it an unfavorable factor in the context presented.

In Panel B: Is ZNF71 KRAB resistant or sensitive to docetaxel? Why and be quantitative?

In Panel B, ZNF71 KRAB is associated with resistance to docetaxel. Here's why, based on the quantitative data provided:

- The expression level of ZNF71 KRAB in the resistant group (blue bar) is higher than in the sensitive group (orange bar).
- Quantitatively, the average expression in TPM for the resistant group is around 4 (or slightly above), while for the sensitive group, it's around 1.5.

This suggests that cell lines with higher expression levels of ZNF71 KRAB are more resistant to docetaxel. The difference in expression levels, specifically around 2.5 TPM or more, indicates a stronger resistance to the drug in those cell lines with higher ZNF71 KRAB expression.

In Panel C: Which genes are associated with ZNF71 in LUAD tumors? Which genes are associated with ZNF71 in non-cancerous adjacent tissues?

In Panel C:

**For LUAD tumors (Lung Adenocarcinoma):**
ZNF71 is associated with the following genes:

- JUN
- SAMHD1

**For NATs (potentially normal adjacent tissues):**
ZNF71 is associated with the following genes:

- RNASEL
- IFNGR1
- IKKBK
- EIF2A
- OAS1
- IFNAR2
- MP3K7

These associations are depicted by the arrows, indicating interactions or influence between ZNF71 and the listed genes in the respective tissues.

In Panel D: What is the R value in Panel C? What is the indication on the expression of CD27 and PQ-401 resistance?

In Panel D:

- The $R$ value, which represents the correlation coefficient, is $R = -0.25$.
- The negative $R$ value indicates a negative correlation between CD271 mRNA expression and ln(EC50) for PQ-401. This means that as the expression of CD271 mRNA increases, the ln(EC50) value decreases, suggesting an increased resistance to PQ-401. Conversely, lower CD271 mRNA expression correlates with higher sensitivity to PQ-401.

In simpler terms, higher expression of CD271 is associated with increased resistance to PQ-401, while lower expression of CD271 is associated with increased sensitivity to the drug.
