## Supplementary File 11 for "Bioinformatics Illustrations Decoded by ChatGPT: The Good, The Bad, and The Ugly"

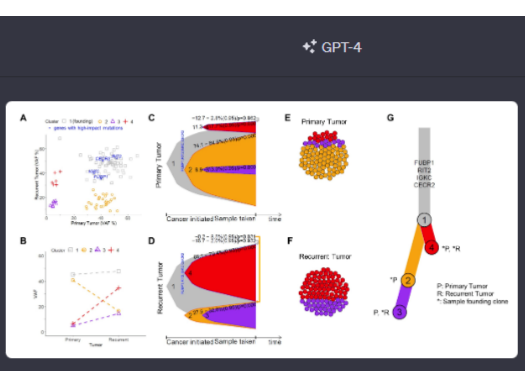


Acting as a bioinformatics professional proficient in cancer genomics data analysis, you will help interpret a figure to better appreciate clonal evolution in multiple myeloma (MM). The attached figure illustrates the results of analyzing somatic mutations detected in a pair of primary and recurrent tumor samples from a patient diagnosed with MM. Please do the following:

1. Provide an overview of each panel. Include the type of plot in the description, e.g., scatter plot, bar plot, etc.

Certainly! Here's an interpretation of the figure's panels:

**Panel A:**

- **Type of plot:** Scatter plot
- **Description:** This plot compares the Variant Allele Frequency (VAF) of mutations in the primary tumor (x-axis) with their VAF in the recurrent tumor (y-axis). Each dot represents a mutation, color-coded by the clone they belong to. Notably, genes with high-impact mutations are highlighted with their respective gene names (e.g., CECR2, RIT2). The plot provides insight into how the prevalence of specific mutations changes between the primary and recurrent tumor stages.

**Panel B:**

- **Type of plot:** Scatter plot with trend lines
- **Description:** This plot showcases the VAF of mutations in primary and recurrent tumors. Different mutation clusters are represented by distinct symbols and color codes. The trend lines illustrate the trajectory of VAF changes from the primary to recurrent stages for each mutation cluster.

**Panel C & D:**

- **Type of plot:** Area plots
- **Description:** These plots illustrate the clonal composition and their respective expansions or contractions in primary and recurrent tumor stages. The x-axis represents a hypothetical timeline from when the cancer was initiated to when the sample was taken. The y-axis could represent the proportion of each clone in the tumor. The p-values suggest significant changes in clonal compositions between the primary and recurrent tumors.

**Panel E & F:**

- **Type of plot:** Cell diagrams
- **Description:** These diagrams visually represent the clonal composition of the primary and recurrent tumors. Each circle represents a cell, and the color indicates its clonal affiliation. This gives a tangible sense of how dominant each clone is within the tumor at different stages.

**Panel G:**

- **Type of plot:** Clonal tree diagram
- **Description:** This is a schematic representation of clonal evolution from the primary to the recurrent tumor. [illegible]nes are depicted as circles, with their size potentially representing their prevalence. Arrows and branching indicate the emergence and evolution of new clones from their predecessors. Gene names listed are probably driver mutations or key mutations that characterize each clone. The annotations "P," "R," and "*" give information about which clones were present in the primary tumor, recurrent tumor, and the founding clone, respectively.

Together, these panels offer a comprehensive view of how tumor composition evolves from the primary to the recurrent stage in this MM patient, highlighting the complexity of tumor heterogeneity and clonal dynamics.

2. Panel A. Identify the number of clusters and their colors. Sort the clusters by size.

In **Panel A**:

**Clusters and Their Colors:**

1. **Cluster 1 (founding)**: Color - Purple
2. **Cluster 2**: Color - Yellow
3. **Cluster 3**: Color - Blue
4. **Cluster 4**: Color - Red

**Sorted by Size**:
By visually examining the scatter plot, we can roughly estimate the number of mutations in each cluster and sort them by size:

1. **Cluster 2 (Yellow)**: This cluster appears to have the most mutations, as seen by the dense collection of yellow dots.
2. **Cluster 3 (Blue)**: Following the yellow cluster, the blue cluster seems to have the second-largest number of mutations.
3. **Cluster 4 (Red)**: This cluster has fewer mutations compared to the yellow and blue ones but more than the purple cluster.
4. **Cluster 1 (founding, Purple)**: This cluster has the least number of mutations, as indicated by the limited number of purple dots.

This order provides an understanding of the prevalence of mutations across different clusters in the tumors.

3. Panel B. Identify clusters showing an increase in VAF from primary to recurrent.

In **Panel B**:

To identify clusters showing an increase in Variant Allele Frequency (VAF) from the primary to the recurrent tumor, we should focus on the trend lines (or trajectories) of each cluster.

From the plot:

1. **Cluster 4 (Red trend line)**: This cluster displays a clear upward trajectory from the primary to the recurrent tumor. The mutations that belong to this cluster show higher VAF values in the recurrent tumor compared to the primary tumor.
2. **Cluster 3 (Purple trend line)**: Though it starts with a high VAF in the primary tumor, there's a subtle increase as we move to the recurrent tumor. This indicates an increase in VAF for this cluster as well.

Clusters 1 and 2, on the other hand, show either a decrease or relatively stable VAF from primary to recurrent.

In conclusion, Clusters 3 and 4 show an increase in VAF from the primary to the recurrent tumor.

4. Panels C and D. Identify the major cluster at the time of sample taken for each panel.

For **Panels C and D**, the major cluster at the time of sample taken can be identified by observing the area plots at the "Sample taken" marker on the x-axis:

**Panel C (Primary Tumor):**

- The largest section of the plot at the "Sample taken" marker is the **orange color**, which corresponds to **Cluster 2**. Therefore, Cluster 2 is the major cluster at the time the primary tumor sample was taken.

**Panel D (Recurrent Tumor):**

- The dominant section of the plot at the "Sample taken" marker is the **red color**, which corresponds to **Cluster 4**. This means that Cluster 4 is the major cluster at the time the recurrent tumor sample was taken.

In summary:

- For the primary tumor (Panel C), Cluster 2 is the major cluster.
- For the recurrent tumor (Panel D), Cluster 4 is the major cluster.

5. Panels E and F. Sort the clusters by size for each panel.

In **Panels E and F**, we can estimate the cluster sizes based on the number of circles (cells) representing each color.

**Panel E (Primary Tumor):**

1. **Yellow (Cluster 2)**: The dominant color with the most circles.
2. **Red (Cluster 4)**: Fewer circles than yellow but more than the other two colors.
3. **Blue (Cluster 3)**: Even fewer circles, making it the third most prevalent.
4. **Purple (Cluster 1)**: The least number of circles in this panel, making it the smallest cluster.

**Panel F (Recurrent Tumor):**

1. **Red (Cluster 4)**: Now the dominant color in this panel with the most circles.
2. **Purple (Cluster 1)**: Significantly increased from the primary tumor and is now the second most prevalent.
3. **Blue (Cluster 3)**: Has fewer circles compared to red and purple, making it the third in size.
4. **Yellow (Cluster 2)**: Drastically reduced in size from the primary tumor and is now the smallest cluster.

In summary:

- For the primary

tumor (Panel E): The order by size is Cluster 2, Cluster 4, Cluster 3, and then Cluster 1.

- For the recurrent tumor (Panel F): The order by size is Cluster 4, Cluster 1, Cluster 3, and then Cluster 2.

6. Panel G: Identify the cluster at the root and the clusters are the leaves.

In **Panel G**, which represents the evolutionary tree of the clusters:

- **Root Cluster**: The cluster at the base or origin of the tree is represented by the circle labeled **1**, which is in **red color**. This corresponds to **Cluster 1**. The root represents the original or founding clone.
- **Leaf Clusters**: The clusters that are at the tips or ends of the branches are considered the "leaves". In this panel, the leaf clusters are represented by circles labeled **2 (in purple color)**, **3 (in blue color)**, and **4 (in orange color)**. These correspond to **Clusters 2, 3, and 4**, respectively.

In conclusion:

- **Cluster 1** is at the root.
- **Clusters 2, 3, and 4** are the leaves.

7. Are there any specific suggestions to improve the data presentation of the figure?

Certainly! While the figure provides a comprehensive representation of clonal evolution in multiple myeloma, there are areas where clarity and presentation can be improved. Here are some suggestions:

1. **Consistency in Color Representation**: Throughout different panels, it's crucial to maintain consistent coloring for clusters. This reduces confusion. For example, in Panel G, Cluster 4 is represented in orange, whereas in other panels it's shown in red.
2. **Enhance Legend Readability**: A consolidated legend indicating each color with its corresponding cluster number and description could be helpful. This way, the reader doesn't need to jump between panels to decode colors.
3. **Clear Labeling of Axes**: In Panels A and B, ensuring that both axes are clearly labeled with their respective metrics can help. For instance, ensuring "VAF %" is clearly visible.
4. **Simplified Statistical Annotations**: The statistical notations in Panels C and D are comprehensive but can be overwhelming. Simplifying these or providing them in a supplementary note might make the main figure more digestible.
5. **Clarity in Cluster Trajectories**: In Panel B, using arrows to indicate the direction from primary to recurrent could provide a clearer understanding of VAF changes.
6. **Zoom or Insets for Dense Areas**: In Panels A and B, where data points are densely packed, a zoomed-in inset could be beneficial for clarity.
7. **Panel G Improvements**:
   - Clarifying the meaning of '*P' and '*R' directly in the panel or in the legend can help.
   - Using line width to indicate the proportion of cells in each evolutionary path could provide a clearer visualization of dominance.
8. **Improved Panel Titles**: More descriptive panel titles or subtitles could provide a quick understanding of each panel's main takeaways.
9. **Use of 3D or Interactive Plots**: If this is for a digital publication, interactive plots where the user can hover over data points to get more information might enhance understanding.
10. **Summary or Key Takeaways**: A small textbox summarizing the key findings from the figure can aid those unfamiliar with the specifics of clonal evolution.

Incorporating these suggestions can make the figure more accessible, especially to those who might be new to the topic or the specific visualization methods used.

8. Draft a title and figure legend. Make sure to include details.

**Title**:
"Clonal Evolution Dynamics in Multiple Myeloma: Analysis of Somatic Mutations in Primary and Recurrent Tumors"

**Figure Legend**:

**Panel A**: Scatter plot representing the Variant Allele Frequency (VAF %) of somatic mutations in primary vs. recurrent tumors. Each point represents a mutation, colored based on its clonal cluster assignment:

- Cluster 1 (Founding): Purple triangles
- Cluster 2: Yellow circles
- Cluster 3: Blue squares
- Cluster 4: Red crosses

**Panel B**: Line plot displaying the change in VAF from primary to recurrent tumors for each clonal cluster.

- Purple dashed line: Cluster 1
- Yellow solid line: Cluster 2
- Blue dash-dot line: Cluster 3
- Red dotted line: Cluster 4

**Panels C & D**: Area plots illustrating the temporal clonal composition of the primary (C) and recurrent (D) tumors, from cancer initiation to the time the sample was taken. Shades represent the proportion of each cluster at different time points.

**Panels E & F**: Hexagonal plots showcasing the clonal composition of the primary (E) and recurrent (F) tumors. Each hexagon represents a cell, colored by its clonal assignment.

**Panel G**: Evolutionary tree diagram depicting the lineage relationship among the clusters. Circle size at the tree nodes corresponds to the cluster's size. The tree's root (Cluster 1) represents the founding clone, while the tips (Clusters 2, 3, and 4) denote evolved clusters.

- '*P' indicates the sample founding clone in the primary tumor.
- '*R' denotes the sample founding clone in the recurrent tumor.

Note: VAF represents the proportion of sequencing reads supporting the mutant allele, providing insights into clonal prevalence.

9. Draft a paragraph to describe the findings from the figure. This is one paragraph for the result part of a manuscript. 150 words or less.

In the presented analysis of somatic mutations from primary and recurrent multiple myeloma tumors, we observed distinct patterns of clonal evolution. Panel A's scatter plot reveals a divergence in the Variant Allele Frequency (VAF) between the tumor stages, with certain mutations becoming more prevalent in the recurrent tumor. This is corroborated in Panel B, which delineates an increased VAF for Clusters 3 and 4 during recurrence. Temporal clonal composition plots (Panels C & D) suggest that by the time of sampling, Cluster 4 dominated in the primary tumor, whereas Cluster 1 was prevalent in the recurrent tumor. The hexagonal plots (Panels E & F) visualize the clonal makeup, highlighting the prominence of Cluster 2 in the primary and Cluster 4 in the recurrent tumors. Panel G's evolutionary tree underscores the foundational role of Cluster 1, with Clusters 2, 3, and 4 emerging as branches, elucidating the dynamic lineage relationships among the clusters in this MM patient.
