## Supplementary File 12 for "Bioinformatics Illustrations Decoded by ChatGPT: The Good, The Bad, and The Ugly"

GPT-4

J

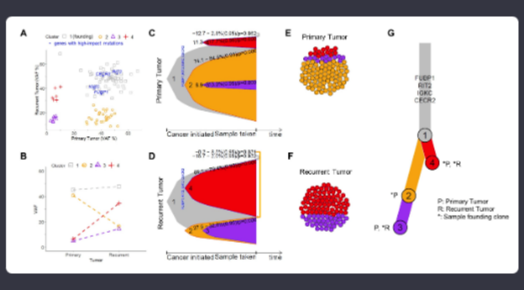


Acting as a bioinformatics professional proficient in cancer genomics data analysis, you will help interpret a figure to better appreciate clonal evolution in multiple myeloma (MM). The attached figure illustrates the results of analyzing somatic mutations detected in a pair of primary and recurrent tumor samples from a patient diagnosed with MM. Please do the following:

1. Provide an overview of each panel. Include the type of plot in the description, e.g., scatter plot, bar plot, etc.

Certainly! Let's dive into the figure and interpret each panel.

**Panel A**

- **Type of Plot:** Scatter plot
- **Description:** This panel displays the variant allele frequency (VAF) of mutations in the primary tumor (on the x-axis) versus the recurrent tumor (on the y-axis). Each point represents a unique mutation. The color and shape of the points represent different genetic clusters. The term "genes with high-impact mutations" indicates that the genes labeled on the plot (e.g., CECR2, RIT2) have mutations that likely have significant functional consequences.

**Panel B**

- **Type of Plot:** Scatter plot with connecting lines
- **Description:** This panel represents the changes in VAF from the primary tumor to the recurrent tumor for specific genetic clusters. The x-axis shows the VAF in the primary tumor and the y-axis for the recurrent tumor. The dotted lines connect the same genetic clusters between the primary and recurrent tumors, indicating the trajectory of mutation frequency changes.

**Panel C & D**

- **Type of Plot:** Area plots
- **Description:** These panels illustrate the evolutionary trajectory of the tumor. The x-axis represents time, starting from when the cancer initiated to when the sample was taken. The colored areas (orange, red, purple) depict the proportion of different genetic clusters over time. The values within the shaded areas indicate percentage change between the primary and recurrent tumors along with statistical significance (p-values).

**Panel E & F**

- **Type of Plot:** Circle packing diagrams
- **Description:** These panels visualize the clonal composition of the primary tumor (E) and recurrent tumor (F). Each circle represents a group of cells with specific mutations. The color indicates different genetic clusters. The size of each circle correlates with the proportion of cells carrying those specific mutations in the tumor.

**Panel G**

- **Type of Plot:** Schematic representation
- **Description:** This panel shows a tree-like structure representing the clonal evolution of the tumor. The trunk represents the founding clone, from which other sub-clones (branches) emerged. The numbers and colors on the tree correspond to the different genetic clusters, while the labels (e.g., FUBP1, RIT2) indicate genes with high-impact mutations. The annotations (*P, *R) suggest which clones were present in the primary and recurrent tumors, respectively.

In summary, this figure provides comprehensive insights into the clonal evolution of a MM patient's tumor from diagnosis (primary) to recurrence. It highlights the genetic diversity, temporal dynamics, and relative abundance of different genetic clones over time.

J

2. Panel A. Identify the number of clusters and their colors. Sort the clusters by size.

From **Panel A**:

**Number of Clusters and Their Colors:**

1. Cluster 1 (founding): Red (`+` symbol)
2. Cluster 2: Yellow (circular symbol)
3. Cluster 3: Purple (triangular symbol)
4. Cluster 4: Light blue (square symbol)

**Sorting Clusters by Size:**
To determine the size of the clusters, we typically look at the number of mutations (points) that belong to each cluster:

1. **Yellow Cluster (Cluster 2)** - This cluster seems to have the highest number of points, making it the largest cluster in terms of mutation count.
2. **Light blue Cluster (Cluster 4)** - This is the next largest cluster based on the number of points.
3. **Red Cluster (Cluster 1)** - This is the founding cluster, and while foundational, it appears to have fewer points than Clusters 2 and 4.
4. **Purple Cluster (Cluster 3)** - This cluster seems to have the fewest points, making it the smallest in terms of mutation count.

So, sorted by size, the clusters are: Yellow (Cluster 2), Light blue (Cluster 4), Red (Cluster 1), and Purple (Cluster 3).

J

3. Panel B. Identify clusters showing an increase in VAF from primary to recurrent.

In **Panel B**:

To determine which clusters show an increase in VAF from primary to recurrent tumors, we need to look at the positioning of the end points of the connecting lines with respect to the diagonal. If the end point for a cluster is above the diagonal, it indicates an increase in VAF from primary to recurrent.

Based on this:

1. **Red Cluster (Cluster 1)**: The line for this cluster moves upward, indicating an increase in VAF from the primary to the recurrent tumor.
2. **Purple Cluster (Cluster 3)**: The line for this cluster also moves upward, showing an increase in VAF from the primary to the recurrent tumor.

Therefore, the clusters showing an increase in VAF from primary to recurrent tumors are the Red (Cluster 1) and Purple (Cluster 3) clusters.

J

4. Panels C and D. Identify the major cluster at the time of sample taken for each panel.

In **Panels C and D**, the major cluster at the time of sample taken can be identified by observing the cluster that occupies the largest area or proportion of the plot at the point labeled "Sample taken."

**Panel C (Primary Tumor):**
The orange shaded region (corresponding to Cluster 4) is the predominant area at the time of sample taken, indicating that Cluster 4 was the major cluster in the primary tumor.

**Panel D (Recurrent Tumor):**
The red shaded region (corresponding to Cluster 1) is the predominant area at the time of sample taken, showing that Cluster 1 was the major cluster in the recurrent tumor.

In summary, for **Panel C**, the major cluster at the time of sample taken is **Cluster 4**, and for **Panel D**, it is **Cluster 1**.

J

5. Panels E and F. Sort the clusters by size for each panel.

Using **Panels E and F**, which employ circle packing diagrams, we can sort the clusters by size based on the total area occupied by each color (or cluster) within the panel.

**Panel E (Primary Tumor):**

1. **Yellow (Cluster 2)**: This color dominates the panel with the most circles, indicating that Cluster 2 is the largest in the primary tumor.
2. **Orange (Cluster 4)**: This is the second largest cluster based on its coverage.
3. **Red (Cluster 1)**: While present, it occupies a smaller portion compared to the Yellow and Orange clusters.
4. **Purple (Cluster 3)**: This seems to be the smallest cluster in terms of size, with only a few circles in the diagram.

**Panel F (Recurrent Tumor):**

1. **Red (Cluster 1)**: In this panel, the red circles dominate, indicating that Cluster 1 is the largest in the recurrent tumor.
2. **Purple (Cluster 3)**: This color is present with significant coverage, making it the second largest cluster.
3. **Orange (Cluster 4)**: Although still present, this cluster occupies a smaller portion compared to Red and Purple clusters.
4. **Yellow (Cluster 2)**: This seems to be the smallest cluster in the recurrent tumor, with very few circles.

In summary, for **Panel E**, the clusters sorted by size are: Yellow (Cluster 2), Orange (Cluster 4), Red (Cluster 1), and Purple (Cluster 3). For **Panel F**, the clusters sorted by size are: Red (Cluster 1), Purple (Cluster 3), Orange (Cluster 4), and Yellow (Cluster 2).

J

6. Panel G: Identify the cluster at the root and the clusters are the leaves.

In **Panel G**, which uses a tree or branch-like representation to illustrate clonal evolution:

**Cluster at the Root:**

- **Red (Cluster 1)**: The base or starting point of the tree is depicted in red, indicating that Cluster 1 is the founding or root cluster.

**Clusters at the Leaves:**

- **Orange (Cluster 4)**: This cluster branches out towards the top, making it one of the leaves.
- **Purple (Cluster 3)**: Another branch that moves outwards, establishing it as a leaf.
- **Yellow (Cluster 2)**: It also branches out from the main stem, indicating it's one of the leaf clusters.

So, **Cluster 1** is at the root, and **Clusters 2, 3, and 4** are the leaves in Panel G.

J

7. Are there any specific suggestions to improve the data presentation of the figure?

Certainly! Here are some suggestions to potentially improve the data presentation of the figure:

1. **Consistency in Color Coding**: While the figure does seem to maintain consistent color coding for clusters across panels, ensuring this consistency can greatly aid in interpretation.
2. **Legends and Labels**:
   - Having a consolidated legend that clearly labels each color to its respective cluster can help reduce confusion.
   - Including clear axis labels and titles can also be beneficial, especially if the figure is viewed outside the context of its original publication.
3. **Clarity in Panels C and D**:
   - The percentages and p-values in Panels C and D are a bit cluttered. Using a larger plot or reducing font size can make it easier to read.
   - Additionally, clearer demarcations or lines at the "Sample taken" point can improve readability.
4. **Use of 3D or Interactive Plots**: If this is for a digital presentation or publication, consider using interactive plots, especially for Panels E and F. This would allow viewers to hover over specific clusters to get more information or to zoom in for a closer look.
5. **Size Representation in Panels E and F**:
   - In the circle packing diagrams, the actual size of individual circles might be confused with the quantity. An accompanying note to clarify that the total area (and not individual circle size) represents cluster size could be beneficial.
6. **Clarify Branch Thickness in Panel G**: If the thickness of the branches in Panel G is indicative of anything (e.g., the number of mutations or the size of the cluster), this should be clearly mentioned.
7. **Descriptive Captions**: While not directly part of the figure, providing a detailed caption that briefly describes each panel can be immensely helpful to readers unfamiliar with the specifics of the data.
8. **Enhance Visual Contrast**: Ensure that there's enough contrast between different colors and the background, making it accessible to all readers, including those with visual impairments.
9. **Annotations**: Brief annotations directly on the plots (especially in Panels C, D, and G) can help highlight key observations or important data points.
10. **Supplementary Materials**: For figures with this level of complexity, it might be useful to have supplementary materials or an appendix that dives deeper into the methodology or offers additional plots for clarity.

Incorporating some or all of these suggestions can make the figure more informative and user-friendly, enhancing its overall impact and clarity.

J

8. Draft a title and figure legend. Make sure to include details.

**Title**:
"Clonal Evolution of Somatic Mutations in Primary and Recurrent Multiple Myeloma"

**Figure Legend**:

**A. Scatter Plot of VAF in Primary vs. Recurrent Tumor**: Each point represents a specific mutation, color-coded based on its respective cluster. The x-axis indicates the Variant Allele Frequency (VAF) in the primary tumor, while the y-axis represents the VAF in the recurrent tumor.

**B. Trajectory of Clusters from Primary to Recurrent Tumor**: This plot tracks the shift in VAF of each mutation cluster from primary to recurrent phase. Each cluster is represented by its unique color and symbol.

**C & D. Temporal Dynamics of Clonal Evolution**: These plots showcase the proportional representation of each mutation cluster over time, from cancer initiation to the point of sample collection. The 'Sample taken' mark indicates when the tumor samples were collected.

- **C** corresponds to the primary tumor and **D** to the recurrent tumor. P-values are provided for significant shifts in proportions.

**E & F. Circle Packing Diagrams of Clusters**:
These diagrams visually represent the relative size of each mutation cluster within the primary (**E**) and recurrent (**F**) tumors. The total area occupied by each color signifies the size of the respective cluster.

**G. Phylogenetic Tree of Clonal Evolution**: Depicting the evolutionary relationship between mutation clusters. The root (red) represents the founding cluster, while the branches culminate in the leaf clusters, indicating subsequent clonal developments.

Note: In all panels, the colors red, yellow, orange, and purple correspond to clusters 1, 2, 4, and 3 respectively. VAF denotes Variant Allele Frequency, an indicator of the proportion of mutated alleles to total alleles in the sample.

J

9. Draft a paragraph to describe the findings from the figure. This is one paragraph for the result part of a manuscript. 150 words or less.

In the study of clonal evolution in multiple myeloma, distinct mutation clusters were identified within primary and recurrent tumor samples from a single patient. As shown in Figure [X], a scatter plot analysis (Panel A) revealed the Variant Allele Frequency (VAF) dynamics between the primary and recurrent tumors, with some clusters demonstrating an increased VAF in the recurrent phase (Panel B). The temporal dynamics (Panels C & D) highlighted the predominant mutation clusters at the time of sample collection. Circle packing diagrams further visualized the relative sizes of these clusters in primary (Panel E) and recurrent (Panel F) tumors. Most notably, a phylogenetic tree (Panel G) delineated the clonal evolutionary trajectory, pinpointing a founding cluster and its subsequent branching into newer clusters. Collectively, these findings illuminate the intricate clonal architecture and evolutionary progression of multiple myeloma, underscoring the dynamic nature of cancer genomics over time.
