## Supplementary File 13 for "Bioinformatics Illustrations Decoded by ChatGPT: The Good, The Bad, and The Ugly"

GPT-4

J

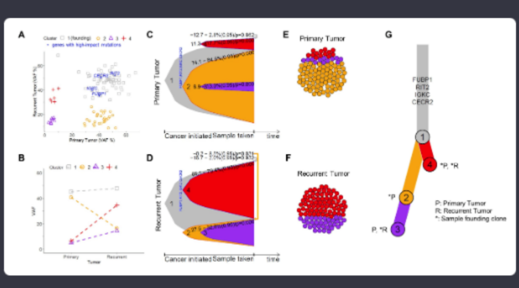


Acting as a bioinformatics professional proficient in cancer genomics data analysis, you will help interpret a figure to better appreciate clonal evolution in multiple myeloma (MM). The attached figure illustrates the results of analyzing somatic mutations detected in a pair of primary and recurrent tumor samples from a patient diagnosed with MM. Please do the following:

1. Provide an overview of each panel. Include the type of plot in the description, e.g., scatter plot, bar plot, etc.

Certainly! Here's a breakdown of each panel in the figure:

**Panel A**

- **Type of Plot**: Scatter plot
- **Overview**: This panel presents the relationship between Variant Allele Frequencies (VAFs) of mutations detected in the primary and recurrent tumors. The x-axis represents the VAF percentages in the primary tumor, while the y-axis denotes the VAF percentages in the recurrent tumor. Different clusters of mutations are color-coded and labeled, with specific genes having high-impact mutations mentioned within the plot.

**Panel B**

- **Type of Plot**: Scatter plot with connected lines
- **Overview**: This panel visually compares the VAF of mutations in the primary tumor to those in the recurrent tumor. Each mutation (or group of mutations) is connected with a dashed line from its primary tumor VAF (x-axis) to its recurrent tumor VAF (y-axis). This kind of plot helps in observing changes in VAF between primary and recurrent tumor stages, indicating potential clonal expansion or reduction.

**Panel C**

- **Type of Plot**: Stacked area plot
- **Overview**: This plot illustrates the trajectory of clonal evolution over time. The x-axis denotes the progression of the disease, from the initiation of the cancer to the point where the sample was taken. The y-axis likely represents the percentage of tumor cells with specific mutations. The colored areas represent different mutation clusters and their prevalence over time. The numerical values and p-values might indicate statistical significance or shifts in the proportions of mutation clusters between primary and recurrent tumors.

**Panel D**

- **Type of Plot**: Stacked area plot
- **Overview**: Similar to Panel C, this plot showcases the clonal evolution trajectory over time for the recurrent tumor. It demonstrates how the proportions of mutation clusters changed from the primary to the recurrent phase.

**Panel E**

- **Type of Plot**: Circle packing diagram (or close packing of circles)
- **Overview**: This diagram represents the primary tumor's cellular composition concerning various mutation clusters. Each circle likely represents a group of cells, with different colors signifying different mutation clusters.

**Panel F**

- **Type of Plot**: Circle packing diagram (or close packing of circles)
- **Overview**: Similar to Panel E, this diagram illustrates the cellular composition of the recurrent tumor concerning different mutation clusters. It shows the changes in cellular composition from the primary to the recurrent phase.

**Panel G**

- **Type of Plot**: Clonal lineage tree
- **Overview**: This tree diagram displays the lineage of mutation clusters, starting with the founding clone and branching out to various derived clones. Labels provide specific genes associated with the mutation clusters. The "*P" and "*R" annotations denote clones present in the primary and recurrent tumors, respectively.

Collectively, these panels provide a comprehensive view of how mutations in the tumor have evolved from the primary to the recurrent phase, offering insights into the clonal dynamics of this MM patient's disease.

J

2. Panel A. Identify the number of clusters and their colors. Sort the clusters by size.

Based on the figure in **Panel A**:

1. **Number of Clusters and Their Colors**:
   - **Cluster 1 (founding)**: Purple triangle
   - **Cluster 2**: Orange circles
   - **Cluster 3**: Red plus signs (+)
   - **Cluster 4**: Blue squares
2. **Sorted by Size**:
   2.1. **Cluster 2** (Orange circles): This seems to be the largest cluster based on the number of data points.
   2.2. **Cluster 3** (Red plus signs): The next in size, with a moderate number of data points.
   2.3. **Cluster 4** (Blue squares): Fewer points compared to Cluster 2 and 3, but still more than the founding cluster.
   2.4. **Cluster 1 (founding)** (Purple triangle): Contains the least number of points, indicating it's the smallest cluster.

Note: This sorting is based on the visible number of data points in Panel A.

J

3. Panel B. Identify clusters showing an increase in VAF from primary to recurrent.

In **Panel B**, clusters that show an increase in VAF from primary to recurrent would be depicted by points that move upwards from the primary (x-axis) to the recurrent (y-axis) tumor VAF.

From the plot:

1. **Cluster 3 (Red plus signs)**: The data points for this cluster seem to show a clear upward trajectory from the primary to the recurrent tumor, suggesting an increase in VAF.
2. **Cluster 4 (Blue squares)**: At least one of the data points for this cluster also seems to move upwards, indicating an increase in VAF for some mutations within this cluster.

The other clusters either remain stable or decrease in VAF from primary to recurrent, based on the observed trajectories of their respective data points.

J

4. Panels C and D. Identify the major cluster at the time of sample taken for each panel.

In **Panels C and D**, the major cluster at the time of sample taken can be identified as the cluster that occupies the largest area or proportion at the far right end of the x-axis (where the sample is taken).

- **Panel C (Primary Tumor)**: The major cluster appears to be the one represented by the **orange** area, which dominates the plot at the time of sample collection. Thus, the major cluster for the primary tumor is the one represented in orange.
- **Panel D (Recurrent Tumor)**: The major cluster in this panel appears to be represented by the **red** area, which dominates the plot at the time of the recurrent sample collection. Hence, the major cluster for the recurrent tumor is the one represented in red.

J

5. Panels E and F. Sort the clusters by size for each panel.

Based on the circle packing diagrams in **Panels E and F**, we can sort the clusters by size by gauging the total area occupied by each color, representing different mutation clusters:

**Panel E (Primary Tumor)**:

1. **Orange Cluster**: This cluster dominates most of the diagram and is the largest.
2. **Red Cluster**: The next largest, though significantly smaller than the orange cluster.
3. **Purple Cluster**: Has fewer circles than the red cluster, making it smaller in size.
4. **Blue Cluster**: Appears to be the smallest, with the least number of circles in the diagram.

**Panel F (Recurrent Tumor)**:

1. **Red Cluster**: This cluster now takes up the most space in the diagram, making it the largest.
2. **Purple Cluster**: Significantly increased from Panel E and now is the second-largest.
3. **Blue Cluster**: Still relatively small, but seems larger than in the primary tumor diagram.
4. **Orange Cluster**: Appears to have shrunk considerably from Panel E and is now the smallest in Panel F.

This indicates dynamic shifts in the clonal populations from the primary to the recurrent tumor stages.

J

6. Panel G: Identify the cluster at the root and the clusters are the leaves.

In **Panel G**, the diagram seems to depict a phylogenetic or clonal evolution tree, where the root represents the founding or original clone, and the leaves represent the evolved or derived clones.

- **Root**: The cluster at the root is represented by the **large circle labeled "1"**, which is colored in red. This indicates the founding or primary clone from which all other clones derived.
- **Leaves**: The clusters that serve as leaves (or terminal nodes) in this evolutionary tree are:
  1. **Circle labeled "2"**, which is colored in purple.
  2. **Circle labeled "3"**, which is also colored in purple.
  3. **Circle labeled "4"**, which is colored in orange.

These leaf clusters represent the most evolved or derived clones from the founding clone, showcasing the clonal heterogeneity and evolutionary trajectory of the tumor.

J

7. Are there any specific suggestions to improve the data presentation of the figure?

Certainly! While the figure is quite informative, there are several suggestions that could potentially improve the data presentation and enhance its clarity:

1. **Consistent Color Coding**: Ensure that the color code remains consistent across all panels. This helps the reader trace the progression of each cluster across different plots without confusion.
2. **Key/Legend**: A unified legend for the entire figure would be beneficial. While some panels have annotations, a legend that details the color, shape, and any other pertinent symbols for each cluster would aid interpretation.
3. **Scale and Axis Labels**: Some panels lack clear axis labels or units. It's essential to label axes clearly in Panels A and B and provide scale where necessary.
4. **Clarity in Panels C and D**: The percentage figures and p-values in Panels C and D are a bit cluttered. Consider using a table or another format to present this numerical data more cleanly.
5. **Enlarge Panels E and F**: These panels are particularly informative but might benefit from being enlarged or given more space for clarity. This would help discern the smaller clusters more easily.
6. **Simplify Panel G**: While the tree diagram in Panel G is informative, it might benefit from some simplification or clearer labeling. For instance, the gray bar in the middle might be made more transparent or labeled to indicate its purpose.
7. **Descriptive Titles for Each Panel**: Brief descriptive titles above each panel would guide the reader on what to expect in each plot, making the figure more intuitive.
8. **Annotate Significant Data Points**: In scatter plots like Panel A and B, consider annotating or highlighting points that are outliers or are of particular significance. This can draw attention to key findings.
9. **Panel Arrangement**: Consider rearranging panels based on the flow of information or the story you want to convey, ensuring that related panels are adjacent or close to each other.
10. **Quality of Text**: Ensure that all text, especially in Panels C and D, is clear, readable, and of high resolution, even when zoomed in.

Incorporating these suggestions could make the figure more reader-friendly, enhancing the understanding and interpretation of the data presented.

J

8. Draft a title and figure legend. Make sure to include details.

**Title**:
"Clonal Evolution and Mutation Dynamics in Primary and Recurrent Multiple Myeloma Tumors"

**Figure Legend**:
**Figure X**: Comprehensive analysis of clonal evolution in a patient with multiple myeloma, from primary to recurrent stages.

- **Panel A**: Scatter plot showing the Variant Allele Frequency (VAF) of mutations in the primary tumor (x-axis) vs. the recurrent tumor (y-axis). Each point represents a mutation, with colors indicating different mutation clusters: Red (Cluster 1 - founding), Orange (Cluster 2), Purple (Cluster 3), and Blue (Cluster 4).
- **Panel B**: Scatter plot illustrating the shift in VAF for each mutation cluster from the primary to the recurrent tumor. Symbols represent mutations within specific clusters: Red plus signs (Cluster 3), Orange circles (Cluster 2), Purple triangles (Cluster 1), and Blue squares (Cluster 4).
- **Panel C**: Temporal clonal evolution diagram for the primary tumor. The width of each colored segment represents the proportion of each mutation cluster at a given time. P-values indicate statistical significance of changes in cluster proportions.
- **Panel D**: Temporal clonal evolution diagram for the recurrent tumor, with a similar representation as in Panel C.
- **Panel E**: Circle packing diagram representing the proportion and distribution of mutation clusters in the primary tumor. Each circle represents a mutation, with colors corresponding to specific clusters.
- **Panel F**: Circle packing diagram for the recurrent tumor, highlighting the shift in mutation clusters from the primary stage.
- **Panel G**: Clonal phylogenetic tree showcasing the evolutionary trajectory of mutation clusters. The root (red circle labeled "1") represents the founding clone, while terminal nodes (circles labeled "2", "3", and "4") represent derived clones.

Note: Colors for mutation clusters are consistent across all panels. Red denotes Cluster 1, Orange represents Cluster 2, Purple is for Cluster 3, and Blue indicates Cluster 4.

J

9. Draft a paragraph to describe the findings from the figure. This is one paragraph for the result part of a manuscript. 150 words or less.

In our examination of clonal evolution in multiple myeloma, the paired analysis of primary and recurrent tumor samples revealed intricate dynamics of clonal shifts. Panel A's scatter plot uncovers a discernible correlation in the Variant Allele Frequency (VAF) between primary and recurrent tumors, with specific mutation clusters emerging prominently in the recurrent phase. This prominence is further visualized in Panel B, where Cluster 3 showcases an increased VAF from primary to recurrent stages. Panels C and D intricately chart the temporal dominance of these mutation clusters, with Cluster 4 becoming a predominant presence by the time of the recurrent sample collection. The circle packing diagrams in Panels E and F further affirm this observation, spotlighting the clonal expansion in the recurrent tumor. The clonal phylogenetic tree in Panel G elegantly encapsulates the evolutionary trajectory, tracing back to a foundational clone and branching out to diverse evolved clones, reflecting the complexity of clonal evolution in multiple myeloma.
