## Supplementary File 14 for "Bioinformatics Illustrations Decoded by ChatGPT: The Good, The Bad, and The Ugly"

GPT-4

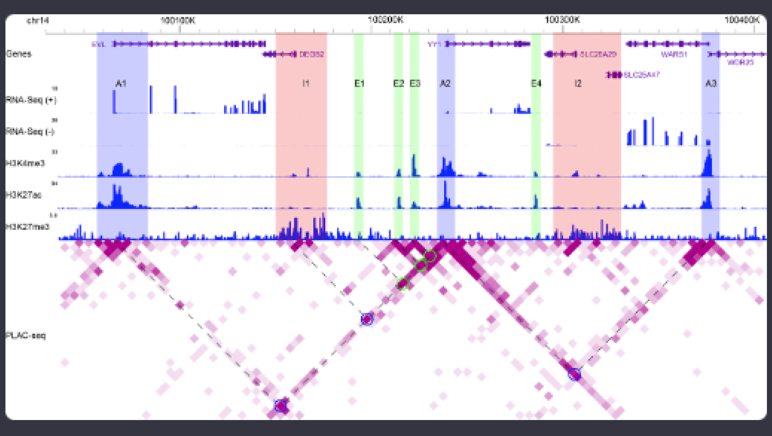

Acting as a bioinformatics professional proficient in RNA-Seq, ChIP-seq, and 3D chromatin analysis, you will help interpret a figure to appreciate transcription regulation at the chromatin level for YY1 in GM12878. The attached figure is a screenshot from the WashU genome browser, encompassing gene annotation, expression data, ChIP-Seq data, and chromatin-chromatin interaction data, for a genomic region enclosing YY1. The aim is to infer transcriptional regulatory mechanisms for YY1 from the epigenomic data. Please do the following.

1. Go over the tracks and explain specific findings addressing the aim.

Certainly, let's break down the figure to understand the transcriptional regulatory mechanisms for the gene YY1 in GM12878 based on the epigenomic data presented.

1. **Gene Track (Genes)**:
   - The gene track shows multiple genes in this genomic region, with YY1 being the main gene of interest.
   - The arrows indicate the direction of transcription. YY1 is transcribed from left to right, meaning it's on the plus strand.
2. **RNA-Seq Tracks (RNA-Seq (+) and RNA-Seq (-))**:
   - These tracks display the RNA sequencing data which reveals gene expression.
   - The plus and minus tracks show the expression levels for genes transcribed on the positive and negative strands respectively.
   - For YY1 (which is on the positive strand), we see distinct peaks, indicating that the YY1 gene is actively transcribed in GM12878.
3. **ChIP-Seq Tracks**:
   - **H3K4me3**: This is a marker for active promoter regions. We observe a strong peak near the start of the YY1 gene, which suggests that this region functions as an active promoter for YY1 in GM12878.
   - **H3K27ac**: This is a marker for active enhancer regions as well as active promoters. We see multiple peaks across the region, indicating potential enhancer sites and further reinforcing the active promoter status of YY1.
   - **H3K27me3**: This is a marker for repressed chromatin. The lack of strong peaks in the vicinity of YY1 suggests that the chromatin is not in a repressed state around this gene.
4. **PLAC-seq Track**:
   - PLAC-seq is a method to capture long-range chromatin interactions.
   - The interaction heatmap reveals regions of the genome that are in close physical proximity, suggesting potential regulatory interactions.
   - Around the YY1 gene, we see some strong interactions (dark pink/red regions). The circled areas indicate particularly significant interactions.
   - These interactions could be between the YY1 promoter and distant enhancer elements. By cross-referencing with the H3K27ac track, we can pinpoint potential enhancer regions that are in physical contact with the YY1 promoter, suggesting a direct regulatory relationship.

**Interpretation**:

- YY1 is actively transcribed in the GM12878 cell line, as indicated by the RNA-Seq data.
- The YY1 promoter region is active, and there are several potential enhancer regions nearby, as shown by the H3K4me3 and H3K27ac peaks.
- The YY1 promoter is interacting with distant regions, some of which may function as enhancers regulating YY1 expression. These interactions can be mediated by protein complexes that bring these distant regions into close physical proximity to enhance YY1 transcription.
- The lack of H3K27me3 suggests that the chromatin environment around YY1 is not repressive in GM12878.

Overall, the data suggests a complex regulatory landscape for YY1 in GM12878, with multiple potential enhancers in interaction with its promoter, ensuring its active transcription in this cell type.

2. Are there any specific suggestions to improve the data presentation of the figure?

Certainly! While the figure provides a wealth of information, there are a few suggestions that could improve its clarity and data presentation:

1. **Color Contrast and Palette**:
   - Ensure there's a good color contrast to make sure each track stands out distinctly. This can help in quickly distinguishing between different data types.
   - Consider using a colorblind-friendly palette for all tracks to make the figure more accessible.
2. **Label Enhancements**:
   - It might be helpful to label the significant peaks directly on the figure with short annotations. For example, label the significant peaks in the H3K27ac and H3K4me3 tracks as potential 'enhancer' or 'promoter', respectively.
   - Enhance the gene labels, especially for YY1, to make it more pronounced than other genes.
3. **Zoomed-in Regions**:
   - For regions with dense interactions or overlapping peaks, consider providing a zoomed-in panel to dissect the intricate details without crowding the main plot.
4. **Scale/Units**:
   - Ensure that each track has a clear scale or unit of measurement (e.g., reads per million for RNA-Seq). This helps in quantitatively comparing the data across different regions.
5. **Legends and Keys**:
   - A clear legend explaining the color intensity in the PLAC-seq heatmap (indicating the strength of interaction) would be beneficial.
   - Add legends or a key for any symbols or special markings, like the circles in the PLAC-seq track.
6. **Interaction Highlight**:
   - Consider using arrows or dashed lines to connect regions in the YY1 gene with the specific distant regions it interacts with on the PLAC-seq track. This can visually guide the reader on which regions are interacting.
7. **Consolidate Tracks**:
   - If certain tracks like RNA-Seq (+) and RNA-Seq (-) can be combined without loss of clarity, it might reduce the overall height of the figure, making it more compact. Alternatively, collapsing tracks that aren't immediately necessary for the main point can also be considered.
8. **Background Shading**:
   - Use alternating light shades for different genomic regions or genes to provide a visual separation between them.
9. **Track Titles**:
   - Make sure each track's title clearly defines what the track represents, possibly including the specific cell line or condition if applicable.
10. **Interactive Features** (for online presentation):
    - If this figure is for an online platform, consider making it interactive. For instance, hovering over a peak could provide additional details, or clicking on an interaction could highlight the corresponding regions on other tracks.

Implementing these suggestions can enhance the figure's comprehensibility, ensuring that the rich data it contains is easily accessible to readers, regardless of their familiarity with the specific field.

3. Draft a title and figure legend. Make sure to include details.

**Title**:
"Epigenomic Landscape and Chromatin Interactions Regulating YY1 Transcription in GM12878 Cells"

**Figure Legend**:
Figure X: Multi-layered epigenomic data elucidating the transcriptional regulatory mechanisms of the YY1 gene in GM12878 cells.

- **Genes**: Genomic annotations displaying the direction of transcription with arrows. The highlighted gene is YY1.
- **RNA-Seq (+) and RNA-Seq (-)**: RNA sequencing tracks represent the expression levels of genes transcribed on the positive and negative strands, respectively. Peaks in the RNA-Seq tracks denote active transcription.
- **H3K4me3**: ChIP-seq data showcasing the trimethylation of histone H3 at lysine 4. Peaks represent active promoter regions.
- **H3K27ac**: ChIP-seq data for histone H3 acetylation at lysine 27, a marker for active enhancer regions as well as active promoters.
- **H3K27me3**: ChIP-seq data indicating the trimethylation of histone H3 at lysine 27. This mark is associated with repressed chromatin. The absence of peaks near YY1 suggests non-repressive chromatin.
- **PLAC-seq**: Chromatin interaction heatmap displaying regions of the genome in close physical proximity, indicating potential regulatory interactions. Darker shades of pink/red signify stronger interactions. Circled regions highlight significant chromatin interactions of interest.

Note: Data derived from the WashU genome browser, focusing on a genomic region encompassing the YY1 gene. This comprehensive view provides insights into the intricate interplay of chromatin modifications and their potential role in regulating YY1 expression in GM12878 cells.

4. Draft a paragraph to describe the findings from the figure. This is one paragraph for the result part of a manuscript. 150 words or less.

In the epigenomic landscape surrounding the YY1 gene in GM12878 cells, our data reveals pronounced transcriptional activity as indicated by prominent RNA-Seq peaks. Concomitantly, the presence of H3K4me3 at the YY1 start site designates an active promoter, while multiple H3K27ac peaks across the genomic stretch signify both an active promoter status and potential enhancer sites. Notably, the absence of H3K27me3 suggests a non-repressed chromatin environment around the YY1 locus. More intriguingly, PLAC-seq data delineates robust chromatin-chromatin interactions involving the YY1 promoter region. These interactions, signified by darker hues on the heatmap, suggest a network of distal regulatory elements potentially orchestrating YY1 transcription. Together, this multi-dimensional epigenomic overview underscores the intricate regulatory mechanisms governing YY1 expression in GM12878, with its promoter engaging in dynamic crosstalk with multiple distant regulatory elements.
