## Supplementary File 15 for "Bioinformatics Illustrations Decoded by ChatGPT: The Good, The Bad, and The Ugly"

GPT-4

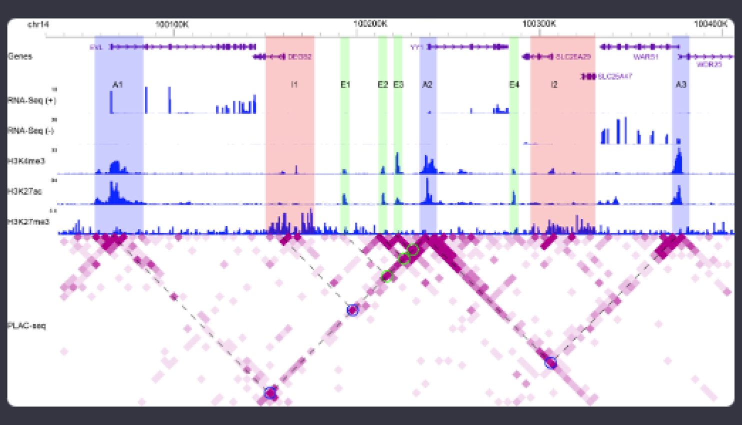


Acting as a bioinformatics professional proficient in RNA-Seq, ChIP-seq, and 3D chromatin analysis, you will help interpret a figure to appreciate transcription regulation at the chromatin level for YY1 in GM12878. The attached figure is a screenshot from the WashU genome browser, encompassing gene annotation, expression data, ChIP-Seq data, and chromatin-chromatin interaction data, for a genomic region enclosing YY1. The aim is to infer transcriptional regulatory mechanisms for YY1 from the epigenomic data. Please do the following.

1. Go over the tracks and explain specific findings addressing the aim.

Certainly! Let's analyze the data step by step:

### 1. Gene Annotation (Genes):

At the top, the figure provides a gene annotation track, illustrating the genomic location and orientation of several genes, including `YY1`. `YY1` is in the forward orientation, as indicated by the arrows.

### 2. Expression Data (RNA-Seq):

Below the gene annotations are two RNA-Seq tracks representing RNA sequencing data on the positive (+) and negative (-) strands. The height of the peaks in the RNA-Seq tracks represents the expression level of the genes.

**Findings:**

- `YY1` appears to be actively transcribed, as indicated by the peaks in the RNA-Seq (+) track directly above its location. Other genes like `DEGS2`, `SLC25A29`, and `WARS1` also show active transcription.

### 3. ChIP-Seq Data for Histone Modifications:

The next three tracks show ChIP-Seq data for three histone modifications: H3K4me3, H3K27ac, and H3K27me3.

**H3K4me3:** This modification is associated with active promoters.
**Findings:** There's a notable peak of H3K4me3 at the beginning of the `YY1` gene, suggesting the presence of an active promoter region there.

**H3K27ac:** This modification indicates active enhancer regions or active promoters.
**Findings:** We see multiple peaks of H3K27ac, especially around the `YY1` gene and between `YY1` and `DEGS2`, suggesting the presence of active enhancers or promoters in these regions.

**H3K27me3:** This mark indicates repressed chromatin.
**Findings:** There's no significant H3K27me3 peak in the vicinity of `YY1`, suggesting that the `YY1` region isn't repressed by this particular histone modification.

### 4. Chromatin-Chromatin Interaction Data (PLAC-seq):

The bottom track illustrates chromatin-chromatin interactions detected by PLAC-seq. Points on this track represent pairs of genomic loci that are in close physical proximity, potentially indicating regulatory relationships.

**Findings:**

- The main diagonal line represents interactions between nearby genomic regions (due to the natural folding of DNA).
- Off the diagonal, we see several highlighted points, indicating specific long-range interactions. The circles highlight potential interactions of interest.
- Notably, there are interactions between regions near the `YY1` promoter (with H3K4me3 and H3K27ac marks) and other regions with H3K27ac marks (indicative of active enhancers). This suggests that the YY1 gene might be regulated by enhancers located at these interacting regions.

### Inference:

From the given epigenomic data, we can infer the following about the transcriptional regulatory mechanisms of `YY1`:

- The `YY1` gene is actively transcribed in GM12878 cells.
- It possesses an active promoter region, as indicated by the H3K4me3 mark.
- Several potential enhancer regions nearby (and possibly some at a distance) could be regulating its transcription, as evidenced by H3K27ac marks and chromatin-chromatin interactions.
- There's no evidence of repression by the H3K27me3 mark in the vicinity of `YY1`.

In conclusion, `YY1` appears to be actively transcribed in GM12878, potentially regulated by multiple enhancer regions both nearby and at a distance, as indicated by the chromatin-chromatin interaction data.

2. Are there any specific suggestions to improve the data presentation of the figure?

Certainly! Effective visualization is critical for the interpretation and communication of complex genomic data. Here are some suggestions to improve the data presentation of the figure:

1. **Contrast & Color Selection:**
   - While the current color scheme is functional, using a more diverse palette can help in distinguishing between different tracks more effectively. However, ensure the palette is colorblind-friendly.
2. **Scale Consistency:**
   - Make sure all tracks that measure similar things (like peak heights in RNA-Seq and ChIP-Seq data) have consistent scales, or at least clearly marked scales, so viewers can compare them directly.
3. **Enhance Interaction Points on PLAC-seq track:**
   - The interaction points on the PLAC-seq track could be made more prominent or color-coded based on interaction strength. This would allow viewers to quickly identify strong versus weak interactions.
4. **Labeling & Annotations:**
   - Provide clear labels for each track. This might seem obvious, but a concise description or title for each track can be incredibly helpful.
   - Offer a legend explaining the color-coding, especially if introducing a new palette.
   - Indicate the direction of gene transcription more clearly. While the arrows are present, making them bolder or more prominent can be helpful.
5. **Zoom-In Views:**
   - For regions of particular interest, like the promoter of `YY1`, a zoomed-in view could be provided in a separate panel to showcase details.
6. **Additional Functional Annotation:**
   - Overlay potential functional annotations, like transcription start sites (TSS), transcription factor binding sites, or DNase hypersensitivity sites, if available. This would give more context to the ChIP-Seq peaks.
7. **Contextual Information:**
   - A brief description or header indicating the cell type (GM12878) and the focus of the analysis (`YY1` transcriptional regulation) would set the context right away.
8. **Interactive Features (for online versions):**
   - If this is being presented on a platform that supports interactivity, tooltips can be added. Hovering over a region might provide additional details, like exact genomic coordinates or interaction strength values.
9. **Reducing Visual Noise:**
   - Some tracks might have regions with minimal or no data. Collapsing or minimizing these areas can help focus attention on the most relevant parts.
10. **Reference Lines:**

- Consider adding vertical reference lines across all tracks at key regions, like the `YY1` start site. This can help in correlating features across tracks.

Implementing these suggestions can make the figure more accessible, interpretable, and valuable to both experts in the field and those less familiar with such data presentations.

3. Draft a title and figure legend. Make sure to include details.

### Title:

**Epigenomic Landscape Surrounding the `YY1` Gene in GM12878 Cells: Insights into Transcriptional Regulation Mechanisms**

### Figure Legend:

**Overview:**
This figure displays a comprehensive epigenomic overview of a genomic region encompassing the `YY1` gene in GM12878 cells. The data is sourced from the WashU genome browser and includes gene annotations, expression profiles, histone modification patterns, and chromatin-chromatin interaction maps.

**Tracks (from top to bottom):**

1. **Genes:** Genomic annotation of genes in the region. The direction of arrows indicates the transcriptional orientation of the genes.
   - Notable genes: `YY1`, `DEGS2`, `SLC25A29`, `WARS1`
2. **RNA-Seq (+) and RNA-Seq (-):** RNA sequencing data representing gene expression on the positive and negative strands. Peak heights are indicative of transcriptional activity.
3. **H3K4me3:** ChIP-Seq data for the H3K4me3 histone modification, a marker for active promoters.
4. **H3K27ac:** ChIP-Seq data for the H3K27ac modification, associated with active enhancer regions and active promoters.
5. **H3K27me3:** ChIP-Seq data for the H3K27me3 modification, indicating regions of repressed chromatin.
6. **PLAC-seq:** Chromatin-chromatin interaction map, revealing physical interactions between different genomic loci. Highlighted points off the diagonal signify specific long-range interactions, with circles marking interactions of particular interest.

**Interpretation:** The presented data offers insights into the transcriptional regulation of the `YY1` gene. Active transcription is observed for `YY1`, and its regulatory landscape includes potential enhancer regions and promoter activity as indicated by the H3K4me3 and H3K27ac marks. The PLAC-seq interactions further suggest distant regulatory elements that may influence `YY1` expression.

4. Draft a paragraph to describe the findings from the figure. This is one paragraph for the result part of a manuscript. 150 words or less.

In our investigation of the transcriptional regulatory mechanisms governing the `YY1` gene in GM12878 cells, we observed distinct epigenomic signatures reflective of its active transcriptional state. RNA-Seq data unequivocally confirmed the gene's expression. A notable peak of the H3K4me3 histone mark, indicative of active promoters, was found at the onset of `YY1`. Furthermore, surrounding regions exhibited prominent H3K27ac peaks, suggesting the presence of active enhancers or additional promoter activities. Intriguingly, these H3K27ac-rich areas participated in notable chromatin-chromatin interactions, as evidenced by our PLAC-seq data. Notably absent were significant H3K27me3 signals, negating the possibility of repression by this histone modification in the vicinity of `YY1`. Cumulatively, our findings underscore the role of a dynamic epigenomic landscape, comprising active promoters, potential enhancers, and long-range chromatin interactions, in modulating the transcriptional activity of `YY1` in GM12878 cells.
