## Supplementary File 16 for "Bioinformatics Illustrations Decoded by ChatGPT: The Good, The Bad, and The Ugly"

GPT-4

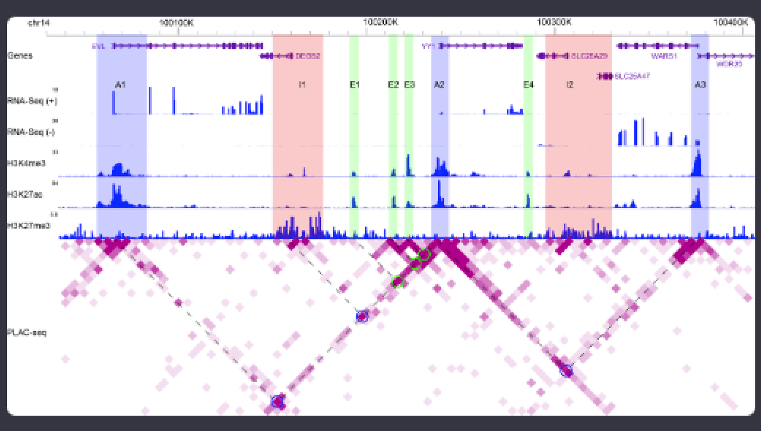


Acting as a bioinformatics professional proficient in RNA-Seq, ChIP-seq, and 3D chromatin analysis, you will help interpret a figure to appreciate transcription regulation at the chromatin level for YY1 in GM12878. The attached figure is a screenshot from the WashU genome browser, encompassing gene annotation, expression data, ChIP-Seq data, and chromatin-chromatin interaction data, for a genomic region enclosing YY1. The aim is to infer transcriptional regulatory mechanisms for YY1 from the epigenomic data. Please do the following.

1. Go over the tracks and explain specific findings addressing the aim.

Sure, let's break down the data tracks presented in this figure:

### 1. Gene Annotation Track:

- The genes in the region are EVL, DEGS2, YY1, SLC25A29, WARS1, SLC25A47, and WDR25.
- Our gene of interest, YY1, is in the middle of this genomic segment and it's transcribed from right to left (denoted by the arrow direction).

### 2. RNA-Seq Tracks:

- There are two RNA-Seq tracks: positive strand (+) and negative strand (-). The peaks on these tracks represent the expression level of genes.
- YY1 shows notable expression peaks in both strands, suggesting it is actively transcribed in GM12878 cells.

### 3. ChIP-Seq Data:

- **H3K4me3 (Tri-methylation of lysine 4 on histone H3)**: This mark is usually associated with active promoters. A distinct peak is observed upstream of the YY1 transcription start site (TSS), suggesting an active promoter region.
- **H3K27ac (Acetylation of lysine 27 on histone H3)**: This mark is found near active regulatory regions (both promoters and enhancers). There are peaks observed in the YY1 region, supporting its active transcription and potential enhancer regions nearby.
- **H3K27me3 (Tri-methylation of lysine 27 on histone H3)**: This mark is associated with transcriptionally repressed regions. There seems to be little to no enrichment of this mark near the YY1 gene, indicating that there isn't significant transcriptional repression in this region for GM12878 cells.

### 4. PLAC-seq (Chromatin-Chromatin Interaction Data):

- The PLAC-seq track displays long-range interactions between different chromatin regions.
- There are interactions (lines) connecting the YY1 region with other distant sites. The highlighted circles signify the strength or frequency of these interactions.
- The areas where YY1 interacts (highlighted in the PLAC-seq track) likely represent regulatory regions like enhancers that might be regulating YY1 transcription, as supported by the presence of the H3K27ac mark.

### Summary:

1. **YY1 is Actively Transcribed**: The RNA-Seq data shows distinct expression peaks for YY1, indicating active transcription in GM12878 cells.
2. **Active Promoter for YY1**: The H3K4me3 peak near the TSS of YY1 suggests the presence of an active promoter.
3. **Potential Enhancer Regions**: The H3K27ac peaks around YY1, and the chromatin-chromatin interactions highlighted in the PLAC-seq data, suggest the presence of potential enhancer regions that might be regulating YY1 transcription.
4. **No Significant Transcriptional Repression**: The lack of a pronounced H3K27me3 peak suggests that the YY1 region isn't transcriptionally repressed in these cells.

In conclusion, the epigenomic data suggests that YY1 is actively transcribed in GM12878 cells, with evidence of both an active promoter and potential enhancer regions that may be involved in its regulation. The chromatin-chromatin interaction data from PLAC-seq provides additional insight into the regulatory landscape, highlighting regions that may be interacting with the YY1 locus to modulate its expression.

2. Are there any specific suggestions to improve the data presentation of the figure?

Certainly! While the provided figure offers a comprehensive view of the genomic landscape surrounding the YY1 gene, there are always ways to improve data visualization for clarity and enhanced interpretation. Here are some suggestions:

1. **Consistent Color Scheme**: Using a consistent color scheme for related data can help in the quick identification of specific tracks. For example, all RNA-Seq data could be in shades of blue, ChIP-Seq data in shades of purple, and interaction data in shades of red.
2. **Scale Adjustment**: Ensure all tracks are on a consistent and relevant scale. If one particular ChIP-seq track has a significantly higher signal than others, consider normalizing or presenting on a log scale to avoid overshadowing other tracks.
3. **Track Labeling**: While the tracks are labeled, providing a brief description or legend for each track might be helpful for those less familiar with the data types. For instance, briefly describing what H3K4me3, H3K27ac, and H3K27me3 represent can aid interpretation.
4. **Highlight Key Regions**: Important regions, like the promoter of YY1 or potential enhancers, can be highlighted or annotated with text labels directly on the figure.
5. **Interactive Features (if feasible)**: If this visualization is meant for an online platform, interactive features like zooming, panning, and tooltips with more information about specific regions or peaks can be extremely helpful.
6. **Simplify Data where Possible**: For a general audience or a high-level presentation, consider simplifying the view by showing only the most relevant tracks. For instance, if the main focus is YY1 transcription, perhaps show only the RNA-Seq data, the H3K4me3, H3K27ac, and the PLAC-seq interaction data.
7. **Grid Lines and Reference Points**: Adding faint grid lines or specific reference points can help in correlating peaks across different tracks.
8. **Improve Resolution of PLAC-seq Data**: The PLAC-seq data is dense, and it might be beneficial to increase the resolution or clarity of interaction points, especially those that are of particular interest.
9. **Order Tracks by Relevance**: Tracks can be ordered by their relevance to the central aim of the study. For instance, if the primary aim is to explore YY1 transcription, then RNA-Seq data can be placed closest to the gene annotation track, followed by ChIP-seq data and then PLAC-seq data.
10. **Unified Peak Highlighting**: If certain peaks or regions in different tracks correspond to the same genomic location, consider using the same color or style to highlight them for easy cross-reference.

Implementing these suggestions can enhance the visual appeal of the figure, streamline data interpretation, and cater to both expert and general audiences.

3. Draft a title and figure legend. Make sure to include details.

Certainly!

---

### Title:

**Epigenomic Landscape and Chromatin Interactions Surrounding the YY1 Gene in GM12878 Cells**

---

### Figure Legend:

**Figure X**: Comprehensive epigenomic profiling of the YY1 gene region in GM12878 cells.

- **Gene Annotation**: This track displays the genomic context, including several genes with YY1 in the center. The direction of arrows indicates the transcription direction.
- **RNA-Seq (+/-)**: Expression levels from RNA sequencing data are shown for both positive (+) and negative (-) strands. Peaks represent active transcription regions. Notably, YY1 shows significant expression in these cells.
- **H3K4me3 ChIP-Seq**: Tri-methylation of lysine 4 on histone H3 is associated with active promoters. A distinct peak upstream of YY1's transcription start site suggests an active promoter region.
- **H3K27ac ChIP-Seq**: Acetylation of lysine 27 on histone H3 indicates active regulatory regions. Peaks around YY1 hint at active transcription and potential enhancer regions.
- **H3K27me3 ChIP-Seq**: Tri-methylation of lysine 27 on histone H3 marks transcriptionally repressed regions. Minimal enrichment near YY1 indicates a lack of significant transcriptional repression.
- **PLAC-seq**: Chromatin-chromatin interaction data unveils long-range interactions between chromatin regions. Interactions connecting YY1 with distant sites could represent enhancer regions or other regulatory elements.

Colored rectangles (e.g., A1, E1, etc.) mark specific regions of interest for more detailed study or reference.

---

When using this title and legend, please ensure that it fits the context and format of the publication or presentation, and adjust as needed.

4. Draft a paragraph to describe the findings from the figure. This is one paragraph for the result part of a manuscript. 150 words or less.

Certainly!

---

In the GM12878 cellular context, the transcriptional regulatory landscape surrounding the YY1 gene was extensively characterized using multi-layered epigenomic data. RNA-Seq analysis confirmed YY1's active transcription, as indicated by prominent expression peaks on both transcriptional strands. The presence of an H3K4me3 peak proximal to the YY1 transcription start site underscores an active promoter, while H3K27ac enrichment suggests regions of transcriptional activation and potential enhancer activity nearby. Notably, the scarcity of the H3K27me3 mark around YY1 indicates minimal transcriptional repression. Importantly, PLAC-seq data revealed long-range chromatin interactions between YY1 and distant genomic regions, hinting at potential distal regulatory elements. Collectively, these data highlight a dynamic chromatin environment around YY1 in GM12878 cells, pointing to multiple regulatory elements that likely govern its transcriptional activity.

---
