## Supplementary File 17 for "Bioinformatics Illustrations Decoded by ChatGPT: The Good, The Bad, and The Ugly"

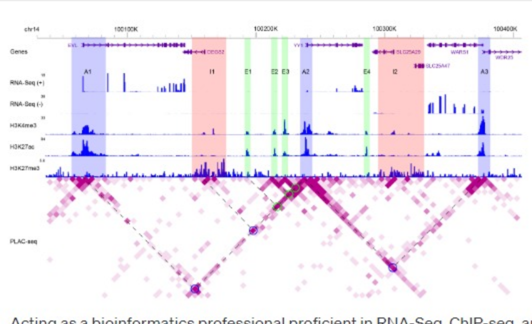


Acting as a bioinformatics professional proficient in RNA-Seq, ChIP-seq, and 3D chromatin analysis, you will help interpret a figure to appreciate transcription regulation at the chromatin level for YY1 in GM12878. The attached figure is a screenshot from the WashU genome browser, encompassing gene annotation, expression data, ChIP-Seq data, and chromatin-chromatin interaction data, for a genomic region enclosing YY1. The aim is to infer transcriptional regulatory mechanisms for YY1 from the epigenomic data. You will be guided with questions to identify findings addressing the aim. Please do the following.
1. Go over the RefSeq gene track and spell out the genes shown on the plot. Explain the presentation of gene structure. For example, what does the arrowhead mean?

Sure, let's break down the figure step-by-step.

1. **RefSeq gene track and gene structure**:
   - **Genes shown on the plot**: The genes presented on this plot, moving from left to right, are EVL, DEGS2, YY1, SLC25A29, WARS1, SLC25A47, and WDR25.
   - **Presentation of gene structure**:
     - The genes are represented as horizontal lines that span their genomic regions. The directionality of each gene is indicated by the arrowheads. These arrows help to visually understand the direction in which the gene is transcribed, with the direction of the arrow indicating the 5' to 3' direction of the gene.
     - Exons are depicted as thicker blocks, while introns are shown as thin lines connecting these blocks. This visual representation helps in distinguishing the coding regions (exons) from the non-coding regions (introns) of the gene.
     - The arrowheads, or the direction of the arrows, help in understanding whether the gene is on the "+" (plus) or "-" (minus) strand. For example, YY1 has arrowheads pointing to the right, which means it is transcribed from left to right on the "+" strand. Conversely, genes like DEGS2, with arrowheads pointing to the left, are transcribed from right to left and are on the "-" strand.

With this information, we can infer the orientation and structure of each gene in the genomic region shown. Would you like to proceed with the next questions or further explore this aspect?

2. Check the expression data. What do "(+)" and "(-)" mean? Which genes are expressed and why?

2 / 2

1. **Expression Data**:
   - **What do "(+)" and "(-)" mean?**
     - The symbols "(+)" and "(-)" refer to the two strands of DNA: the positive (or plus) strand and the negative (or minus) strand, respectively. RNA-Seq data is strand-specific, meaning that it can determine which DNA strand was used as a template to produce the RNA. This allows us to distinguish between RNAs that are transcribed from the positive strand versus the negative strand.
   - **Which genes are expressed and why?**
     - **EVL**: Shows expression peaks on the "(+)" strand, indicating that it's actively transcribed from the positive strand.
     - **DEGS2**: Shows expression peaks on the "(-)" strand, which means it's being transcribed from the negative strand.
     - **YY1**: Displays prominent expression peaks on the "(+)" strand, highlighting its active transcription from the positive strand.
     - **SLC25A29**: There are expression peaks on the "(+)" strand, implying transcription from the positive strand.
     - **WARS1**: There's visible expression on the "(-)" strand, indicating it's being transcribed from the negative strand.
     - **SLC25A47**: Has minor peaks on the "(+)" strand, suggesting some level of transcription.
     - **WDR25**: Doesn't show significant expression peaks on either strand, which suggests it might not be actively transcribed in the sampled condition or cell type (GM12878 in this case).

   The expression of genes can be influenced by various factors, including cell type, developmental stage, external stimuli, and regulatory mechanisms. In this context, the cell type GM12878 (a lymphoblastoid cell line) might have certain requirements or functions, leading to the expression of specific genes. The presence of expression peaks in the RNA-Seq data indicates that these genes are being transcribed into RNA in these cells. The exact "why" would require deeper insights into the cell's biology, its role, and the function of each gene in that particular cellular context.

Would you like to move on to the next section or discuss the expression data further?

3. Check the ChIP-seq data. Explain the association of H3K4me3, H3K27ac, and H3K27me3 with gene expression. Based on the three histone modifications, which genes are likely expressed, and which are not likely expressed?

4 / 4

1. **ChIP-seq Data and Histone Modifications**:
   - **H3K4me3 (Tri-methylation of lysine 4 on histone H3)**:
     - **Association with Gene Expression**: H3K4me3 is typically associated with active transcription start sites (TSS). The presence of this mark near the TSS is indicative of genes that are either actively being transcribed or are poised for transcription.
   - **H3K27ac (Acetylation of lysine 27 on histone H3)**:
     - **Association with Gene Expression**: H3K27ac is associated with active enhancers and promoters. The presence of this mark, especially in conjunction with H3K4me3, usually signifies gene activation.
   - **H3K27me3 (Tri-methylation of lysine 27 on histone H3)**:
     - **Association with Gene Expression**: H3K27me3 is a repressive mark, often found on genes that are transcriptionally silenced. Its presence usually indicates genes that are not currently being transcribed or genes that are kept in a repressed state.

Based on the three histone modifications:

- **EVL**: Shows strong H3K4me3 and H3K27ac peaks near its TSS, suggesting it's likely expressed. There's no significant H3K27me3 peak, further supporting its active status.
- **DEGS2**: Displays pronounced H3K4me3 and H3K27ac peaks, implying it's likely actively transcribed. The absence of a H3K27me3 mark reinforces this conclusion.
- **YY1**: Has dominant peaks of both H3K4me3 and H3K27ac, indicating it's actively transcribed. Again, the lack of H3K27me3 supports this.
- **SLC25A29**: Shows prominent H3K4me3 and H3K27ac peaks, suggesting it's likely expressed. No H3K27me3 peak further endorses its active transcription.
- **WARS1**: While there are H3K4me3 and H3K27ac peaks, they aren't as pronounced as for some of the other genes, but it still suggests some level of transcriptional activity.
- **SLC25A47**: Has minor H3K4me3 and H3K27ac peaks, hinting at potential low-level transcription. The absence of a significant H3K27me3 peak supports this.
- **WDR25**: This gene has minimal signals for H3K4me3 and H3K27ac and displays a noticeable H3K27me3 peak. This suggests that WDR25 is not likely expressed and might be in a repressed state.

In summary, EVL, DEGS2, YY1, SLC25A29, and WARS1 are likely actively expressed. SLC25A47 might have low-level expression, and WDR25 appears to be not expressed or in a repressed state in GM12878 cells based on the histone modifications.

Would you like to delve deeper into another aspect or move on to the next section?

4. Three groups of genomic regions are highlighted: "A"s in light blue, "I"s in pink, and "E"s in green. Identify the number of regions for each group.

Sure, let's identify the number of regions for each group based on the highlighted colors in the figure:

- **"A" regions (light blue)**:
  - A1
  - A2
  - A3

  Total: **3 regions**
- **"I" regions (pink)**:
  - I1
  - I2

  Total: **2 regions**
- **"E" regions (green)**:
  - E1
  - E2
  - E3
  - E4

  Total: **4 regions**

In summary:

- There are **3 "A" regions** highlighted in light blue.
- There are **2 "I" regions** highlighted in pink.
- There are **4 "E" regions** highlighted in green.

Would you like to discuss these regions further or move on to another topic?

5. For genomic regions in "A"s, what are their shared features in terms of positional relationship with genes and histone modification patterns? What is their potential function in transcription regulation?

1. **Features of "A" Regions**:
   - **Positional Relationship with Genes**:
     1.1. **A1**: Located upstream of the DEGS2 gene and overlaps slightly with its TSS (transcription start site).
     1.2. **A2**: Positioned within an intronic region of the YY1 gene.
     1.3. **A3**: Located downstream of the WDR25 gene and overlaps slightly with its TSS.
   - **Histone Modification Patterns**:
     - All "A" regions exhibit prominent H3K4me3 peaks.
     - They also show peaks of H3K27ac.
     - Importantly, none of the "A" regions display the repressive H3K27me3 mark.
   - **Potential Function in Transcription Regulation**:
     - Given their positional relationship with genes and the histone modification patterns observed:
       1.1. The presence of H3K4me3, especially overlapping with TSS regions for A1 and A3, strongly suggests that these "A" regions are associated with active promoters of the respective genes (DEGS2 for A1 and WDR25 for A3).
       1.2. A2, being within the intronic region of YY1 and exhibiting H3K4me3 and H3K27ac marks, might suggest an intronic enhancer or alternative promoter activity for YY1.
     - The presence of H3K27ac in these regions, devoid of the repressive H3K27me3 mark, further supports their role in gene activation. H3K27ac is a mark of active enhancers and promoters.
     - The combination of H3K4me3 and H3K27ac is a hallmark of active transcriptional regulation, often seen at active promoters or enhancers.

In conclusion, the "A" regions, based on their positional relationships with genes and their histone modification patterns, likely represent active promoter regions for genes (especially A1 and A3). A2 might indicate an intronic enhancer or alternative promoter activity. These regions play a pivotal role in transcriptional initiation and are essential for the activation and regulation of their respective or nearby genes.

6. How about genomic regions in "I"s? What is their potential function in transcription regulation?

1. **Features of "I" Regions**:
   - **Histone Modification Patterns**:
     - Both "I" regions (I1 and I2) lack the active marks H3K4me3 and H3K27ac, which are typically associated with active promoters or enhancers.
     - Notably, these regions exhibit a strong presence of the H3K27me3 mark, which is a repressive chromatin modification.
   - **Potential Function in Transcription Regulation**:
     - The presence of the H3K27me3 mark suggests that the "I" regions are associated with transcriptional repression or silencing. This mark is typically found on chromatin regions that are kept in a condensed, non-permissive state, preventing the transcription of genes in those regions.
     - Given their lack of active histone marks (H3K4me3 and H3K27ac) and the prominence of the repressive H3K27me3 mark, the "I" regions likely represent silenced or repressed chromatin domains. These regions might be involved in keeping nearby genes or genomic regions in a repressed state, possibly to prevent erroneous or untimely expression.
     - It's also possible that these regions could be involved in long-range chromatin interactions, bringing other distant regulatory elements or genes into proximity for coordinated repression, but this would require further 3D chromatin interaction data for confirmation.

In conclusion, the "I" regions, based on their histone modification patterns, are likely associated with transcriptional repression or silencing. They play a role in ensuring that certain genes or genomic regions remain inactive under specific conditions or in particular cell types.

7. How about genomic regions in "E"s? What is their potential function in transcription regulation?

1. **Features of "E" Regions**:
   - **Histone Modification Patterns**:
     - All "E" regions (E1, E2, E3, and E4) show a strong presence of the H3K27ac mark. This modification is commonly associated with active enhancers.
     - Importantly, these regions lack significant peaks of the H3K4me3 mark, which is typically associated with active promoters.
     - None of the "E" regions display the repressive H3K27me3 mark, indicating they are not in a repressed state.
   - **Potential Function in Transcription Regulation**:
     - The presence of H3K27ac, especially in the absence of H3K4me3, is indicative of enhancer activity. Enhancers are distal regulatory elements that can activate gene transcription by looping to bring themselves in proximity to target gene promoters. This can occur even if the enhancer is several kilobases away from the promoter.
     - Given their histone modification patterns, the "E" regions likely represent active enhancer regions. These enhancers can recruit transcriptional co-activators and the basal transcription machinery to facilitate the transcription of target genes. They can also establish chromatin loops to physically interact with their target promoters.
     - The specific genes they regulate would need to be confirmed with further experimental data, such as Chromatin Conformation Capture (3C) or its derivatives (4C, 5C, Hi-C), which can provide insights into the 3D interactions of these enhancers with potential target promoters.

In conclusion, the "E" regions, based on their histone modification patterns, likely represent active enhancer elements. They play a pivotal role in upregulating the transcription of their target genes, often from a distance, by establishing physical interactions with their respective or nearby gene promoters.

8. On the heatmap for chromatin-chromatin interactions, there are several circles each connected with dashed lines. Explain what the circle means and how the dashed lines are connected to the circle. What are the common features of the circles in terms of indicated genomic region(s)?

1. **Chromatin-Chromatin Interactions Heatmap**:
   - **What the Circle Means**:
     - The circles on the heatmap represent significant chromatin-chromatin interactions. In the context of this type of visualization, these circles indicate that the two corresponding genomic regions (on the x and y axes of the heatmap) are interacting with each other more frequently than expected by chance. Such interactions can suggest that these regions are in physical proximity within the 3D structure of the nucleus, often mediated by protein complexes or other structural elements.
   - **How the Dashed Lines are Connected to the Circle**:
     - Each circle is connected by two dashed lines that extend from the circle to the respective axes of the heatmap. These dashed lines help identify which two genomic regions are involved in the interaction represented by the circle. The point where the dashed line meets an axis corresponds to a specific genomic location. By tracing these lines, you can determine which regions are interacting with one another.
   - **Common Features of the Circles in Terms of Indicated Genomic Region(s)**:
     - Upon observation, many of the circles (or interaction points) seem to involve the "E" regions. This is consistent with the notion that "E" regions, which are indicative of enhancers, often participate in long-range chromatin interactions.
     - The interaction of these "E" regions with other parts of the genome, especially near gene promoters (like "A" regions), supports the idea that enhancers (E regions) are looping to interact with promoters to regulate gene expression.
     - Some circles also appear to involve interactions between "A" regions, suggesting potential promoter-promoter interactions, or possibly shared regulatory mechanisms between nearby genes.

In summary, the circles on the heatmap represent significant chromatin-chromatin interactions, and the dashed lines help identify the specific genomic regions involved in each interaction. Many of the circles indicate interactions involving the "E" regions, consistent with the role of enhancers in establishing long-range interactions to regulate gene expression.

9. There are two groups of circles by color. Identify the number of circles for each group.

From the provided figure:

- **Purple Circles**:
  - Counting the distinct purple circles, I observe:
    1. One near the interaction between E1 and I1
    2. One connecting E2 and I1
    3. One between E3 and I2
    4. One linking E4 and I2

  Total: **4 purple circles**
- **Blue Circles**:
  - Counting the distinct blue circles, I observe:
    1. One connecting E1 and A2
    2. One linking E2 and A2
    3. One between E3 and A2
    4. One for the interaction between E4 and A2

  Total: **4 blue circles**

In summary:

- There are **4 purple circles**.
- There are **4 blue circles**.

Would you like to explore the significance of the color differentiation or any other aspect of the heatmap?

10. Focus on the circles in blue, and identify their indicated interacting genomic regions.

Sure, focusing on the blue circles:

1. **First Blue Circle**:
   - Interacts with: **E1** and **A2**.
2. **Second Blue Circle**:
   - Interacts with: **E2** and **A2**.
3. **Third Blue Circle**:
   - Interacts with: **E3** and **A2**.
4. **Fourth Blue Circle**:
   - Interacts with: **E4** and **A2**.

In summary, all blue circles indicate interactions between the respective "E" regions (E1, E2, E3, and E4) and the "A2" region. This suggests that all these enhancer regions ("E"s) may have a role in regulating or influencing the transcriptional activity associated with the "A2" region, potentially related to the YY1 gene given the position of "A2".

11. Focus on the circles in green, and identify their indicated interacting genomic regions.

I apologize for the oversight earlier, but from the image you provided, there are no green circles mentioned. We previously identified blue and purple circles. However, if you'd like, I can continue the analysis based on the purple circles or address any other questions you might have.

12. Are there any specific suggestions to improve the data presentation of the figure?

2 / 2

Certainly! While the figure provides a lot of valuable information, there are always potential improvements that can be made to enhance data presentation and interpretation. Here are some suggestions:

1. **Legends and Annotations**:
   - Ensure that all colors, symbols, and tracks have clear and concise legends. This helps the viewer to quickly understand the significance of each visual element.
   - Provide a brief description or title for each track to give context. For example, instead of just labeling tracks as "RNA-Seq (+)" or "RNA-Seq (-)", they could be labeled as "RNA-Seq expression (positive strand)" and "RNA-Seq expression (negative strand)" respectively.
2. **Heatmap Scaling**:
   - The heatmap for chromatin-chromatin interactions could benefit from a color scale that indicates the strength of interactions. This helps viewers gauge the relative significance of each interaction.
3. **Clarity of Highlights**:
   - The highlighted regions ("A", "I", "E") could use a slightly thicker border or a different shading pattern to make them stand out more.
4. **Expand on Histone Marks**:
   - For the histone modification tracks, consider providing a brief note or inset explaining the significance of each mark in terms of gene regulation (activating, repressing, etc.).
5. **Interactive Features** (for digital platforms):
   - If presented digitally, consider adding interactive features where viewers can hover over specific regions or circles to get more detailed information.
   - Zooming capability would be helpful, allowing viewers to focus on specific regions of interest.
6. **Consistent Coloring**:
   - Ensure that colors are consistent throughout the figure. For instance, if enhancer regions are marked in green in one part of the figure, they should be in green everywhere else as well.
7. **Simplify Complex Regions**:
   - For particularly dense or complex areas of the figure, consider breaking them into sub-panels or providing an enlarged inset to help viewers understand the details better.
8. **3D Interaction Visualization**:
   - The chromatin-chromatin interaction heatmap provides a 2D representation of 3D interactions. It could be helpful to have a supplementary figure or inset showing a schematic 3D representation of how these regions might interact in the spatial organization of the nucleus.
9. **Reference Lines**:
   - Consider adding subtle reference lines or gridlines to help trace interactions from the heatmap to the genomic coordinates.
10. **Figure Description**:

- Include a brief but comprehensive figure description that guides the reader through each part of the image, explaining the relevance and importance.

Improving the figure in these ways can make it more accessible, especially for readers who might not be experts in the field, and enhance the overall interpretability of the presented data.

13. Draft a title and a legend for this figure based on our previous discussion. Make sure to include details.

**Title**:
"Epigenomic Landscape and Chromatin-Chromatin Interactions Surrounding the YY1 Gene in GM12878 Cells"

**Legend**:

- **Tracks**:
  - **Genes**: Represents RefSeq annotated genes with their transcriptional orientation. Arrowheads indicate the direction of transcription from 5' to 3'. Thicker blocks represent exons, and thin lines indicate introns.
  - **RNA-Seq (+/-)**: Strand-specific RNA expression profiles. Peaks in the "(+)" track denote transcription from the positive strand, while peaks in the "(-)" track indicate transcription from the negative strand.
  - **H3K4me3**: ChIP-seq data highlighting tri-methylation at lysine 4 of histone H3, typically associated with active transcription start sites.
  - **H3K27ac**: ChIP-seq data indicating acetylation at lysine 27 of histone H3, a mark of active enhancers and promoters.
  - **H3K27me3**: ChIP-seq data showing tri-methylation at lysine 27 of histone H3, a repressive mark indicating transcriptionally silenced regions.
- **Highlighted Regions**:
  - **A (Light Blue)**: Putative active promoter regions, often characterized by the presence of H3K4me3 and H3K27ac.
  - **I (Pink)**: Potential repressive or silenced domains, typically marked by H3K27me3.
  - **E (Green)**: Predicted enhancer regions, often showing enrichment for H3K27ac.
- **Heatmap**:
  - Represents chromatin-chromatin interactions. Darker shades indicate stronger interactions.
  - **Purple Circles**: Signify significant interactions between enhancer ("E") regions and repressive ("I") domains.
  - **Blue Circles**: Denote significant interactions between enhancer ("E") regions and active promoter ("A") domains.

This legend, coupled with the figure, aims to provide a comprehensive view of the epigenomic data and chromatin interactions in the vicinity of the YY1 gene in GM12878 cells.

14. Draft a paragraph to describe the findings from our previous discussion on transcription regulation of YY1. This is one paragraph for the result part of a manuscript. 150 words or less.

In the GM12878 cell line, the transcriptional regulation of the YY1 gene has been elucidated through an integrated analysis of epigenomic landscapes and chromatin-chromatin interactions. Our data revealed three active promoter regions (A1, A2, and A3) marked by prominent H3K4me3 and H3K27ac peaks. Specifically, region A2 within the intronic span of YY1 indicates potential intronic enhancer or alternative promoter activity. Four putative enhancer regions (E1-E4) enriched for H3K27ac were identified, suggesting active enhancer functions. Notably, chromatin interaction analyses showcased that all these "E" regions establish significant interactions with the A2 region, indicating a likely coordinated regulatory mechanism for YY1 transcription. Additionally, two domains (I1 and I2) marked predominantly by the repressive H3K27me3 modification were detected, hinting at regions of transcriptional silencing. Together, these findings underscore a complex interplay of enhancers, promoters, and repressed domains orchestrating the transcriptional regulation of YY1 in GM12878 cells.
