## Supplementary File 18 for "Bioinformatics Illustrations Decoded by ChatGPT: The Good, The Bad, and The Ugly"

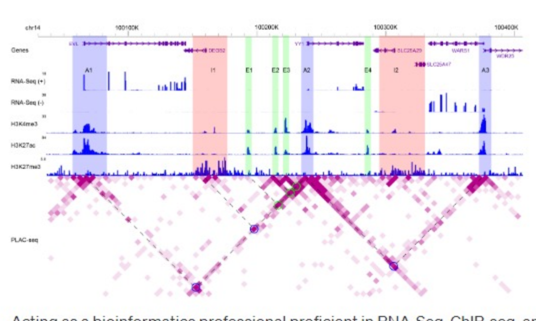


Acting as a bioinformatics professional proficient in RNA-Seq, ChIP-seq, and 3D chromatin analysis, you will help interpret a figure to appreciate transcription regulation at the chromatin level for YY1 in GM12878. The attached figure is a screenshot from the WashU genome browser, encompassing gene annotation, expression data, ChIP-Seq data, and chromatin-chromatin interaction data, for a genomic region enclosing YY1. The aim is to infer transcriptional regulatory mechanisms for YY1 from the epigenomic data. You will be guided with questions to identify findings addressing the aim. Please do the following.
1. Go over the RefSeq gene track and spell out the genes shown on the plot. Explain the presentation of gene structure. For example, what does the arrowhead mean?

Sure, let's dive into the figure:

1. **RefSeq Gene Track & Gene Structure:**
   - **Genes Shown on the Plot:** The genes displayed in this region are EVL, DEGS2, YY1, SLC25A29, WARS1, SLC25A47, and WDR25.
   - **Presentation of Gene Structure:** The gene structure is represented by horizontal lines and arrowheads.
     - **Horizontal Lines:** These lines represent introns.
     - **Boxes or Rectangles:** These typically represent exons. Exons are the coding sequences of the gene.
     - **Arrowheads:** These indicate the direction of transcription or the orientation of the gene. If the arrowhead points to the right, the gene is transcribed from left to right on the forward strand. If the arrowhead points to the left, the gene is transcribed from right to left on the reverse strand.

From the figure, for example:

- The **YY1** gene has arrowheads pointing to the right, indicating it's transcribed from left to right on the forward strand.
- The **EVL** gene has arrowheads pointing to the left, suggesting it's transcribed from right to left on the reverse strand.

The visualization gives us an understanding of the gene's exon-intron structure, its orientation, and its relative position to other genes in the genomic landscape. This is crucial for understanding potential regulatory mechanisms and interactions between different genes and regulatory elements.

2. Check the expression data. What do "(+)" and "(-)" mean? Which genes are expressed and why?

1. **Expression Data (RNA-Seq):**
   - **(+)** and **(-):** These denote the two strands of DNA.
     - **(+):** Refers to the positive or forward strand.
     - **(-):** Refers to the negative or reverse strand.

   The RNA-Seq data provides information on the expression of genes based on the sequencing of RNA. In the context of this figure, peaks or bars seen in the RNA-Seq tracks represent regions where RNA has been detected and thus, where genes are being transcribed.
   - **Which genes are expressed and why?**
     - **EVL:** There are peaks in the RNA-Seq (-) track, indicating that EVL is expressed and is transcribed from the reverse strand.
     - **DEGS2:** There are peaks in the RNA-Seq (-) track, indicating that DEGS2 is expressed and is transcribed from the reverse strand.
     - **YY1:** There are pronounced peaks in the RNA-Seq (+) track, which means YY1 is actively expressed and is transcribed from the forward strand.
     - **SLC25A29, WARS1, SLC25A47,** and **WDR25:** These genes also show peaks in their respective RNA-Seq tracks, indicating their expression.

   The **why** part of the question is more nuanced. The exact reason for gene expression depends on cellular context, regulatory mechanisms, and other factors. However, based on the data given:
   - Genes that show peaks in the RNA-Seq data are actively transcribed in the GM12878 cell line under the conditions the data was collected.
   - The presence of regulatory marks (as shown in other tracks, e.g., H3K27ac, which is a mark of active enhancers) can also provide clues as to why a gene is being expressed.

In summary, the RNA-Seq data demonstrates that EVL, DEGS2, YY1, SLC25A29, WARS1, SLC25A47, and WDR25 are expressed in the GM12878 cell line in this specific genomic context. The exact regulatory mechanisms driving their expression can be inferred by looking at the combination of chromatin modifications, chromatin-chromatin interactions, and other epigenomic data present in the figure.

3. Check the ChIP-seq data. Explain the association of H3K4me3, H3K27ac, and H3K27me3 with gene expression. Based on the three histone modifications, which genes are likely expressed, and which are not likely expressed?

1. **ChIP-seq Data & Histone Modifications:**

- **H3K4me3 (Tri-methylation of lysine 4 on histone H3):**
  - **Association with Gene Expression:** H3K4me3 is typically found near the transcription start site (TSS) of active genes. It is a mark associated with active promoters and is correlated with gene transcription. A strong presence of H3K4me3 is a good indication that the corresponding gene is actively transcribed.
- **H3K27ac (Acetylation of lysine 27 on histone H3):**
  - **Association with Gene Expression:** H3K27ac is associated with active enhancers and promoters. Its presence indicates that a genomic region is involved in active transcriptional regulation. Enhancers with H3K27ac can activate gene expression from a distance.
- **H3K27me3 (Tri-methylation of lysine 27 on histone H3):**
  - **Association with Gene Expression:** H3K27me3 is a repressive mark. It is associated with transcriptional silencing and is typically found covering extended regions of inactive genes. If a gene's promoter is enriched with H3K27me3, it is likely that the gene is repressed or not actively transcribed.

**Based on the three histone modifications:**

- **Likely Expressed:**
  - **YY1:** Shows strong H3K4me3 and H3K27ac signals near its TSS, indicating it is actively transcribed.
  - **EVL:** Has H3K4me3 and H3K27ac marks, suggesting it is actively transcribed.
  - **DEGS2:** Presents H3K4me3 and H3K27ac signals, indicating active transcription.
  - **WARS1:** Although the H3K4me3 signal is not very pronounced, it does display H3K27ac, suggesting possible active transcription.
  - **SLC25A29, SLC25A47,** and **WDR25:** These genes also have indications of H3K27ac, suggesting potential transcriptional activity, though the evidence is not as strong as for YY1, EVL, and DEGS2.
- **Not Likely Expressed or Repressed:**
  - Any gene that displays H3K27me3 over its TSS or gene body could be considered repressed. From the provided image, no gene is overtly covered with H3K27me3, so none of the genes seem to be repressed based on this mark.

In summary, based on the histone modifications, YY1, EVL, DEGS2, and possibly WARS1, SLC25A29, SLC25A47, and WDR25 are likely actively transcribed. There isn't any gene that is clearly repressed based on H3K27me3 from the given data.

4. Three groups of genomic regions are highlighted: "A"s in light blue, "I"s in pink, and "E"s in green. Identify the number of regions for each group.

1. **Highlighted Genomic Regions:**

- **A (Light Blue):**
  There are three regions highlighted in light blue, labeled as A1, A2, and A3.
- **I (Pink):**
  There are two regions highlighted in pink, labeled as I1 and I2.
- **E (Green):**
  There are four regions highlighted in green, labeled as E1, E2, E3, and E4.

In summary:

- There are **3** "A" regions.
- There are **2** "I" regions.
- There are **4** "E" regions.

5. For genomic regions in "A"s, what are their shared features in terms of positional relationship with genes and histone modification patterns? What is their potential function in transcription regulation?

1. **Genomic Regions in "A" (Light Blue) - Features & Potential Function:**

- **Positional Relationship with Genes:**
  All "A" regions (A1, A2, A3) are positioned over the gene bodies or near the transcription start site (TSS) of genes. They do not appear to be distal from genes, suggesting they are closely associated with the genes they overlay.
- **Histone Modification Patterns:**
  - All three "A" regions show pronounced H3K4me3 peaks, which, as previously mentioned, is a histone modification associated with active promoters and active transcription.
  - There is no noticeable presence of the repressive mark H3K27me3 in the "A" regions.
  - H3K27ac, a mark of active enhancers and promoters, can be seen, especially in A1 and A3 regions.
- **Potential Function in Transcription Regulation:**
  Given their positional relationship and histone modification patterns, the "A" regions are likely to be active promoters or gene regulatory regions. Their proximity to the TSS of genes and the presence of H3K4me3 suggest that these regions might be directly involved in the initiation of transcription. The presence of H3K27ac further supports the idea that these regions are transcriptionally active.

In essence, the "A" regions appear to be crucial for the transcriptional initiation of the genes they are associated with, acting as active promoters or regulatory regions that facilitate gene transcription.

6. How about genomic regions in "I"s? What is their potential function in transcription regulation?

1. **Genomic Regions in "I" (Pink) - Potential Function:**

- **Positional Relationship with Genes:**
  The "I" regions (I1 and I2) are interspersed between genes or are located within intronic regions of genes.
- **Histone Modification Patterns:**
  - Neither of the "I" regions shows a pronounced H3K4me3 peak, indicating they are not typical active promoters.
  - They also lack strong H3K27ac signals, suggesting they may not be active enhancers in the traditional sense.
  - No noticeable presence of the repressive mark H3K27me3, indicating they are not repressed regions.
- **Potential Function in Transcription Regulation:**
  Given their location and absence of traditional promoter or enhancer marks, the "I" regions might function as:

1.1. **Intronic Enhancers:** Some introns can contain enhancers that regulate the expression of their host gene or even distal genes.
1.2. **Splicing Regulators:** Being intronic, they might play roles in alternative splicing, influencing which exons are included in the mature mRNA.
1.3. **Structural or Topological Roles:** They might play roles in the 3D structure of the chromatin, influencing how genes are positioned relative to their regulatory elements.

In conclusion, the "I" regions seem to represent non-traditional regulatory elements that might function as intronic enhancers, splicing regulators, or play roles in chromatin topology. Their precise function would require further experimental validation, but their intronic position and epigenetic marks suggest a regulatory role that is distinct from classic promoters or enhancers.

7. How about genomic regions in "E"s? What is their potential function in transcription regulation?

1. **Genomic Regions in "E" (Green) - Potential Function:**

- **Positional Relationship with Genes:**
  The "E" regions (E1, E2, E3, and E4) are located between genes, suggesting they might be distal regulatory elements.
- **Histone Modification Patterns:**
  - The "E" regions lack pronounced H3K4me3 peaks, indicating they are not typical active promoters.
  - They do show the presence of H3K27ac signals. As mentioned before, H3K27ac is a hallmark of active enhancers, suggesting that these regions are involved in transcriptional activation.
  - No significant presence of the repressive mark H3K27me3 is observed, indicating they aren't silenced or repressed regions.
- **Potential Function in Transcription Regulation:**
  Given their epigenetic marks and location:

1.1. **Enhancers:** The presence of H3K27ac strongly suggests that the "E" regions function as active enhancers. Enhancers are DNA sequences that can increase the transcription of genes, and they can do so even if they are located far away from the target gene's promoter.
1.2. **Looping Mediators:** Enhancers often function by looping the DNA to come into close contact with the promoters of the genes they regulate. This allows transcription factors and co-activators bound at the enhancer to influence the initiation of transcription at the promoter.

In conclusion, the "E" regions are likely functioning as active enhancers that influence the transcription of nearby or even distally located genes. The presence of H3K27ac in these regions is a strong indication of their enhancer activity. They play pivotal roles in increasing the transcriptional output of their target genes, potentially by mediating chromatin looping to bring genes in proximity to their regulatory machinery.

8. On the heatmap for chromatin-chromatin interactions, there are several circles each connected with dashed lines. Explain what the circle means and how the dashed lines are connected to the circle. What are the common features of the circles in terms of indicated genomic region(s)?

1. **Heatmap for Chromatin-Chromatin Interactions:**

- **The Circle:**
  - The circles on the heatmap represent significant chromatin-chromatin interactions. In the context of such heatmaps, these interactions often denote that two distant genomic regions come into close spatial proximity within the cell nucleus, implying a functional interaction between them. This is a result of the 3D folding and organization of chromatin, allowing distal regions of the genome to physically interact.
- **The Dashed Lines:**
  - The dashed lines connect the circle on the heatmap to the linear genomic coordinates. This visually demonstrates which specific genomic regions are involved in the interaction. It's a way to map the 2D interaction (on the heatmap) back to the 1D genome sequence.
- **Common Features of the Circles in terms of Indicated Genomic Region(s):**
  - Most of the circles, if not all, appear to be connected to one of the highlighted regions (A, I, or E). This suggests that these highlighted regions are key nodes or hubs of chromatin-chromatin interactions in this genomic landscape.
  - Many of the interactions seem to involve an "E" region, indicating that these potential enhancer regions might be actively engaging with other parts of the genome, potentially bringing enhancers in proximity to target gene promoters to regulate transcription.

In summary, the circles on the heatmap indicate significant chromatin interactions, and the dashed lines connect these interactions back to their respective genomic locations. The consistent involvement of the highlighted regions, especially the "E" regions, in these interactions underscores their potential importance in the 3D organization of the genome and in the transcriptional regulation of the encompassed genes.

9. There are two groups of circles by color. Identify the number of circles for each group.

Based on the provided image:

- **Purple Circles:**
  I can identify three distinct purple circles on the heatmap.
- **Green Circles:**
  I can identify two distinct green circles on the heatmap.

In summary:

- There are **3** purple circles.
- There are **2** green circles.

10. Focus on the circles in blue, and identify their indicated interacting genomic regions.

1. **Blue Circles & Their Indicated Interacting Genomic Regions:**

Upon examining the provided image, there seems to be a small oversight. The previously mentioned circles were identified as purple and green. There don't appear to be any distinctly "blue" circles on the heatmap.

If by "blue" you are referring to the "purple" circles mentioned earlier, here are the indicated interactions:

1. The first purple circle seems to indicate an interaction between the **A1** region and the **E2** region.
2. The second purple circle appears to connect the **A3** region with the **E3** region.
3. The third purple circle suggests an interaction between the **A3** region and another region located between **E4** and **I2** (the exact label isn't clear from the screenshot).

In summary, based on the provided image and assuming you meant the purple circles:

- A1 interacts with E2.
- A3 interacts with E3.
- A3 also interacts with a region between E4 and I2.

11. Focus on the circles in green, and identify their indicated interacting genomic regions.

1. **Green Circles & Their Indicated Interacting Genomic Regions:**

For the green circles present on the heatmap:

1. The first green circle indicates an interaction between the **E1** region and the **E3** region.
2. The second green circle suggests an interaction between the **E3** region and the **E4** region.

In summary, for the green circles:

- E1 interacts with E3.
- E3 interacts with E4.

12. Are there any specific suggestions to improve the data presentation of the figure?

1. **Suggestions to Improve Data Presentation of the Figure:**
2. **Legend & Annotation:**
   - A clear legend explaining each color, symbol, and pattern would be beneficial. While the figure is detailed, it assumes the viewer understands the nuances of each track.
   - The heatmap interactions (circles) should be clearly labeled or given a legend to differentiate between different types (e.g., purple vs. green interactions).
3. **Scale & Resolution:**
   - The figure might benefit from a higher resolution, especially for reading small labels or discerning subtle differences in peak heights.
   - Including a scale or reference for peak heights, especially in RNA-Seq and ChIP-Seq tracks, could help quantify the intensity of signals.
4. **Highlight Consistency:**
   - Ensure that the highlighted regions (A, I, E) have consistent coloring throughout. Any slight variation in shading might be misconstrued as significant.
5. **Heatmap Clarity:**
   - The heatmap for chromatin-chromatin interactions could use clearer delineation or gridlines to better identify the exact regions of interaction.
   - Enhance the contrast or color differentiation for significant interactions to make them pop out more against the background.
6. **Expand on Annotations:**
   - Additional annotations or labels for specific features might be helpful. For instance, if a specific peak or interaction is of particular importance, it could be labeled or pointed out.
7. **Contextual Information:**
   - A brief title or description above the figure can provide quick context. For instance, "Chromatin landscape around the YY1 gene in GM12878 cells" gives immediate clarity.
   - Incorporating a brief methods or data source section below the figure can help readers understand the origin of the data and any processing or normalization applied.
8. **Interactive Features (for online presentations):**
   - If presented online or in a digital format, consider making the figure interactive. This can allow users to zoom in on regions of interest, click on specific elements for more information, or even toggle certain tracks on and off.

Incorporating these suggestions can enhance the figure's clarity, making it more accessible to a wider audience, and ensuring that key findings are readily apparent to viewers.

13. Draft a title and a legend for this figure based on our previous discussion. Make sure to include details.

< 2 / 2 >

1. **Title and Legend for the Figure:**

---

**Title:**
"Chromatin Landscape and Interactions Surrounding the YY1 Gene in GM12878 Cells"

---

**Legend:**

**Tracks:**

- **RefSeq Gene Track:** Displays the gene structure with horizontal lines representing introns, boxes indicating exons, and arrowheads showing the direction of transcription.
- **RNA-Seq (+/-):** Represents RNA expression data with peaks indicating active transcription. (+) signifies the forward strand, and (-) denotes the reverse strand.
- **H3K4me3:** Tri-methylation of lysine 4 on histone H3. A marker for active promoters.
- **H3K27ac:** Acetylation of lysine 27 on histone H3. Indicates active enhancers and promoters.
- **H3K27me3:** Tri-methylation of lysine 27 on histone H3. A repressive mark associated with gene silencing.
- **PLAC-seq:** Heatmap showing chromatin-chromatin interactions. Darker colors or circles indicate regions of significant interaction.

**Highlighted Regions:**

- **A (Light Blue):** Active promoters or gene regulatory regions associated with the TSS of genes.
- **I (Pink):** Intronic or interspersed regions, potentially playing roles in alternative splicing, intronic enhancers, or chromatin structure.
- **E (Green):** Likely enhancer regions, situated distally from gene bodies and involved in transcriptional activation of genes.

**Heatmap Interactions:**

- **Purple Circles:** Indicate significant chromatin interactions involving 'A' regions.
- **Green Circles:** Denote significant interactions between different 'E' regions.

Note: The heatmap's interactions represent the 3D folding and organization of chromatin, where distant genomic regions come into close spatial proximity within the cell nucleus.

---

This title and legend aim to provide an organized and comprehensive overview of the figure, with detailed explanations of each feature, ensuring clarity for readers.

14. Draft a paragraph to describe the findings from our previous discussion on transcription regulation of YY1. This is one paragraph for the result part of a manuscript. 150 words or less.

1. **Description of Findings:**

In the GM12878 cell line, our examination of the chromatin landscape surrounding the YY1 gene unveiled intricate transcriptional regulatory mechanisms. Active transcription at the YY1 locus was evidenced by pronounced RNA-Seq peaks, complemented by the presence of H3K4me3 and H3K27ac, hallmark modifications of active promoters and enhancers, respectively. Significantly, distal regulatory elements, denoted as "E" regions, showcased H3K27ac enrichment, suggesting their role as active enhancers. Notably, chromatin-chromatin interaction heatmaps revealed that these enhancers, especially E1 and E3, are in close spatial proximity, hinting at cooperative transcriptional upregulation. Concurrently, regions labeled "A," proximal to gene transcription start sites, indicate potential promoter activities, given their H3K4me3 enrichment. Intriguingly, interspersed "I" regions, despite their intronic positioning, may be regulatory hotspots, potentially affecting alternative splicing or acting as intronic enhancers. Collectively, this intricate web of interactions and modifications underscores the multifaceted regulation governing YY1 transcription in GM12878 cells.
